# Deep learning-based image-analysis identifies a DAT-negative subpopulation of dopaminergic neurons in the lateral Substantia nigra

**DOI:** 10.1101/2022.12.14.520432

**Authors:** Nicole Burkert, Shoumik Roy, Max Häusler, Dominik Wuttke, Sonja Müller, Johanna Wiemer, Helene Hollmann, Marvin Oldrati, Jorge Ramirez-Franco, Julia Benkert, Michael Fauler, Johanna Duda, Jean-Marc Goaillard, Christina Pötschke, Moritz Münchmeyer, Rosanna Parlato, Birgit Liss

**Affiliations:** Institute of Applied Physiology, Medical Faculty, Ulm University, 89081 Ulm, Germany; Wolution GmbH & Co. KG, 82152 Munich, Germany; UMR_S 1072, Aix Marseille Université, INSERM, Faculté de Médecine Secteur Nord, Marseille, France; INT, Aix Marseille Université, CNRS, Campus Santé Timone, Marseille, FRANCE; Institute of General Physiology, Medical Faculty, Ulm University, 89081 Ulm, Germany; University of Wisconsin-Madison, Dept. of Physics, Madison, WI, US; Linacre College & New College, Oxford University, OX1 4 BH Oxford, UK

## Abstract

Here we present a deep learning-based image analysis platform (DLAP), tailored to autonomously quantify cell numbers, and fluorescence signals within cellular compartments, derived from RNAscope or immunohistochemistry. We utilized DLAP to analyse subtypes of tyrosine hydroxylase (TH)-positive dopaminergic midbrain neurons in mouse and human brain sections. These neurons modulate complex behaviour, and are differentially affected in Parkinson’s and other diseases. DLAP allows the analysis of large cell numbers, and facilitates the identification of small cellular subpopulations. Specifically, we identified a small subpopulation of TH-positive neurons (∼5%), mainly located in the very lateral Substantia nigra (SN), that was immunofluorescence-negative for the plasmalemma dopamine transporter (DAT), with ∼40% smaller cell bodies. These neurons were negative for aldehyde dehydrogenase 1A1, with a lower co-expression rate for dopamine-D2-autoreceptors, but a ∼7-fold higher likelihood of calbindin-d28k co-expression (∼70%). Our results have important implications, as DAT is crucial for dopamine-signalling, and is commonly used as a marker for dopaminergic SN neurons.

## Introduction

Dopamine releasing (DA) neurons within the midbrain are important for a variety of brain functions and behavioral processes like voluntary movement control, learning, cognition, and motivation (Berke, 2018; Björklund & Dunnett, 2007; Liu & Kaeser, 2019; Wise, 2004). The cell bodies of DA midbrain neurons are predominantly located in two overlapping nuclei, the Substantia nigra pars compacta (SN) and the ventral tegmental area (VTA), with axonal projections to the dorsal striatum, or the ventral striatum and prefrontal cortex, respectively (Gantz et al, 2018; Haber & Fudge, 1997; Halliday et al, 2012; Roeper, 2013). In line with these differential functions and projections, dysfunction of the dopaminergic midbrain system can lead to severe diseases like Parkinson’s, Schizophrenia, addiction, or attention deficit hyperactivity disorders, ADHD (Chen et al, 2021; Klein et al, 2019; Lerner et al, 2015; Montero et al, 2021). Importantly, not all DA midbrain neurons are affected equally in these diseases. More precisely, in Parkinson’s disease (PD), the second most common neurodegenerative disorder, a subpopulation of mesostriatal SN DA neurons is progressively degenerating, while neighboring VTA DA neurons remain largely intact (Bloem et al, 2021; Damier et al, 1999; Hirsch et al, 1997; Surmeier et al, 2017). In contrast, in Schizophrenia, mesocorticolimbic VTA DA neurons display complex dysfunctions, while SN DA neurons are mainly unaffected (Jauhar et al, 2022; Kegeles et al, 2010; Owen et al, 2016). The cause for this differential vulnerability of dopaminergic midbrain neurons is still unclear, and only unspecific, symptomatic therapies are available, with their respective side-effects (Smeland et al, 2020; Vijiaratnam et al, 2021).

One prerequisite for curative and cell-type specific therapies is a better understanding of the distinct subpopulations of dopaminergic neurons and their selective pathophysiology (Gantz et al, 2018; Roeper, 2013). Analysis of mRNA and protein expression with single cell resolution and its correlation with anatomical locations and projections in health and disease states is an essential approach for identifying molecular determinants of differential neuronal vulnerability (Dumrongprechachan et al, 2021; Glock et al, 2021; Hobson et al, 2022a; Hobson et al, 2022b; Paget-Blanc et al, 2022). For this, labelling of mRNA or proteins in tissue sections, followed by microscopic imaging, are important tools (Jonkman et al, 2020; Lichtman & Conchello, 2005; Reilly & Obara, 2021). However, manual neuronal mapping, cell counting, and quantification of mRNA- and protein-derived fluorescence signals is very time consuming and prone to human error (Maier-Hein et al, 2018; Schmitz et al, 2011; Sommer & Gerlich, 2013). Here we present a deep learning-based image analysis platform (DLAP) that overcomes these issues. Our single cell DLAP approach and six individual algorithms were tailored to quantify autonomously (i) the number of distinct cell-types in defined areas, as well as (ii) fluorescence signals, derived from either RNAscope probes (mRNA) or immunohistochemistry (proteins), in defined compartments - more precisely in plasma-membranes, cell body/cytoplasm, and nuclei of individual neurons. We utilized artificial neuronal networks (ANN) based artificial intelligence/deep learning approaches for image analysis. ANN are biologically inspired computer programs, designed to simulate the way in which the human brain processes information. ANN gather their knowledge by detecting the patterns and relationships in data (segmentation), and learn (or are trained) through experience, not by programming (Agatonovic-Kustrin & Beresford, 2000; Hassabis et al, 2017; LeCun et al, 2015; Yang & Wang, 2020). ANN consist of artificial neurons, arranged in different layers (Angermueller et al, 2016; Greener et al, 2022), where each consecutive layer obtains inputs from its preceding layer. As only the first (input) and the last (output) layer are visible/accessible, all layers in between are referred to as hidden layers. Network architectures containing hundreds of hidden layers are called deep networks (LeCun et al, 2015). After suitable training, deep learning networks can effectively extract relevant cellular features for automated image analysis (Falk et al, 2019).

To facilitate easy adaption to the most common respective tasks in life science, we have predesigned six algorithms, based on two distinct network structures, DeepLab3 (DL3) and fully convolutional neural networks (FCN) (Chen et al, 2018; Long et al, 2015). We trained them to detect, count, and analyse individual DA midbrain neurons, labelled for tyrosine hydroxylase (TH), the key-enzyme for synthesis of dopamine and other catecholamines (Daubner et al, 2011). However, the flexible design of the algorithms and the workflow allows quick and easy adaptions to other cell types and specimens, and thus is of general interest. Here we detail the DLAP pipeline, including all six predesigned algorithms for automated image processing. As proof-of principle, we demonstrate its reliable identification and count of cells in defined areas, as well as fluorescence-signal quantification in cellular compartments, by analyzing about 40.000 TH-positive midbrain neurons. Using this approach for quantification of the immunofluorescence signal of the plasmalemma dopamine transporter (DAT) in TH-positive SN neurons, we identified a small subpopulation of DAT immuno-negative neurons (∼5%), mainly located in the caudo-lateral parts of the SN (∼37% of all lateral TH-positive SN neurons). These neurons had ∼40% smaller cell bodies and also showed a co-expression profile for three additional markers of subpopulations of SN DA neurons (dopamine-D2-autoreceptor, Ca^2+^ binding protein calbindin-d28k, and aldehyde dehydrogenase 1) that is rather untypical for classical SN DA neurons. The DAT is an electrogenic symporter of the Na^+^/Cl^−^ dependent transporter family (SLC6). It is crucial for re-uptake of dopamine at somatodendrites and axons, and also mediates an electric conductance (Ryan et al, 2021). The identification of a DAT-negative TH-positive SN neuron population has not only physiological relevance (Bu et al, 2021; Miller et al, 2021; Savchenko et al, 2022), but it has additional implications, as DAT is commonly used as marker for SN DA neurons and for their specific targeting, e.g. to generate transgenic mice, expressing Cre-recombinase under the control of the DAT-promoter (Lammel et al, 2015; Papathanou et al, 2019; Stuber et al, 2015).

## Results

Based on the Wolution web-interfaces (https://wolution.ai/), we developed a deep learning based, automated image analysis platform (DLAP) as well as six tailored algorithms (two based on FCN, four based on DL3) for fast and reliable identification and quantitative analysis of TH-positive midbrain neurons within the SN and VTA of mouse and human *post mortem* brain sections. These six DLAP algorithms were trained by hundreds of manually marked TH- and DAPI-positive cell bodies of midbrain neurons. **Figure 1** illustrates the general workflow. **Table 1, 2 and S1** provide details of algorithms, trainings, and performances (further details in methods and supplement). For all approaches, the region of interest (ROI) for cell type identification and quantification (i.e the SN) was marked manually, according to anatomical landmarks (Paxinos & Keith B. J. Franklin, 2007). All six algorithms allowed reliable identification of target cells, as well as their quantification in a given ROI (DLAP-3 & 4), and/or quantification of mRNA-derived (DLAP-1 &-2) or protein-derived (DLAP-5 & 6) fluorescence signals of distinct genes of interest in these target cells - all common tasks in life science.

**Figure 1:**
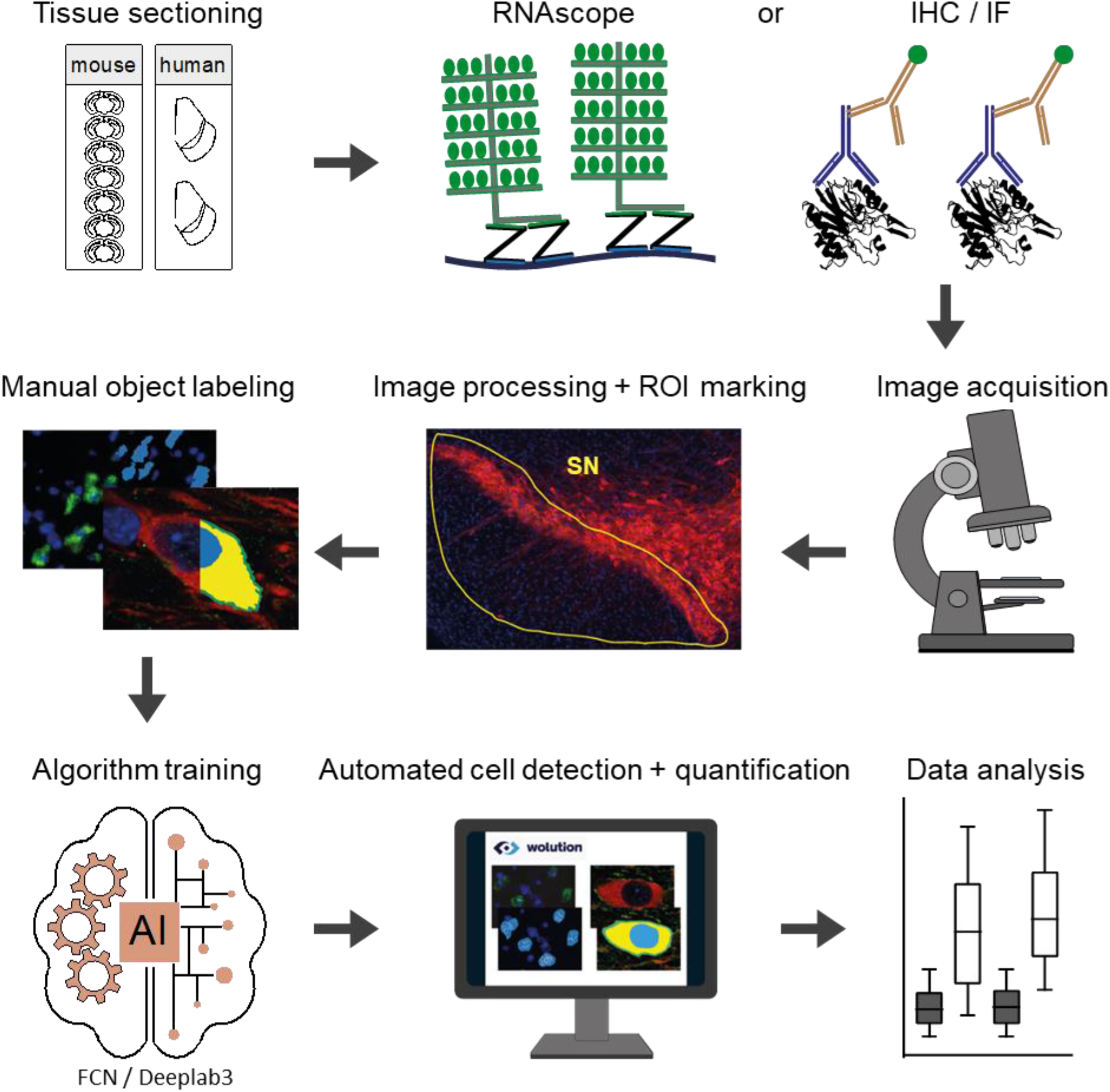
Graphic illustration of the deep learning-based image analysis platform (DLAP). After sectioning of murine/human midbrain samples, either mRNA-(RNAscope) or protein-(IHC/IF) labelling is performed. After image acquisition, images are preprocessed and the ROIs (e.g. SN) are manually marked. Cell types and compartments (e.g. cell body, plasma membrane, nucleus) were manually labeled in a training dataset to generate the ground truth data that was used to train the individual algorithms (FCN- or Deeplab3-based, depending on complexity of signals to identify). After sufficient training, images are automatically analysed (detection of ROI, quantification of cell numbers and signal intensities).

**Table 1:** Quality evaluation parameters for all six algorithms. Performances were determined by comparing DLAP analysis results of the test image sets to the ground truth data (provided by manual labeling). Class name defines the identified molecular or cellular structures. The following quality measures were assessed separately for each class: pixel error rate, rate for true positive (TP), true negative (TN), false positive (FP), false negative (FN) detection, as well as relative specificity (spec.) [defined as TN/(TN+FP)], and sensitivity (sens.) [defined as TP/(TP+FN)].

| DLAP algorithm | class name | Error [%] | TP [%] | TN [%] | FP [%] | FN [%] | Spec [%] | Sens [%] |
| --- | --- | --- | --- | --- | --- | --- | --- | --- |
| 1 | cell body | 5.08 | 10.3 | 84.7 | 4.1 | 0.8 | 95.3 | 92.7 |
|  | mRNA |  | 0.3 | 98.9 | 0.7 | 0.0 | 99.3 | 92.5 |
| 2<br>(human) | cell body | 4.13 | 5.2 | 91.0 | 2.5 | 1.4 | 97.3 | 80.8 |
|  | mRNA |  | 0.2 | 98.9 | 0.9 | 0.0 | 99.1 | 94.7 |
| 3 | nucleus | 0.61 | 0.5 | 98.9 | 0.5 | 0.1 | 99.5 | 85.5 |
| 4 | nucleus | 0.26 | 0.1 | 99.7 | 0.2 | 0.0 | 99.8 | 59.8 |
| 5 | nucleus | 3.3 | 0.9 | 98.5 | 0.3 | 0.3 | 99.7 | 74.5 |
|  | cytoplasm |  | 4.1 | 92.9 | 2.2 | 0.8 | 97.7 | 83.7 |
| 6 | nucleus | 5.49 | 3.0 | 93.5 | 2.5 | 1.0 | 97.4 | 75.6 |
|  | cytoplasm |  | 9.7 | 89.0 | 0.8 | 0.5 | 99.1 | 95.0 |
|  | membrane |  | 11.7 | 85.2 | 2.2 | 0.9 | 97.5 | 92.4 |

**Table 2:** Overview of six individual DLAP algorithms and workflow, optimized for distinct image sets and analysis tasks. For detailed description, see methods. Abbreviations: BF brightfield, DAB diaminobenzidine, DAPI 4′,6-diamidino-2-phenylindole, FCN fully convolutional neural network, HE hematoxylin, IF immunofluorescence, NM neuromelanin, RGB red-green-blue, TH tyrosine hydroxylase. * For training, only the cellular nuclei were marked, not the full cell body, similar as in stereology; only clearly visible, full nuclei were counted.

| DLAP algorithm | Trained by / optimized for | Algorithm task | Network-Type | Training-set |  | post training image analysis optimization (a-e) |
| --- | --- | --- | --- | --- | --- | --- |
|  |  |  |  | images | cells |  |
| **1** | RNAscope fluorescence: TH-mouse (DAPI) | 1) Identifies cells of interest and delineates the shape of cell bodies by TH-derived fluorescence signals (green channel).<br>2) Within these marked cell bodies, target gene derived individual fluorescent dots (red channel) are detected, marked, and counted. | FCN | 100 | ~1000 | (a) watershed algorithm for improved separation of mRNA molecules.<br>(b) morphological operations (type dilation and erosion) applied, for better separation of neurons. |
| **2**<br>modification of algorithm 1 | RNAscope fluorescence: TH-human (DAPI) | 3) A fourth BF channel was included, as a measure for NM-content. After Step (1) and (2) from algorithm 1, in an additional step, the BF signal intensity is quantified within the marked cell bodies (including nuclei). | FCN | 200 | ~500* | (a) watershed algorithm for improved separation of mRNA molecules.<br>(b) morphological operations (type dilation & erosion) applied, for better separation of neurons. |
| **3** | DAB-labelling: TH-mouse | Identifies and counts nuclei of DAB-positive cells (TH-stained). | DeepLab3* | 100 | ~3000* | none |
| **4**<br>modification of algorithm 3 | DAB-labelling: TH-mouse & HE-counterstain | Extends the automated counting of DAB-positive cells (TH-stained) to sections that are counterstained with HE. | DeepLab3* | 300 | ~2500* | (a) watershed algorithm for improved separation of individual nuclei.<br>(c) filling holes within recognized compartments. |
| **5** | IF-labelling: TH-mouse, DAPI<br>DAT-mouse<br>D2/CB/Aldh1A1-mouse | 1) Identifies and counts cells of interest (i.e TH-positive cells). It delineates the shape of the cell body by immunofluorescence signals for TH (red channel) and of the nucleus by DAPI fluorescence signal (blue channel).<br>2) Within these marked compartments, the relative signal intensities are quantified for all four fluorescent channels. | DeepLab3 | 95 | ~500 | (a) watershed algorithm for improved separation of neurons.<br>(c) filling holes within recognized compartments.<br>(d) removing target gene channel during cell identification.<br>(e) RGB intensity threshold for background-signal determination. |
| **6** | IF-labelling: TH-mouse, DAPI<br>Kv4.3-mouse, | 1) Identifies cells of interest and delineates the shape of the cell body by immunofluorescence signals for TH (red channel), of the nucleus by DAPI fluorescence signal (blue channel), and of the membrane (originally learned by Kv4.3 staining, after training no membrane marker needed).<br>2) Within these marked compartments, the relative signal intensities are quantified for all three fluorescent channels. | DeepLab3 | 94 | ~110 | (c) filling holes within recognized compartments.<br>(d) removing target gene channel during cell identification.<br>(e) RGB intensity threshold for background-signal determination. |

Deep learning networks for image segmentation of complicated scenes, such as the Pascal Visual Object Classes (VOC) dataset (Everingham et al, 2015), that we used in the DeepLab3 based algorithms, are usually very large, and their filters are particularly trained to learn abstract concepts such as “tree” or “car”. Particularly, for the analysis of simple, RNAscope derived fluorescence signals (i.e. individual small dots, diameter Ø ∼ 0.3 µm), we found that such a complicated pre-trained neural network was not of benefit. RNAscope-derived fluorescence signals are mainly small dots (Ø ∼ 0.3 µm) within the cytoplasm, which makes threshold based automatic recognition/segmentation of the cell body difficult. Thus, we custom-designed and optimized two less complex FCN-based algorithms (DLAP-1 and 2), and we demonstrate that they were better suited for the analysis of RNAscope data, compared to DeepLab3 approaches. The latter algorithm failed to recognize the small dots (mRNA molecules) as evident from the sensitivity-value of 0.0% for this class, compared to 92.5% and 94.7% for the FCN-based algorithm (**Table S1**). For protein-derived immunohistochemistry signals (diaminobenzidine, DAB and immunofluorescence IF), tailored DeepLab3 algorithms performed better, as indicated by lower overall error (**Table S1**; DLAP-3 to -6).

### DLAP based automated quantification of mRNA molecules in individual target cells via RNAscope approaches

RNAscope *in situ* hybridization allows absolute quantification of distinct mRNA molecules, directly via hybridization-probes, with each fluorescence dot representing one target mRNA-molecule (Jolly et al, 2019; Wang et al, 2012). Due to multiple probes for each target-gene, it is hardly affected by degraded RNA. We have used RNAscope for determining mRNA molecule numbers in individual TH-positive SN DA neurons in fresh-frozen, PFA-postfixed mouse midbrain sections (Benkert et al, 2019). For automated quantification of numbers of RNAscope probe derived fluorescence dots, two custom-designed FCN based algorithms were used (DLAP-1 & 2). The general principle of the FCN algorithms is given in **Figure 2**. DLAP analysis of RNAscope signals was performed in two steps: in a first step, the cell body was recognized and marked according to the TH-signals, and in a second step, individual target gene mRNA dots (labelled with a different fluorophore) were identified and counted within the marked TH-positive cell body, including the nucleus. DLAP-1 is optimized for detecting TH-positive DA neurons from mice, DLAP-2 for detecting human TH-positive DA neurons, containing neuromelanin (NM), a dark pigment present in DA neurons from humans and primates (Bazelon et al, 1967; Vila, 2019), visible only in bright field (BF).

**Figure 2:**
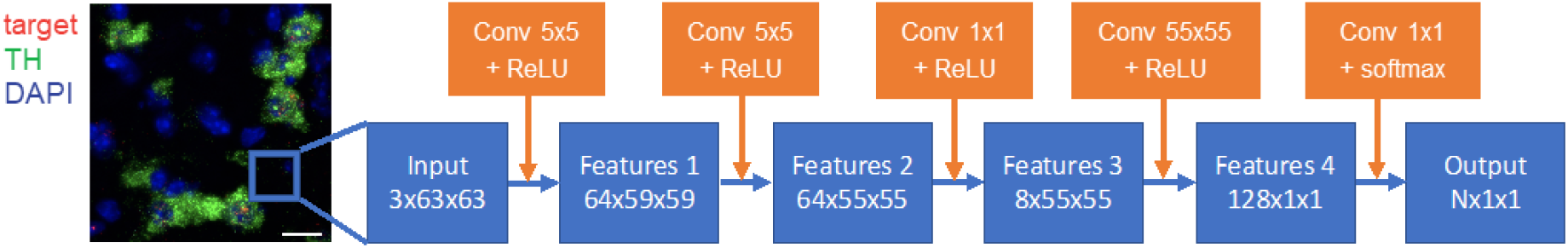
Architecture of the fully convolutional neural network (FCN), used for RNAscope-derived signals. Left: the input image shows SN DA neurons, RNAscope-labelled for tyrosine hydroxylase (TH) mRNA (green), target gene mRNA (red) and DAPI (blue). Scale bar: 20 µm. Right: the image is passed through five convolutional layers, connected by the Rectified Linear Units (ReLU) and the softmax activation function, as indicated. The latter normalises the network output to the probability distribution. The size of the convolutional kernels is specified in the orange boxes, the dimensions of the resulting feature maps is given in the blue boxes [pixels]. The network analyzes image patches of size 63×63 pixels to determine the class of the central pixel in the patch. It is applied convolutionally to every central pixel in the image, so that the final output image has the same dimensions as the input image.

**Figure 3a** shows a representative image of TH-positive SN neurons in a coronal mouse brain section, after RNAscope *in situ* hybridization, before (input image, left) and after automated identification of TH-positive cell bodies and quantification of individual fluorescence dots, derived from target mRNA molecules. Cell bodies were identified and marked according to the TH-derived fluorescence signal (green dots), and target mRNA were identified as probe-derived fluorescence-signals (red dots, **Table S2**). Results for two different target genes - with lower (Cav1.3) and higher (Cav2.3) mRNA abundance - are given (**Figure 3b, Table S3, S4**). Both target genes code for a specific voltage gated Ca^2+^ channel α-subunit that has been linked to Parkinson’s disease (Benkert et al, 2019; Liss & Surmeier, 2022; Ortner, 2021). The manual and the automated analysis (DLAP-1) resulted in very similar results for both genes, over the full range of mRNA molecule numbers (Cav1.3 manual: 24±1 mRNA molecules, DLAP-1: 24±1, N=3, n=499 neurons analysed, p=0.999, R^2^=0.92; Cav2.3 manual: 69±5, DLAP-1: 69±5; N=3, n=372; p=0.999, R^2^=0.92; **Figure 3b/c, S1, Table S3, S4**). Accordingly, the significant, ∼3-fold higher number of Cav2.3 mRNA molecules in mouse SN DA neurons, compared to those of Cav1.3 is robustly detected with both manual and DLAP-1 analysis (p<0.0001 for both), similar as previously described (Benkert et al, 2019).

**Figure 3:**
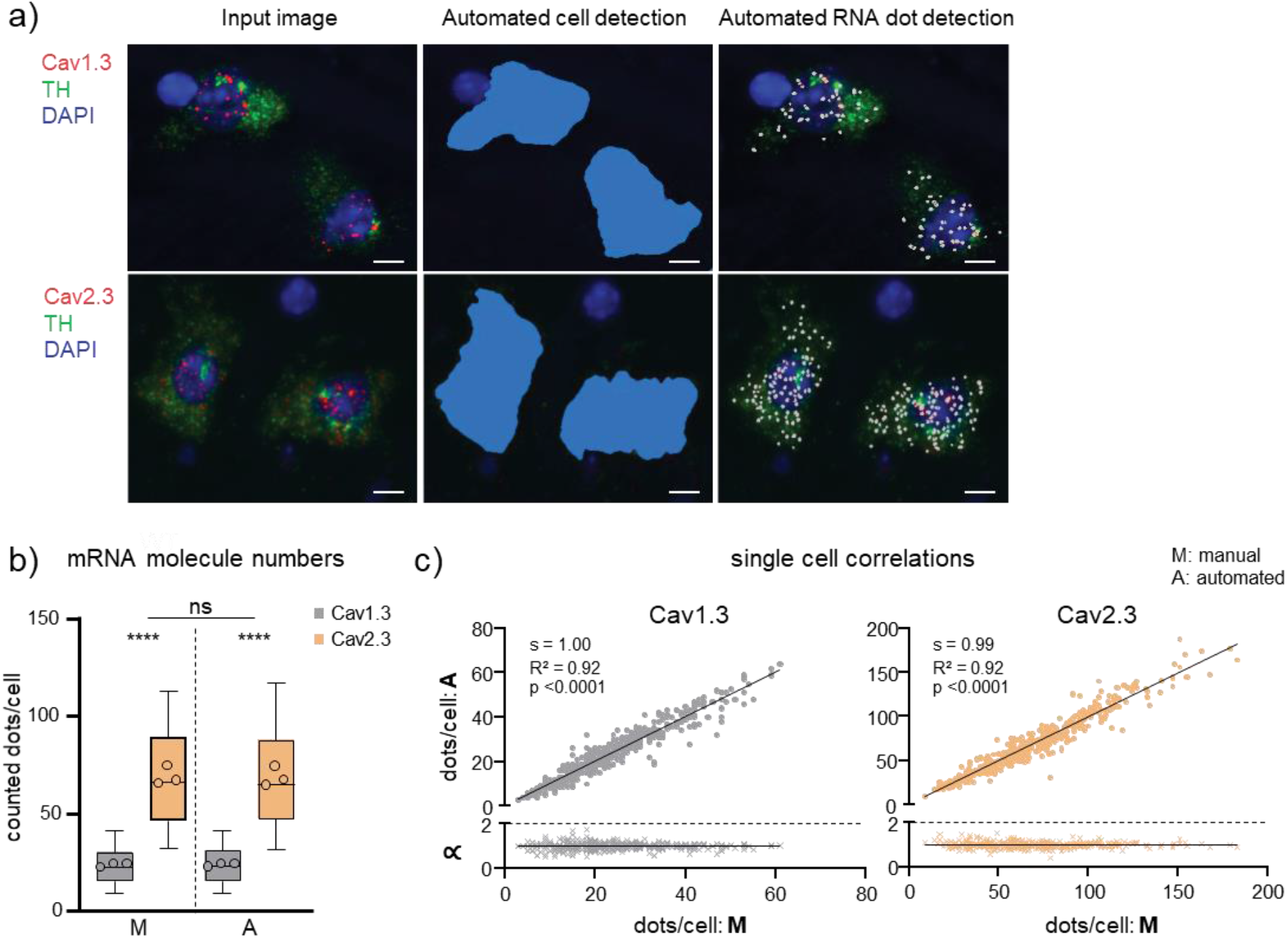
Comparison between manual and automated (DLAP-1) RNAscope-derived image analysis of mouse midbrain sections. a) SN DA neurons in coronal midbrain section from adult mice after RNAscope for tyrosine hydroxylase (TH, green) and for the voltage gated Ca^2+^ channel α-subunits Cav1.3 or Cav2.3 (red). Nuclei are marked by DAPI-staining (blue). Left: input images. Middle and right: images after automated recognition of the TH labelled cell bodies (middle, blue area) and of target mRNA molecules (right, white dots) by DLAP-1. Scale bars: 5 μm. b) mRNA molecule numbers/ neuron, counted manually (M) or automatically (A) via DLAP-1, as indicated. Data are given as boxplots (median, 10-90 percentile) for all analysed neurons (Cav1.3: n = 449, Cav2.3: n = 372). The open circles indicate mean values for each analysed animal (N = 3 each). Significant differences (for analysed neurons, n) according to two-way ANOVA with Tukey’s multiple comparison tests are indicated by asterisks (****: p < 0.0001, ns: p > 0.05). c) Upper: single neuron correlations of mRNA-derived dot counts/cell between manual and automated analysis according to Pearson correlation test (s = slope). Lower: corresponding proportionality constants ∝, calculated from the manual and automated dot count ratios. All data are detailed in Table S3, S4 and Figure S1.

Similar robust results were obtained for RNAscope analysis of human SN DA neurons with the DLAP-2 algorithm. To optimize DLAP for human SN DA neurons, the DLAP-1 algorithm was extended to quantify the brightfield (BF) intensity, as a measure for the variable NM content of human SN DA neurons, to evaluate possible confounding effects of different NM content on RNAscope results (**Figure 4, S2, Table 2**). Cav1.3 mRNA molecules as well as NM content were compared in SN DA neurons from adult and aged individuals (adult: 42±7 years; aged: 78±3, details in **Table S5**). Manual and automated DLAP-2 analysis provided very similar results, and we detected a strong linear correlation between manually and automatically determined mRNA molecule numbers as well as NM-content (**Figure 4b/d/e, Table S6, S7**). Interestingly, the number of Cav1.3 mRNA molecules was significantly lower (∼60%) in SN DA neurons from aged individuals (**Figure 4b left**, manual: adult 38±5, aged 14±3; N=3, n=223; p<0.0001; DLAP-2: adult 40±7, aged 17±4; N=3, n=258; p<0.0001).

**Figure 4:**
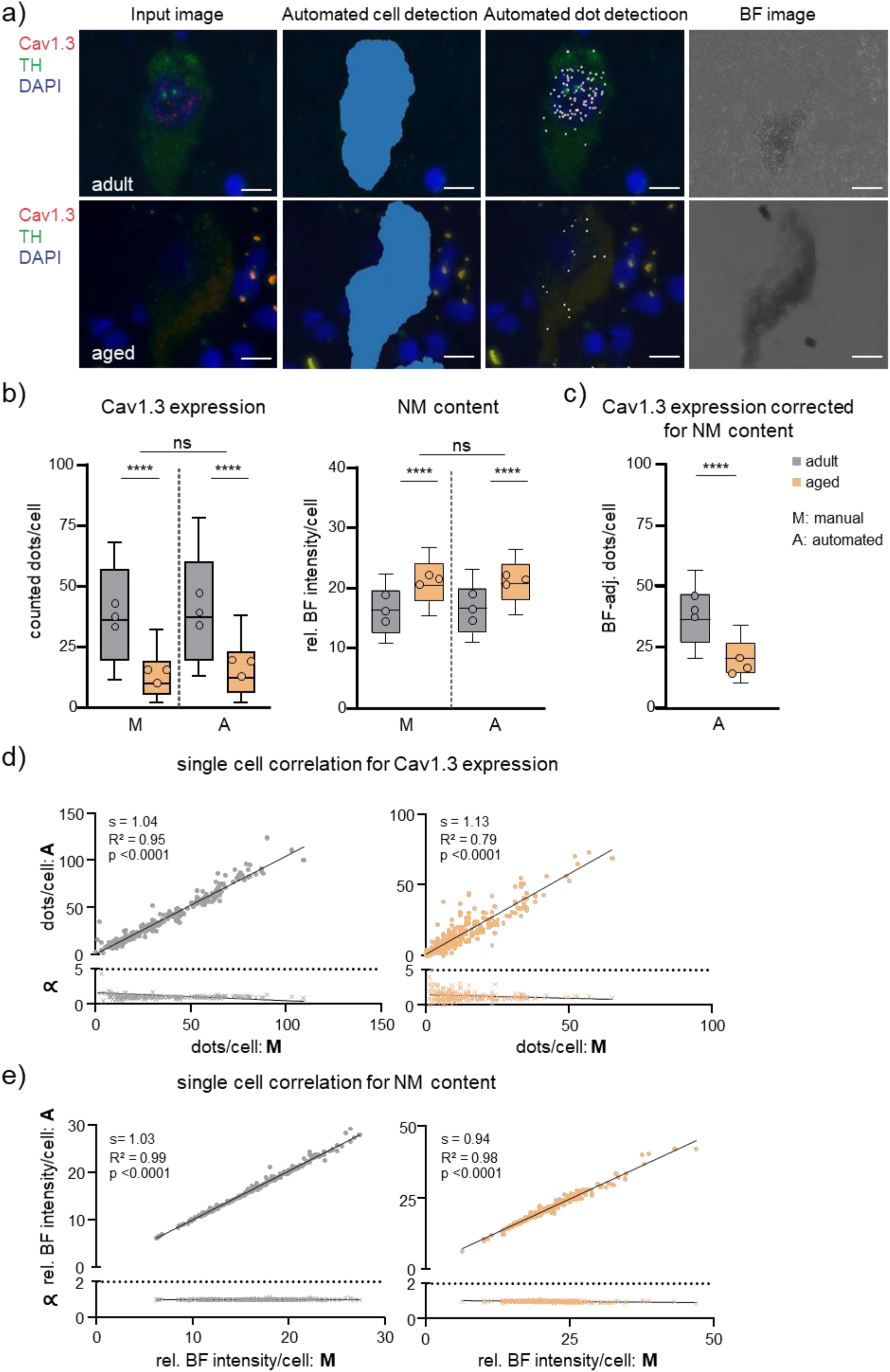
Comparison between manual and automated (DLAP-2) RNAscope-derived image analysis of human midbrain sections. a) SN DA neurons in a horizontal midbrain section from human adult and aged *post mortem* brain samples after RNAscope for tyrosine hydroxylase (TH, green) and for the voltage gated Ca^2+^ channel α-subunit Cav1.3 (red). Nuclei are marked by DAPI-staining (blue). Left: input images. Middle: images after automated recognition of the TH labelled cell bodies (middle left, blue area) and of target mRNA molecules (middle right, white dots) with DLAP-2 Right: corresponding brightfield (BF) images used to quantify neuromelanin (NM) content. Relative BF intensities were measured within the detected cell bodies. Scale bars: 10 μm. b) mRNA molecule numbers/neuron (left) and NM content (rel. BF signal intensity; right), quantified manually (M) or automatically (A) via DLAP-2, as indicated. Data are given as boxplots (median, 10-90 percentile) for all analysed neuros (adult: n = 223, aged: n = 258). The open circles indicate mean values for each analysed brain sample (N = 3 each). Significant differences (for analysed neurons, n) according to two-way ANOVA with Tukey’s multiple comparison tests are indicated by asterisks (****: p < 0.0001, ns: p > 0.05). c) mRNA molecule numbers/SN DA neuron, determined via DLAP-2, adjusted for individual NM using a Mixed Effects Model. Data are given as boxplots (median, 10-90 percentile) for all analysed neuros (adult: n = 206, aged: n = 231). The open circles indicate mean values for each analysed brain sample (N = 3 each). Significant differences (for analysed neurons) according to Mann-Whitney tests (****: p < 0.0001). d/e) Upper: single neuron correlations of Cav1.3 mRNA-derived dot counts/cell (d) and rel. BF signal intensity/cell (e) between manual and automated analysis according to Pearson correlation test. Lower: corresponding proportionality constants ∝, calculated from the manual and automated dot count ratios. All data are detailed in Table S5-7 and Figure S2.

Noteworthy, the BF-value derived NM content was also significantly different between adult and aged samples, with ∼30% higher NM content in aged SN DA neurons (p<0.0001, **Figure 4b right, Table S6, S7**). To exclude that the lower number of mRNA molecules determined in aged SN DA neurons were artificially caused by the higher BF values (as measure for NM content), we addressed possible confounding parameters by applying a mixed effects model optimized for these kind of human sample-derived data that are in general more heterogeneous (Duda et al, 2018; Schlaudraff et al, 2014). As the model indicated that the NM content indeed affected the detected Cav1.3 mRNA molecule numbers, we corrected the automated data for the NM-content attributed effect. However, as evident in **Figure 4c, Table S6**, we still detected a similar highly significant lower number of Cav1.3 mRNA molecules (∼35%) in SN DA neurons from aged donors (manual: adult 32±2, aged 17±3; N=3, n=180; p<0.0001; DLAP-2: adult 32±3, aged 20±3; N=3, n=244; p<0.0001).

### DLAP based automated quantification of immuno-labeled neuron numbers in tissue sections

Determination of numbers of cellular populations in defined ROI is a common task e.g. for quantifying selective cell loss in neurodegenerative diseases, and to assess neuroprotective therapeutic effects (Golub et al, 2015). However, commonly used stereological approaches (still the gold-standard), are time consuming, and they provide only a - more or less exact - estimation of cell numbers, depending on the size of the manually counted fraction of cells (Brown, 2017; West, 1999). Approaches have been developed to increase sampling sizes and thus accuracy of estimates (Mouton et al, 2017). Nevertheless, estimates are prone to over- or underestimating total cell numbers, particularly if the target cells are not homogeneously distributed in the ROI, as it is the case e.g. due to differential neuronal vulnerability in degenerative disease (Baddeley, 2001; Miles & Davy, 1976; Noori & Fornal, 2011). However, manual approaches to count all neurons in a given ROI are extremely time consuming and prone to bias/human error. To overcome these issues, we developed three DLAP algorithms, dramatically reducing the time for valid determination of cell numbers, tailored to detect and count target cells, immuno-labelled by either DAB - without (DLAP-3) or in the presence of a hematoxylin counterstain (DLAP-4) - or by IF (DLAP-5). According to our stereological procedures, the DLAP-3 and 4 algorithms were trained by marking the neuronal nuclei only, not the whole cell bodies, to count only TH-positive cells with clearly visible nuclei.

DAB-labelling is commonly used for determining cell-numbers in histological samples, as stained sections can be archived and e.g. (re-) analysed. **Figure 5, S3, Table S8, S11** summarize the results for counts of TH-positive SN neurons (unilaterally) from juvenile mice after DAB immunostaining, comparing stereological estimates with automated counting, utilizing DLAP-3. The stereological estimates and automated neuron counts resulted in very similar neuron numbers (Stereology: 4951±1079, DLAP-3: 4317±912 neurons unilaterally; p=0.218, N=10. linear correlation/section: R^2^=0.79). These SN DA neuron numbers correspond well to the literature for C57bl/6 mice, ranging from 4000-6000 neurons unilaterally (Brichta & Greengard, 2014; Nelson et al, 1996).

**Figure 5:**
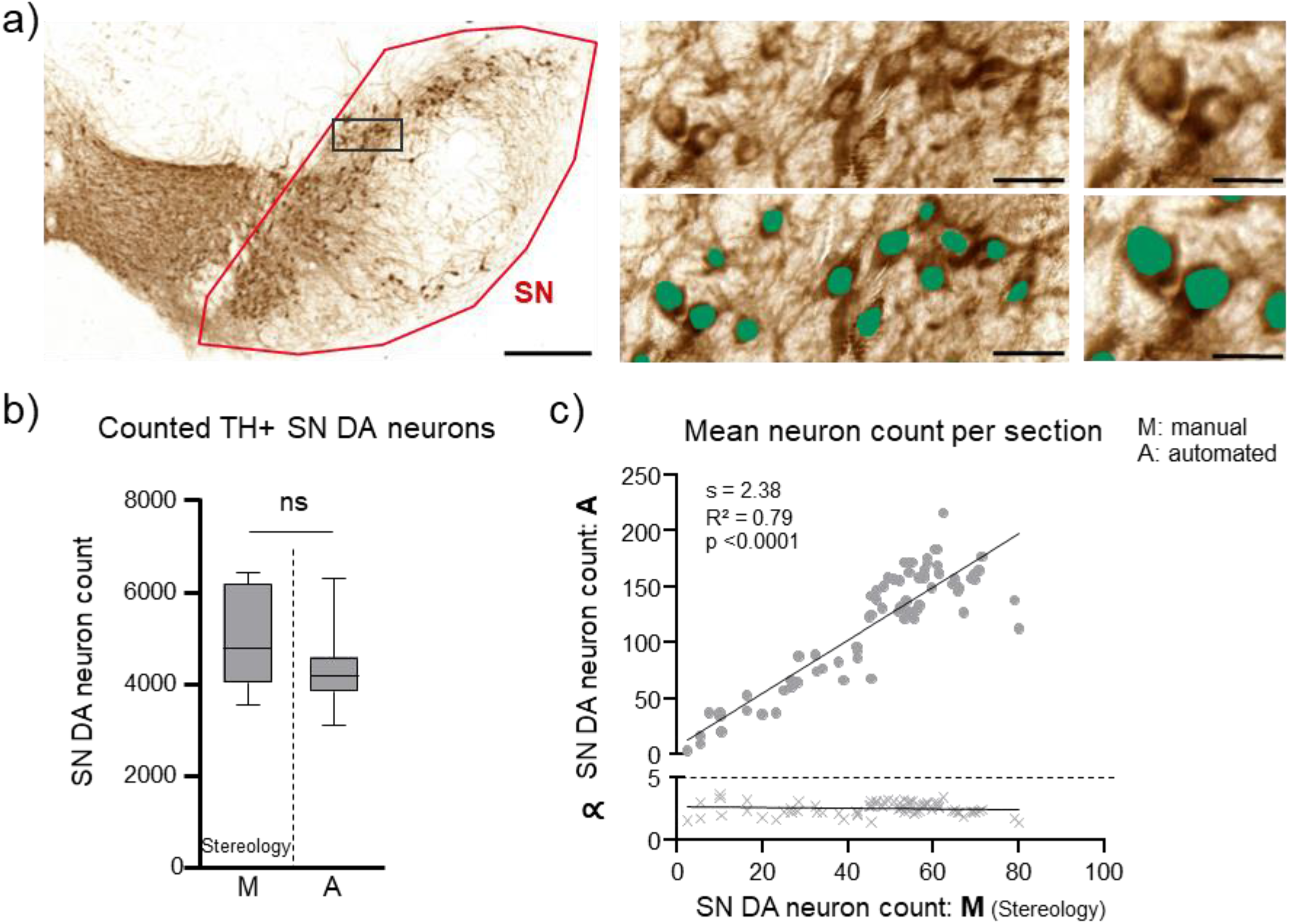
Comparison between unbiased stereology and automated (DLAP-3) IHC-derived image analysis of juvenile mouse midbrain sections. a) SN DA neurons in coronal midbrain section from juvenile mice after DAB-IHC for tyrosine hydroxylase (TH, brown). Left: input image with SN outlined in red. Scale bar: 200 µm. Middle and right: enlarged images before (top) and after (bottom) automated recognition of the nuclei within TH labelled cell bodies (green area) with DLAP-2 Scale bars: 50 μm (middle) and 10 µm (right). b) Numbers of TH-positive SN DA neurons/mouse, quantified via unbiased stereology (extrapolated via the optical fractionator method, M) or automatically (A) via DLAP-3, as indicated. Data are given as boxplots (median, 10-90 percentile) for all analysed mice (N = 10). No significant differences according to Mann-Whitney test (ns: p > 0.05). c) Upper: correlations of SN DA neuron counts/section between stereology and automated analysis, according to Pearson correlation test. Lower: corresponding proportionality constants ∝, calculated from the manual and automated cell count ratios. All data are detailed in Table S8, S11 and Figure S3.

Immunohistochemistry sections are often counterstained with an unspecific cellular marker (like Nissl, hematoxylin). For analysis of SN DA neurons, such counterstains are often performed to confirm that a loss of TH-immunopositive neurons (e.g. in a PD-model) indeed corresponds to a loss of the respective neurons, and does not reflect a mere downregulation of TH-expression (Alam et al, 2017; Healy-Stoffel et al, 2012; Rausch et al, 2022). As we use hematoxylin-counterstains in such experiments (Benkert et al, 2019; Liss et al, 2005), we optimized DLAP-4 to detect DAB stained TH-positive SN neurons in such counterstained sections. With DLAP-4, we re-analysed a cohort of 21 mice that we had already analysed (Benkert et al, 2019) to quantify the effect of a PD drug (1-Methyl-4-phenyl-1,2,3,6-tetrahydropyridin, MPTP) that induces preferential loss of SN DA neurons. Adult C57bl/6J mice were either injected with saline (N=9) or with MPTP/probenecid (N=12). We compared DLAP-4 analysed data with the respective already published stereology-derived data, and also with another automated neuron count approach, based on the Aiforia platform (**Figure 6, S4, Table S9-11**). We had used the Aiforia platform (https://www.aiforia.com) for this cohort, to confirm relative numbers of remaining SN DA neurons after drug-treatment. Indeed, the relative determined cell loss by the Aiforia approach was similar to that estimated by stereology (∼40%, p=0.183; **Figure 6b, Table S10**, and Benkert et al (2019)). However, the absolute number of neurons determined with the Aiforia approach was almost double as high (saline: stereology 3959±643, Aiforia 7048±843, p<0.0001, drug: stereology 2504±933, Aiforia 4108±1596, p=0.004; **Figure 6b, S4, Table S9**). This overcount, and the fact that the convolutional neuronal network (CNN)-based Aiforia system does not allow algorithm training by the users (Penttinen et al, 2018), let us switch to the Wolution-platform in the first instance.

**Figure 6:**
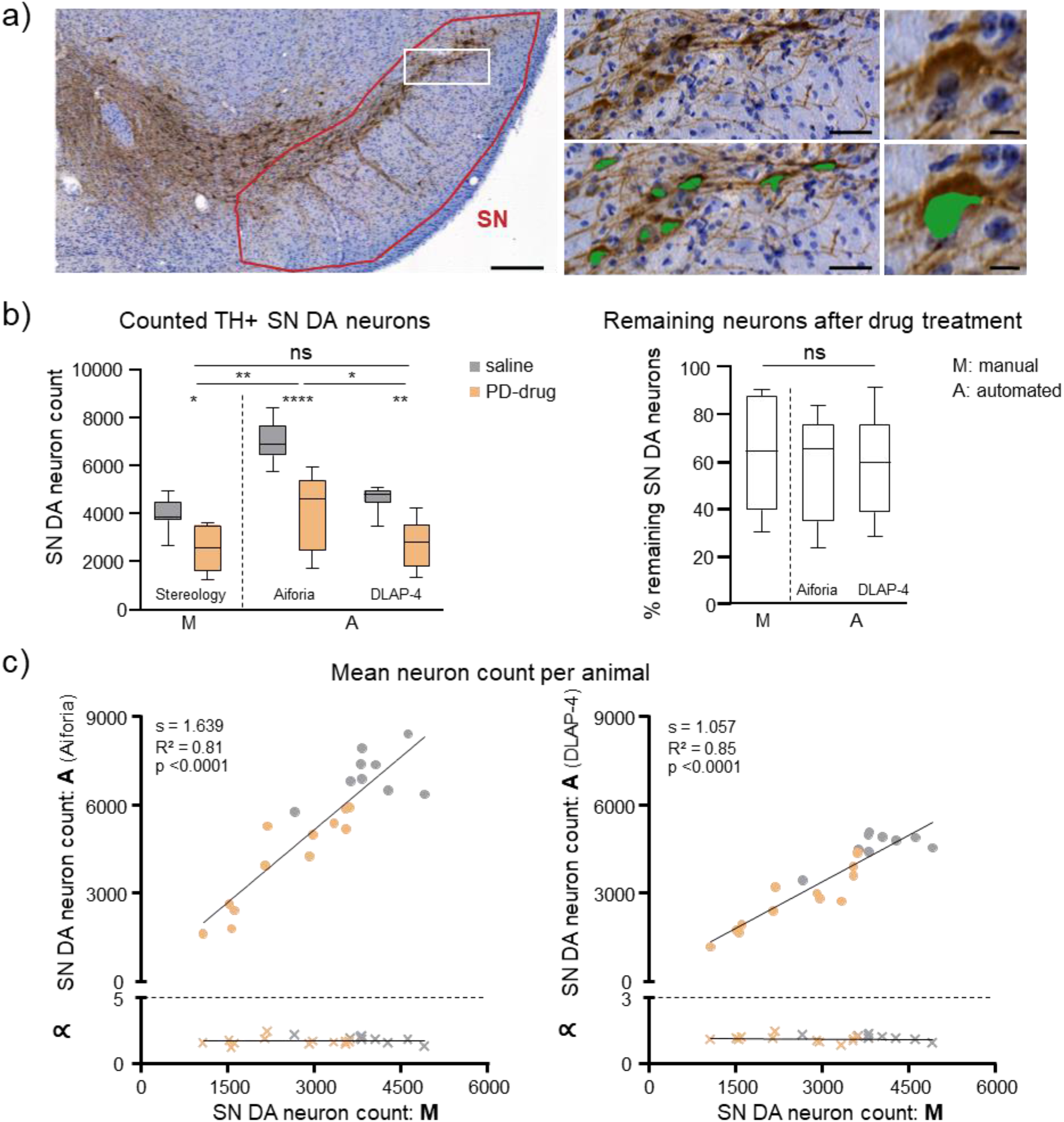
Comparison between unbiased stereology and automated (DLAP-4) IHC-derived image analysis of adult mouse midbrain sections. a) SN DA neurons in coronal midbrain section from adult mice after DAB-IHC for tyrosine hydroxylase (TH, brown). Nuclei are marked by hematoxylin-staining (blue). Left: input image where the SN is outlined in red. Scale bar: 200 µm. Middle and right: enlarged images before (upper) and after (lower) automated recognition of the nuclei within TH labelled cell bodies (green area) with DLAP-4. Scale bars: 50 μm (middle) and 10 µm (right). b) Left: number of TH-positive SN DA neurons/mouse quantified via unbiased stereology extrapolated via the optical fractionator method (M), or automatically (A), either via an algorithm provided by Aiforia or DLAP-4, as indicated. Right: remaining TH-positive neurons [%] in mice, treated with a neurodegenerative drug, calculated relative to the mean of the saline-treated group. Data are given as boxplots (median, 10-90 percentile) for all analysed mice (saline: N = 9; PD-drug: N = 12). Significant differences, according to two-way ANOVA with Tukey’s (neuron counts) or Kruskal-Wallis with Dunn’s multiple comparison test (percentage of remaining neurons), indicated by asterisks (ns: p > 0.05, *: p < 0.05, **: p < 0.01, ****: p < 0.0001). c) Upper: correlations of SN DA neuron counts/animal between stereology and automated analysis (left: Aiforia, right: DLAP-4), according to Pearson correlation test. Lower: corresponding proportionality constants ∝, calculated from the manual and automated cell count ratios. Stereology data and Aiforia data modified from *Benkert et al., Nat. Commun. 2019*. All data are detailed in Table S9-11 and Figure S4.

With the DLAP-4 approach, a similar relative % of remaining TH-positive SN neurons in the drug-treated mouse cohort was detected, as with stereology or Aiforia (DLAP-4: 59±21%, stereology: 63±59%, p=0.234; Aiforia: 58±23%, p=0.988). Moreover, also the absolute numbers of SN DA neurons/mouse, counted by DLAP-4 were similar to the respective stereological estimates (Saline: DLAP-4 4614±500, stereology 3959±643, p=0.744; Aiforia 7048±823, p<0.0001. Drug: DLAP-4 2708±975; stereology: 2504±922, p=0.996; Aiforia 4108±1596, p=0.016). The linear correlation coefficients between manually and automatically counted neuron numbers were similarly high for both automated approaches. However, the linear regression slope, was higher for the Aiforia algorithm compared to our DLAP-4 algorithm (1.6 vs 1.0), in line with the higher, over-counted neuron-number by the Aiforia approach (**Figure 6, S4, Table S11**).

### DLAP-5 analysis of immunofluorescence labelled cells defines a novel population of DAT-negative neurons in the lateral SN

We extended our automated cell quantification approach from DAB/BF to cells labelled by immunofluorescence (IF). IF has a greater dynamic signal range (Hofman, 2002; Peck et al, 2016), and allows the parallel quantification of more than one cell-population, defined by distinct markers (Katikireddy & O’Sullivan, 2011). Moreover, it can provide additional information regarding protein localization and relative expression levels.

We trained DLAP-5, based on TH and DAPI signals only, to detect and count immunofluorescence TH-positive SN DA neurons in DAPI co-stained sections, and to mark the neuronal cell body and the nucleus of TH-positive neurons. After identification of TH-positive cell bodies and nuclei, in a next step, DLAP-5 quantifies IF-intensities separately in both identified compartments for currently up to four distinct fluorescent channels, enabling co-expression analyses. **Figures 7a and S5a** illustrate the automated identification of TH IF-positive SN neurons (red) in a coronal mouse midbrain section.

**Figure 7:**
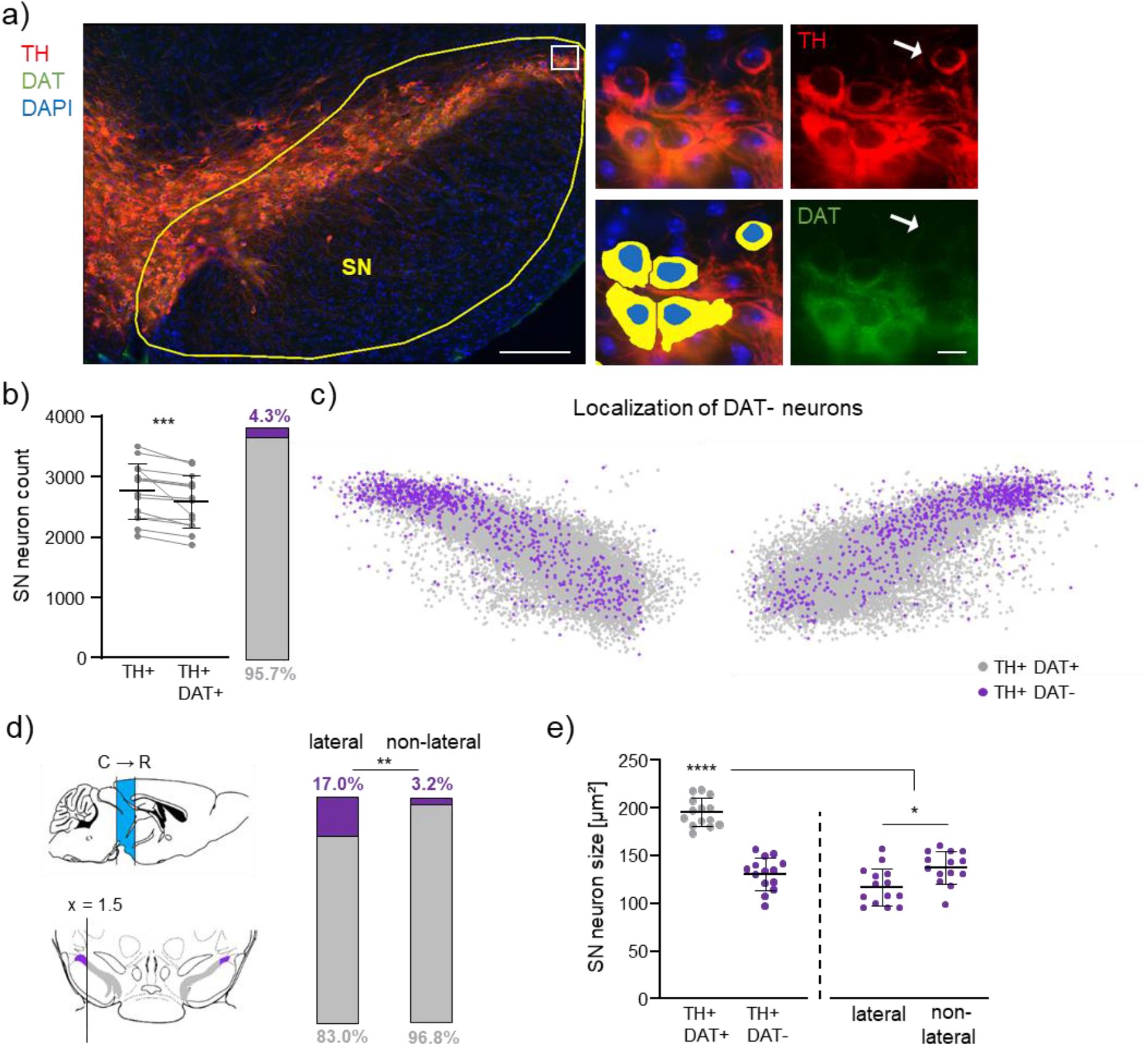
Automated (DLAP-5) image analysis of IF-derived signals identifies a DAT-negative subpopulation of TH-positive SN neurons. a) Left: neurons in a coronal midbrain section from adult mice after IF for tyrosine hydroxylase (TH, red). Nuclei are marked by DAPI-staining (blue), the SN is outlined in yellow, the white box in the lateral SN indicates the location of DAT immuno-negative (DAT-) neurons. Middle: enlarged image of TH-positive neurons within the white box from the left, before (top) and after (bottom) automated recognition of TH-positive cell bodies (yellow) and nuclei (blue) with DLAP-5. Right: single channel input image, for TH-IF (red, top) and DAT-IF (green, bottom). White arrows point to a TH-positive (TH+), DAT-negative (DAT-) SN neuron. Scale bars: 200 μm (left) and 10 µm (middle, right). b) TH-positive (TH+) and TH- and DAT-positive (TH+ DAT+) neurons/mouse and their relative amounts, quantified via DLAP-5. Given are the numbers of TH+ neurons that were further analysed for semi-quantitative DAT-analysis (compare Figure S5b for criteria). Data are given as scatterplots and mean±SD for all analysed mice (N = 14). Significant differences according to Wilcoxon tests (***: p < 0.001). c) Plotted are the individual TH+ neurons for all analysed animals (n = 38504, N = 14), according to their scaled x,y-coordinates, determined via DLAP-5. The resulting anatomical 2D-maps display the medio-lateral distribution of the TH+ DAT-neurons (violet) within the SN (TH+ DAT+ in grey). d) Left: sagittal and coronal mouse brain sections, modified from (Paxinos & Keith B. J. Franklin, 2007), illustrating the analysed caudo-rostral extent of the SN (bregma: -3.9 to -2.7, sagittal, blue), and the definition of its lateral parts in coronal sections (defined as >1.5 scaled x-units (>377.8 µm) lateral from each SN hemisphere-center (0.0); lateral: violet, non-lateral: grey, compare Figure S6). Right: relative amounts of TH+ DAT-neurons in the lateral and non-lateral SN. Significance according to Fisher’s exact test (**: p < 0.01). e) Mean cell body sizes (x,y-area) of TH+ DAT+ and TH+ DAT-SN neurons as indicated, determined via DLAP-5. Data are given as scatterplots and mean±SD for all analysed animals (N = 14). Significant differences according to Mann-Whitney tests (*:p<0.05). All data are detailed in Table S12, S16, S17, Figure S5, S6, S7.

As the expression of the dopamine transporter DAT is commonly used as an additional maker for SN DA neurons besides TH, we used DLAP-5 to quantify the relative DAT-IF signals in the marked TH-positive cell bodies. Our central aim here was to quantify the co-expression of TH and DAT in SN neurons, and their relative expression levels, rather than determining absolute numbers of TH-positive SN neurons (**Figure 7-9, S5-S7**). Thus, we only included those TH-positive SN neurons into further quantitative DAT analysis that were clearly separated/segmented, with marked cell bodies of proper size and clearly visible full nuclei. Our exclusion criteria are specified in methods and in **Figure S5b**, and resulted in manual exclusion of ∼30±6% of the automatically identified TH-positive SN neurons (**Figure S5c, Table S12**). In the remaining number of TH-positive SN neurons (2750±462 neurons/mouse, N=14), we detected only in ∼95% a DAT-IF signal above the background signal (2633±442 neurons/mouse, p=0.0001). The remaining ∼5% (4.3±1.7%) of TH-positive SN neurons were immunofluorescence negative for DAT (**Figure 7b, S5d, Table S12, S13, S16**).

In order to further characterize this small subpopulation of TH-positive but DAT-negative SN neurons, we used the x,y,z -coordinates of the analysed TH-positive neurons to generate 2D and 3D anatomical maps for each analysed coronal section, and merged maps for all mice **Figure 7c, 8, S6, online-Figure S7, Table S16**). These maps revealed that the TH-positive DAT-negative neurons were clustered in the very lateral parts of the SN, with 17% of all TH-positive neurons being DAT-negative compared to only ∼3% in the whole non-lateral SN (lateral – defined as >1.5 scaled x-axis units (>377.8 µm) lateral from each SN hemisphere-center (0.0): DAT negative: 17.0±5.8%, non-lateral: 3.2±1.6%, **Figure 7d, S6**). This location excludes that they constitute a contamination by VTA DA neurons, overlapping with the medial

SN. Moreover, the 3D-maps identified a ∼7-fold enrichment of the DAT-negative TH-positive neurons in the caudo-lateral SN, compared to its rostral parts (caudal: 36.7±14.8%, medial: 15.0±5.3%, rostral: 5.0±5.3%, p<0.0001; **Figure 8, S6/7, Table S12**), and a ∼12-fold enrichment compared to the non-lateral parts of the SN. Moreover, cell bodies of the DAT-negative TH-positive SN neurons were about 35% smaller compared to the DAT-positive neurons (DAT-positive: 195±15 µm², DAT-negative: 131±17 µm², p<0.0001), with those of the lateral DAT-negatives being about a further 20% smaller than their non-lateral counterparts, but no further differences between the caudo-lateral and rostro-lateral neurons (DAT-negative: lateral 118±20 µm² vs. non-lateral 138±17 µm², p<0.011, caudo-lateral 116±20 µm² vs rostro-lateral 124±33 µm², p=0.650, **Figure 7e, Table S17**).

**Figure 8:**
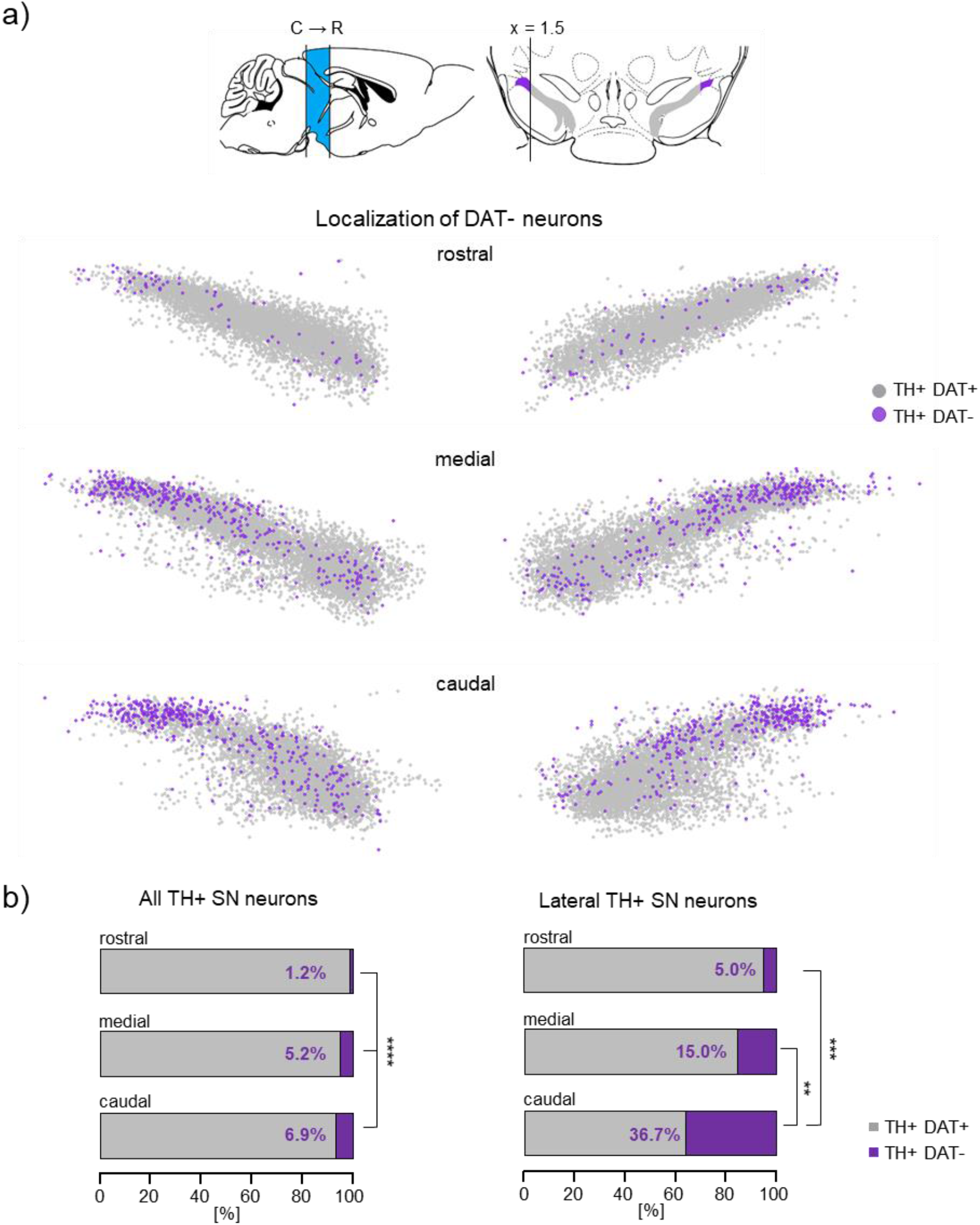
Caudo-rostral distribution of DAT-negative TH-positive mouse SN neurons. a) Upper: illustration of the caudo-rostral extent of the SN (bregma: -3.9 to -2.7), the definition of its lateral parts in coronal sections, and colour coding, as in Figure 7d, S6. Lower: plotted are the individual TH-positive neurons according to their scaled x,y-coordinates, separately for rostral, medial and caudal sections (according to bregma: rostral: -2.7 to -3.1 mm; medial: -3.1 to -3.5; caudal:-3.5 to -3.9; N = 14; rostral: TH+ DAT-n = 145, TH+ DAT+ n = 12263; medial: TH+ DAT-n = 742, TH+ DAT+ n = 13821; caudal: TH+ DAT-n = 797, TH+ DAT+ n = 10736). b) relative abundancies of TH+ DAT+ and TH+ DAT-neurons, as indicated. Significant differences according to Chi square tests (**: p < 0.01, ****: p < 0.0001). All data are detailed in Table S12. Corresponding 3D maps are given in Figure S7 (interactive HTML-file).

For subpopulations of VTA DA neurons, DAT-expression gradients have been described (Fu et al, 2012; Li et al, 2013). To systematically address whether a gradient in DAT expression is present within the SN, we used the relative, non-saturated DAT fluorescence signal intensities (DAT-RF) of each analysed TH IF-positive neuron, determined by DLAP-5, and the respective x,y,z-coordinates to generate heat-maps of RF-signal intensities (**Figure 9a, S7**). Those maps indicated a medio-lateral as well as a caudo-rostral DAT-protein gradient, with lowest DAT-immunosignals (within the background-range) in the very (caudo-) lateral SN. Corresponding heat-maps for TH suggested a similar, but much less prominent gradient (**Figure 9b, S7**), similar as previously described (Blanchard et al, 1994; Fu et al, 2012).

**Figure 9:**
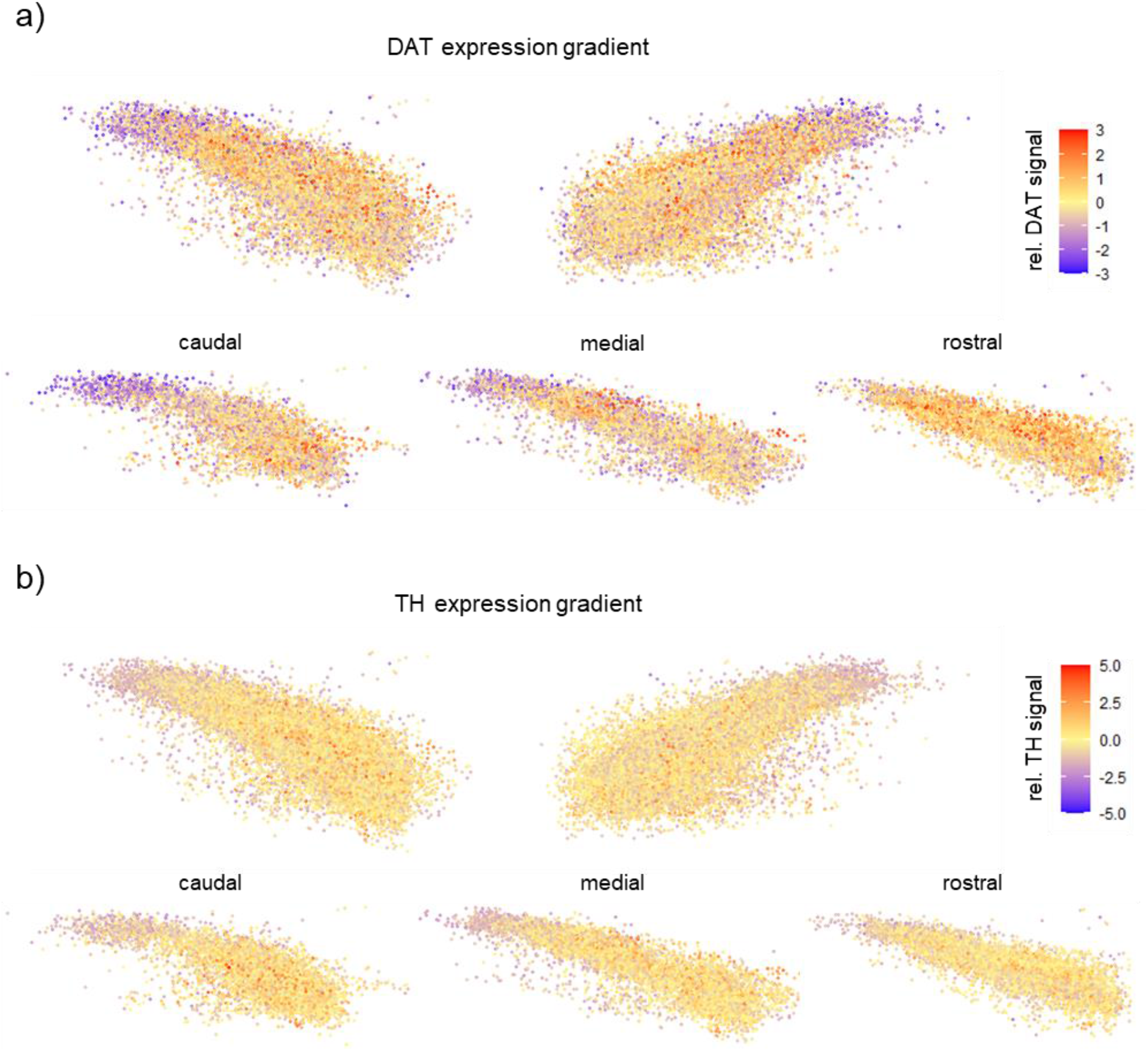
IF-derived relative TH- and DAT-protein expression in TH-positive SN neurons. Plotted are the individual TH-positive neurons from Figure 8, according to their scaled x,y-coordinates, determined via DLAP-5, for all analysed animals (n = ∼38500, N = 14). Scaled relative (rel.) fluorescence-values of DAT (a) and TH (b) signals are colour coded according to their individual deviation from the scaled mean-values (0.0) for each animal. Corresponding 3D maps are given in Figure S7 (interactive HTML-file).

To further characterize these atypical TH-positive but DAT-negative lateral SN neurons, we next analysed the co-expression patterns and expression-maps of the dopamine-D2-autoreceptor (D2-AR), the Ca^2+^ binding protein calbindin-d28k (CB), and the aldehyde dehydrogenase 1 (Aldh1A1), all markers for subpopulations of SN DA neurons. D2-AR is known to be abundantly expressed in classical (DAT-positive and CB-negative (Dopeso-Reyes et al, 2014; Lammel et al, 2008)) SN DA neurons (Ford, 2014; Reyes et al, 2013), and accordingly, we robustly detected D2-co-expression in almost all (∼93%) TH- and DAT-positive SN neurons (**Figure 10a, 11a, S8, Table S14-17)**. In the DAT-negative SN neurons however, D2-AR co-expression was significantly less abundant (∼70%, p<0.0001; N= 3), with no further difference between neurons in the lateral and non-lateral SN. The cytosolic Ca^2+^ binding protein CB is a marker for SN DA neurons that are less vulnerable in PD-paradigms (Brichta & Greengard, 2014; Garritsen et al, 2023; German et al, 1992; Liang et al, 1996). Accordingly, and in line with previous publications (Fu et al, 2012; Gerfen et al, 1985; Liang et al, 1996; Neuhoff et al, 2002; Ricke et al, 2020), we detected only in a small percentage of TH- and DAT-positive SN neurons co-expression of CB (9±1%, n= 865 from 8446 neurons, N=3); **Figure 10b, 11b, S8b, Table S14-17**), and the cell bodies of these neurons were ∼20% smaller compared to the CB-negative SN DA neurons (TH&DAT+ CB+: 173±7 µm^2^, TH&DAT+ CB-: 195±8 µm^2^, p<0.0001). In contrast, in the TH-positive but DAT-negative SN neurons, the CB co-expression rate was ∼4-fold higher (39±7% n= 143 from 383 neurons; p<0.0001, **Figure 11b**). Additionally, those neurons had the smallest cell bodies from all four TH-positive groups; they were only about half of the size of that of classical DAT-positive and CB-negative SN DA neurons (DAT-CB+: 104±8 µm^2^; p<0.0001). Moreover, the DAT-negative but CB-positive neurons clustered in the lateral SN, with a ∼7-fold higher co-expression rate (67%), compared to all DAT-positive SN neurons, (**Figure 11b**), had the smallest cell bodies (96±5 µm^2^, Table S17). Among the CB-positive neurons, the relative CB-expression levels were about 12 % higher in the DAT-negative SN neurons, compared to the DAT-positives (DAT+: 2.6±0.9, DAT-: 3.0±0.8, p<0.0001). In contrast, Aldh1A1, a marker for highly vulnerable ventral tier SN DA neurons (Carmichael et al, 2021; Garritsen et al, 2023) was rather not expressed in the TH-positive but DAT-negative lateral SN neurons (**Figure 10c, 11c, S8c, Table S14-17**), compared to ∼60% co-expression in the DAT-positive non-lateral SN DA neurons, with a higher co-expression rate in more medio-ventrally located SN DA neurons, in line with previous descriptions (Anderegg et al, 2015; Poulin et al, 2020).

**Figure 10:**
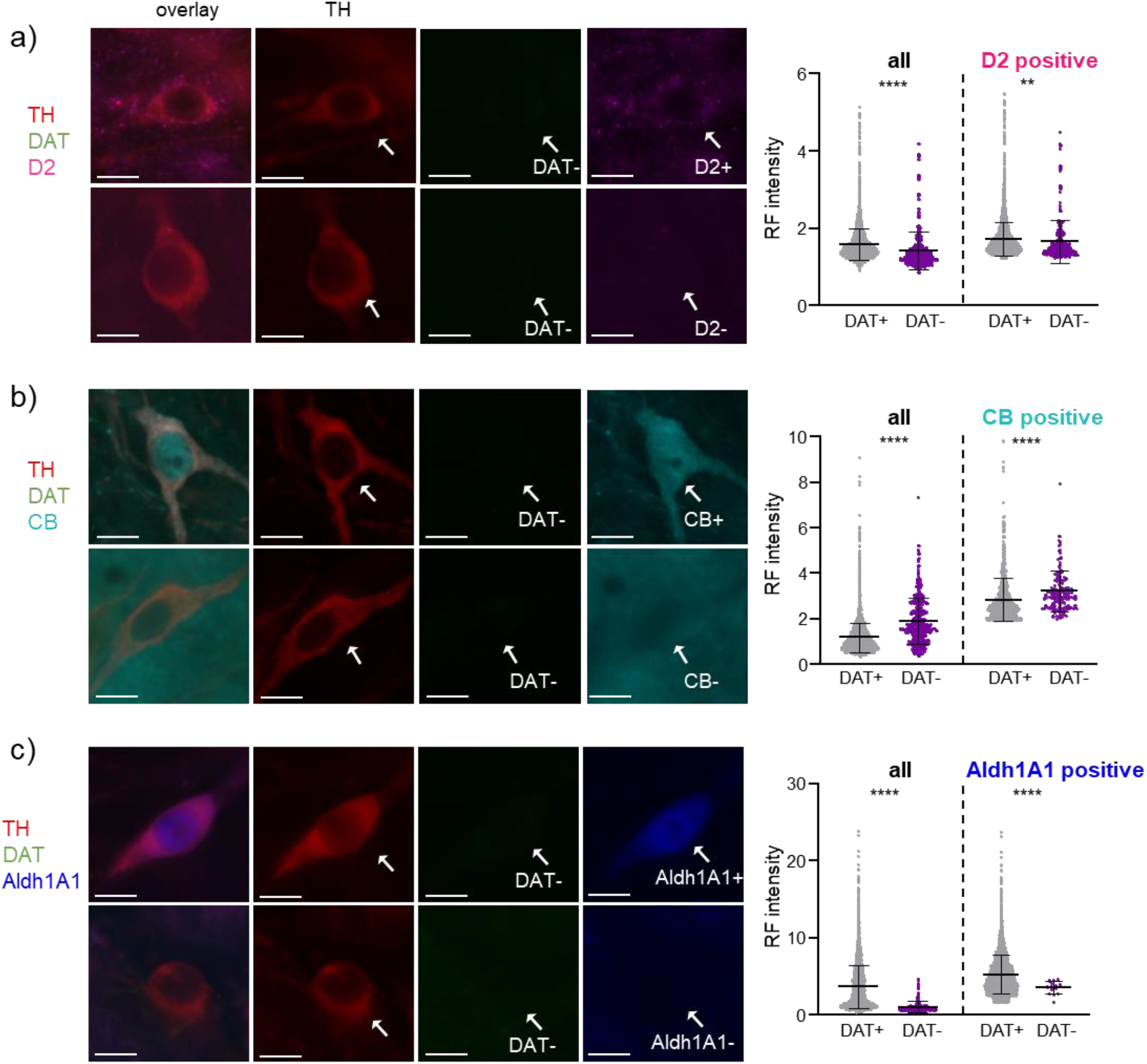
Co-expression of D2, CB, and Aldh1A1 in TH-positive DAT-negative lateral SN neurons, determined by DLAP-5. Left: SN neurons in coronal midbrain section from adult mice after IF for tyrosine hydroxylase (TH, red), dopamine transporter (DAT, green) and a) dopamine D2 autoreceptor (magenta), b) calbindin (CB, cyan), or c) aldehyde dehydrogenase (Aldh1A1, blue). Scale bars: 10 μm. White arrows indicate TH-positive, DAT-negative neurons that are positive (top) or negative (bottom) for the respective target gene. Right: relative fluorescence (RF) signal intensities for D2, CB, and Aldh1A1 in DAT-positive (DAT+) and DAT-negative (DAT-) neurons for all analysed TH-positive neurons (left), and for those TH-positive neurons only that were also immuno-positive for D2, CB, or Aldh1A1, respectively (right). Data are given as scatterplots, and mean±SD for all analysed animals (N = 3; DAT+ n = 722-8206, DAT-n = 14-383). Significant differences according to Mann-Whitney tests (**: p < 0.01, ****: p < 0.0001). Data are detailed in Figure S8 and Table S14, S15.

**Figure 11:**
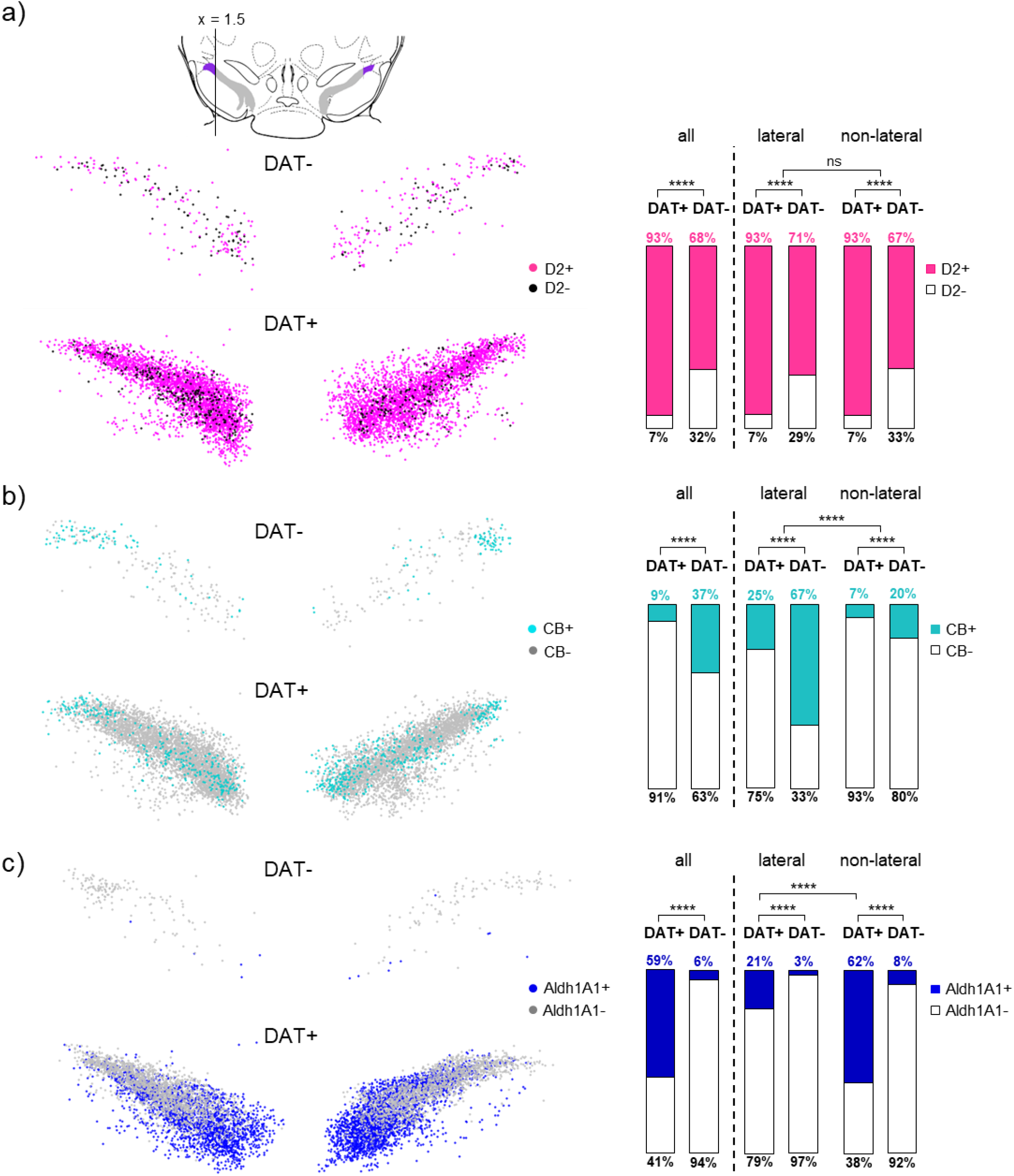
Co-expression of D2, CB, and Aldh1A1 in TH-positive SN neurons. a) Upper: illustration of the definition of the lateral SN, as in Figure 7d, S6. Left: plotted are the respective co-expression patterns of individual TH-positive neurons, as indicated, for all analysed animals (N = 3; DAT+ n = 124-7341, DAT-n = 3-209), according to their scaled x,y-coordinates, determined via DLAP-5 (similar as in Figure 7c). The resulting anatomical 2D-maps display the medio-lateral distribution of a) D2, b) CB, and c) Aldh1A1 in the DAT-negative and DAT-positive SN neurons. Right: respective co-expression rates in all TH-positive SN neurons, and separately for the lateral and non-lateral SN, as indicated. Significant difference according to Chi square tests (****: p < 0.0001). Data are detailed in Table S16 and Figure S8.

To our knowledge, neurons in the very caudo-lateral SN that are immuno-positive for TH but immuno-negative for DAT and, a medio-lateral and caudo-rostral DAT-gradient in the SN, has not yet been systematically described. We enabled this by analysing DAT-expression in about 40,000 SN neurons - hardly possible without the automated DLAP-5 approach. However, this algorithm does not allow conclusions regarding the sub-cellular location of the IF-signal.

### DLAP-6 based relative quantification of immuno-fluorescence derived signals in cellular compartments

As proteins can mediate different functions in dependence of their cellular localization, analysis with higher resolution is desired. To enable relative quantification of immunofluorescence signals in plasma-membranes, cytoplasm and nucleus, we developed the DLAP-6 algorithm, optimized to detect and analyse neurons in high-resolution confocal fluorescent images.

DLAP-6 was specifically trained by using TH as marker for the cell body/cytoplasm, DAPI as marker for the nucleus, and the ion channel subunit Kv4.3 as marker for the plasma-membrane of SN DA neurons. Kv4.3 is not a specific marker for dopaminergic neurons, but it is highly expressed in plasma-membranes of SN DA neurons, and it is the pore-forming subunit of voltage and Ca^2+^ sensitive A-type K^+^ channels that modulate the activity and the vulnerability of SN DA neurons (Aidi-Knani et al, 2015; Dragicevic et al, 2015; Haddjeri-Hopkins et al, 2021; Liss et al, 2001; Serôdio & Rudy, 1998; Subramaniam et al, 2014). The Ca^2+^-sensitivity of Kv4.3 channel complexes is mediated by plasma-membrane associated KChip3. However, in the cytoplasm, KChip3 is also known as the enzyme calsenilin, regulating presenilins, and in the nucleus, KChip3 acts as gene-transcription-repressor DREAM (downstream regulatory element antagonist modulator), illustrating the importance of subcellular localization-analysis (Burgoyne et al, 2019; Naranjo & Mellström, 2012). A prerequisite for DLAP-6 training is the availability of a very well-suited antibody to detect and separately analyze signals in plasma-membrane, cytoplasm and nucleus of DA neurons. Therefore, we utilized a suitable Kv4.3 antibody that we and others had already used for immunoelectron-microscopy (Haddjeri-Hopkins et al, 2021; Liss et al, 2001; Subramaniam et al, 2014).

**Figure 12a** shows a representative confocal image of a TH-positive SN neuron in a coronal mouse brain section, after IF-staining, before automated cell body detection (input image), and after manual as well as algorithm-based identification of the cell cytoplasm (TH, red), the nucleus (DAPI, blue), and the plasma-membrane (Kv4.3, green). As expected, the highest relative fluorescence (RF) signals for each gene were detected in these respective compartments, with manual as well as DLAP-6 analysis (RF for TH in cytoplasmic compartment: manual 55±3% vs. DLAP-6 57±4%, p=0.5862; DAPI/nucleus: manual 76±7% vs. DLAP-6 76±8%, p=0.9270; Kv4.3/membrane: manual 75±8% vs. DLAP-6 68±9%, p<0.0001; **Figure 12b, Table S18, S19**).

**Figure 12:**
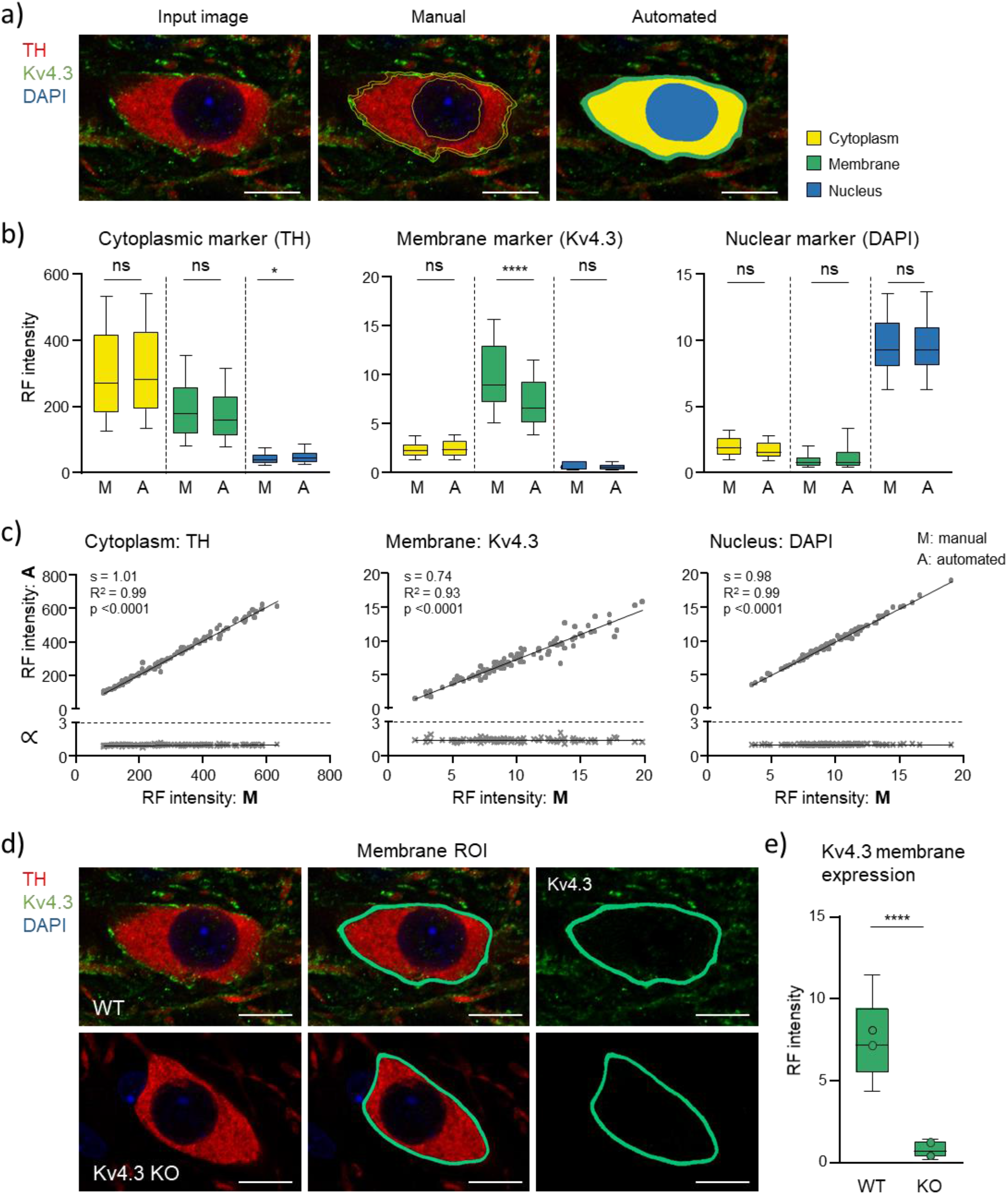
Comparison between manual and automated (DLAP-6) IF-derived image analysis in cellular compartments of TH-positive SN neurons. a) SN DA neurons in coronal midbrain section from adult mice after IF for tyrosine hydroxylase (TH, red) and the K^+^ channel α-subunit Kv4.3 (green). Nuclei are marked by DAPI-staining (blue). Left: input image. Middle: image after manual labeling of cell body, nucleus and membrane. Right: image after automated recognition of the TH labelled cell body (yellow area), nucleus (blue area) and membrane (green area) with DLAP-6. Scale bars: 10 μm. b) Relative fluorescence (RF) intensity/cell in different sub-cellular compartments, quantified manually (M) or automatically (A), as indicated. Data are given as boxplots (median, 10-90 percentile) for all analysed neurons (n = 94). Significant differences according Mann-Whitney tests (ns: p > 0.05, ****: p < 0.0001). c) Upper: single neuron correlations of RF intensities/cell between manual and automated analysis for all three marker genes in their respective compartment according to Pearson correlation test. Lower: corresponding proportionality constants ∝, calculated from the manual and automated signal intensity ratios. d) SN DA neurons in coronal midbrain section from adult WT (top) and Kv4.3 KO (bottom) mice after IF for TH and Kv4.3, similar as in a). e) Kv4.3 RF intensity/neuron-membrane compartment in WT and KO mice, as indicated. Data are given as boxplots (median, 10-90 percentile) for all analysed neuros (WT: n = 179, Kv4.3 KO: n = 200). Open circles indicate mean values for each analysed animal. Significant differences according to Mann-Whitney test (****: p < 0.0001). All data are detailed in Table S18-20 and Figure S9.

Particularly for detecting the RF-signal in the plasma-membrane compartment, the algorithm appeared superior or more consistent/precise than the manual analysis. Accordingly, the single cell correlation manual vs DLAP-6 is less good - but still very high - (R^2^=0.93 for membrane compared to 0.99 for cytoplasm and nucleus, **Figure 12c, Table S19**). Most importantly, the DLAP-6 algorithm has successfully learned to identify SN DA neuron plasma-membranes, irrespective of a Kv4.3 derived fluorescence signal, and thus enables a reliable identification of the membrane-compartment in the absence of a membrane-derived fluorescence-signal.

For a direct proof, we tested the DLAP-6 approach on SN DA neurons from Kv4.3 knock-out (KO) mice (**Figure 12d, S9, Table S20**). As expected, SN DA neurons from Kv4.3 KO mice did not show any Kv4.3 derived membrane-signal higher than the background, while in wildtype (WT), mean RF-values were ∼10-fold higher (DLAP-6: RF KO: 0.8±0.5, RF WT: 7.7±2.9, p<0.0001). Nevertheless, the plasma-membrane compartment was marked by the DLAP-6 approach in TH-positive neurons from KO mice with similar robustness as for neurons from wildtype mice - a nearly impossible task by manual analysis.

## Discussion

Here, we detail a deep learning-based image analysis platform (DLAP), including six pre-designed algorithms for automated image analysis that are easily adapted to distinct scientific needs, and allows rapid and unbiased cell count, as well as quantification of fluorescence signals (e.g. derived from mRNA or proteins) in a given ROI. These are common tasks in cell biology, but manual analysis is time consuming and prone to human error (Schmitz et al, 2011; Sommer & Gerlich, 2013). However, automated AI based and related approaches, like ImageJ/Fiji plugins (Schindelin et al, 2012) such as the Trainable Weka Segmentation (Arganda-Carreras et al, 2017), CellProfiler (Carpenter et al, 2006; McQuin et al, 2018), QuPath (Bankhead et al, 2017), U-Net (Falk et al, 2019), and DeepImageJ (Gómez-de-Mariscal et al, 2021), Cellpose (Pachitariu & Stringer, 2022; Stringer et al, 2021), or the HALO® image analysis platform by Indica Labs (Brodie, 2020), require significant computational skills and/or high-end hardware, a significant barrier for their routine use (Kraus et al, 2017; Levet et al, 2021; von Chamier et al, 2019). Hence, manual approaches are still widely used for image-analysis in life sciences (Brown, 2017; Helmstadter et al, 2021; Holland & Davies, 2020).

Our DLAP approach overcomes these issues by providing already pre-trained neural networks that can learn to detect new labels and extract desired features, with only a small amount of additional training-images. The six DLAP algorithms are pre-designed to suit most common life science tasks, and the flexible design allows their quick and easy adaption to other cell types, specimens, and scientific questions, by retraining them with a few additional respective images via the user-friendly web-platform. The FCN algorithm is not pre-trained, but can be easily trained from scratch, as less training data is required, because of its smaller size compared to the Deeplab3 neural network. For the loss function (Rajaraman et al, 2021) we briefly experimented with training the FCN with the dice loss instead of the common pixel-wise cross-entropy (Yeung et al, 2022), but did not find a significant improvement (see (Wang et al, 2022) for a recent discussion of these loss functions in the context of medical image segmentation).

DLAP dramatically reduces the respective analysis-time, and it is straightforward to use (utilized by under-graduate students in our lab). Our approach directly combines deep learning based segmentation with a variety of more conventional post-processing methods (such as the watershed algorithm), leading to high quality segmentation results. Moreover, as training and image-analyses are carried out via an easy-to-use web-based interface (https://wolution.ai/), and Wolution provides in case in-depth support for specific adjustments, our approach does not require any sophisticated hardware, software, or programming skills. Importantly, DLAP details are not proprietary, but all steps and strategies are provided here for free and in full detail, to facilitate distribution and use within the scientific community. These features represent important advantages in comparison to other cloud-based analysis pipelines, for example Aiforia (Benkert et al, 2019; Penttinen et al, 2018), CDeep3M (Haberl et al, 2018), or Visiopharm (Maffeis et al, 2019) that do not disclose insights into the underlying algorithms, do not allow user-based modifications, or require advanced programming experience (e.g. DeepCell Kiosk or Cellpose (Bannon et al, 2021; Stringer et al, 2021)). For example, the recent freely distributed well-suited algorithm for cell-segmentation, Cellpose2.0 did enable equally proper segmentation of TH-positive neurons from our IF-images, as our DLAP algorithms. However, in contrast to DLAP, it does not enable any automated post-processing or post-analysis of segmented ROIs for further downstream analysis (like whatershed algorithms, threshold based background exclusion, or quantification of fluorescent dot numbers or intensities) in one single package, but the user would require to write additional scripts for such downstream-application. Hence, as Cellpose2.0 relies on Python packages, despite its Graphical User Interface, basic programming knowledge is required, e.g. for troubleshooting package related issues or run-time errors. One particular additional advantage of our DLAP-6 is that it automatically segments the cellular membrane compartment, in the absence of any membrane marker, and thus enables a more detailed sub-cellular analysis.

The here detailed DLAP approach allows the systematic quantitative analysis and anatomical mapping of several thousands of neurons in reasonable time, and thus facilitates the identification and characterization of small cellular subpopulations. Accordingly, by DLAP-analysis of about 0,000 TH-immuno-positive SN neurons in PFA-fixed brain-sections from adult mice, and by generation of expression-maps according to anatomical coordinates, we defined a small subpopulation of neurons (∼5% of all TH-positive SN neurons). These neurons were immuno-negative for the dopamine transporter DAT, were mainly located in the caudo-lateral SN, and had ∼40% smaller cell bodies. Due to their localization in the lateral SN, we can exclude that the DAT-negative TH-positive neurons are VTA neurons (Anderegg et al, 2015). However, we do not exclude that they are non-dopaminergic, as TH-expression alone is not a proof of neurons being dopaminergic (Björklund & Dunnett, 2007). TH-positive but DAT negative striatal interneurons have been described (Xenias et al, 2015). However, interneurons in the SN are rare (Brown et al, 2014; Smith & Masilamoni, 2010) and rather small (∼ three times smaller than classical DAT-positive CB-negative SN DA neurons, (Liss et al, 1999). Given the still relatively large cell body size of the DAT-negative lateral SN neurons (∼120 µm² compared to ∼195 cm^2^), here we would not unequivocally conclude that these are interneurons, while we do not rule out this possibility.

DAT is a major determinant of activity and excitability of DA neurons, and of dopamine homeostasis and transmission (Bu et al, 2021; Condon et al, 2019; Miller et al, 2021). DAT is inhibited e.g. by methylphenidate and amphetamines as treatment for ADHD and depression (Kegeles et al, 2010; Torres et al, 2003), and changes in DAT expression have been reported in schizophrenia, ADHD, and Parkinson’s (Fisher, 1996; Salatino-Oliveira et al, 2018). However beyond disease states, up to now, only for VTA DA neurons lower DAT expression has been analysed (Fu et al, 2012; Lammel et al, 2008; Li et al, 2013). On the contrary, for SN DA neurons, DAT is commonly used as specific marker besides TH (Fisher, 1996; Kim et al, 2015; Poulin et al, 2020), as well as for their targeting, for instance, via DAT-Cre transgenic mouse lines, expressing the Cre-recombinase under the DAT-promotor (Anderegg et al, 2015; Lammel et al, 2015; Papathanou et al, 2019; Stuber et al, 2015; Todd et al, 2022). Our results imply that with such approaches, target-gene expression might not be affected in those lateral TH-positive SN neurons that are immuno-negative for DAT protein. However, it must be noted that we did not perform an absolute DAT-protein quantification. Hence, we explicitly do not exclude very low expression of DAT protein in the plasma-membranes of these lateral SN neurons, within the range of the respective detected DAT-background immuno-signals (similar applies for D2, CB and Aldh1A1). Such a low DAT expression would still be sufficient for successful DAT-Cre recombination (Feil et al, 1996; Song & Palmiter, 2018).

We found that the DAT-negative TH-positive SN neurons in the lateral SN were negative for Aldh1A1, a marker for more vulnerable, (medio-)ventral tier SN DA neurons, while the co-expression rate was massively (∼7-fold; ∼70% vs 8%) increased for CB, a marker for less vulnerable SN DA neurons. For classical DAT- and D2-AR-positive SN DA neurons, CB expression and/or absence of Aldh1A1 is a marker for less vulnerable neurons, suggesting that the non-classic DAT-negative neurons in the caudo-lateral SN might be less vulnerable as well. However the absence of inhibitory D2-AR in ∼30% of DAT-negative SN neurons might render them more vulnerable toward excitotoxicity, as their activity control by dopamine is missing (Dagra et al, 2021; Liss & Surmeier, 2022). On the other hand, the mesocortical VTA DA neurons, that are hardly affected in PD, express no functional D2-AR (Lammel et al, 2008; Morales & Margolis, 2017). For more lateral SN DA neurons, a higher sensitivity towards PD-stressors is described (Betarbet et al, 2000; Damier et al, 1999; Gibb & Lees, 1991; Okamura et al, 1995; Rodríguez et al, 2001). In terms of axonal projections, it is described that the (caudo-) lateral SN DA neurons project into the (rostro-) dorsolateral striatum (Ambrosi & Lerner, 2022), and it corresponds to the ventro-lateral SN DA neurons in humans (Düzel et al, 2009; McCutcheon et al, 2019). These neurons have been described to display a higher vulnerability in Parkinson’s compared to rostro-medial SN DA neurons (Damier et al, 1999; German et al, 1989; Hassler; Jellinger, 2012; Kordower et al, 2013). However, whether these neurons are DAT-positive or DAT-negative has not yet been analysed, and in general, there is no clear correlation between DAT-expression and vulnerability (González-Hernández et al, 2010), suggested in earlier studies (Hurd et al, 1994; Uhl et al, 1994). Hence, additional studies are necessary to address the vulnerability of the here identified DAT-negative caudo-lateral SN neurons.

Very lateral SN DA neurons have been reported to be positive for the vesicular glutamtate transporter Vglut2, and rather negative for the transcription factor SOX6 (Anderegg et al, 2015; Garritsen et al, 2023; Poulin et al, 2020). In line, our preliminary single nuclei sequencing data indicate that the here described DAT-negative SN neurons are also Vglut2-positive, while SOX6 expression is at least significantly reduced, and they confirmed the absence of Aldh1A1 in TH-positive and DAT-negative SN neurons at the mRNA level. In accordance with this and the high degree of CB co-expression, in a similar approach, it recently has been shown that one type of SN DA neuron clusters preferentially expressed SOX6 mRNA while other clusters preferentially expressed CB (Kamath et al, 2022).

What is known about the function of lateral SN DA neurons? Vglut2- and CB-positive but Aldh1A1-negative lateral SN DA neurons are important for responses to novel cues and salience (Carmichael et al, 2021; Garritsen et al, 2023; Kim et al, 2015; Menegas et al, 2017; Todd et al, 2022). Sox6-negative SN DA neurons have been shown to project to the medial, ventral, and caudal striatum and respond to rewards (Pereira Luppi et al, 2021). Some lateral SN DA neurons are activated by aversive stimuli or cues that predict aversive stimuli (Matsumoto & Hikosaka, 2009). However, these studies either analysed only DAT-positive neurons or they did not report whether the analysed neurons were DAT-positive or -negative. Lateral SN DA neurons *in vivo* display a higher burst activity (that is particularly metabolically demanding) compared to medial SN DA neurons (Farassat et al, 2019). *In vitro*, the pacemaker frequency of lateral but not medial SN DA neurons is positively coupled to the activity of Cav1.3 L-type voltage gated Ca^2+^ channels (Shin et al, 2022). These channels have been linked to the vulnerabilities of SN DA neurons in PD (Duda et al, 2016; Liss & Surmeier, 2022; Ortner, 2021; Sandoval et al, 2022). However, it should be noted that the definition of the “lateral SN” in this – and most of the studies cited above - is much broader, and likely includes the here defined very lateral SN DA neurons (compare Figure S6), but also more medial, DAT-positive neurons in the lateral SN. One recent study has indeed addressed the electrophysiological properties exclusively of only very lateral SN DA neurons, expressing CB or Vglut2, most likely similar to the very lateral DAT immuno-negative neuronal population that we describe here. The cell bodies of these neurons were also significantly smaller than those of classical SN DA neurons, had a higher input resistance and excitability, and they displayed a less precise pacemaker, with lower frequencies and less prominent after-hyperpolarisations, compared to classical SN DA neurons (Sansalone et al, 2022). However, future studies are necessary to fully characterize the here defined and quantified subpopulation of lateral SN DA neurons, in terms of their distinct molecular identity, their axonal projections, their physiological function, and their fate in disease.

## Author contributions/Acknowledgment

NB carried out the RNAscope analysis. SR carried out the protein quantification in cellular compartments, and co-supervised MH and MO with neuron count and protein quantification in IF images. DW and MM implemented and customized all algorithms for automated analysis. SM performed IHC and IF staining, JW performed stereology and automated cell count analysis for juvenile mice, HH and JB for adult mice. JRF and JMG provided Kv4.3 KO mice, and high resolution Kv4.3 immunofluorescence images. MF performed model-based data adjustments. CP generated anatomical maps, JD and RP helped with supervision and analysis. BL designed and supervised the study, and wrote the manuscript together with NB and SR. All authors revised the manuscript.

We are particularly grateful to the brain donors and the German BrainBank. We thank Matthias Bayerle and Dennis Kätzel for help with initial DAT-immunohistochemistry. This study was supported by the German DFG (LI-1745/11-3, GRK1789, SFB1506), the Austrian FWF (F44-12), the Alfried Krupp Foundation, the Boehringer Ingelheim Ulm University BioCenter (BIU), a Wellcome Trust Collaborative Award, and a Research Fellowship by the Hamburg Institute for Advanced Study (HIAS), all to B.L. NB was supported by the International Graduate Schools of Molecular Medicine and of Aging (CEMMA) at Ulm University. MH was supported by an “Experimental Medicine” scholarship of Ulm University.

### Data availability statement

The minimum dataset, all accession codes, unique identifiers, and web links are given in the manuscript. All original data and further information to interpret, verify and extend the research in this article are available upon request.

### Code availability

Further information regarding the utilized algorithms and the individual utilized algorithms are available upon request.

## Methods

### DLAP algorithms 1-6; neural networks, training, post processing parameters

#### Convolutional neuronal networks - DeepLab3 and FCN

We have used two very different neuronal network architectures, based on either the DeepLab3 network architecture or on a fully convolutional neural network (FCN). DeepLab3 based networks are more powerful for detecting complicated larger objects, like complex neuronal structures, and beyond, while FCN perform in general better for fine structures (like RNAscope probe derived, small individual dots), rather than large global concepts (Chen et al, 2017; Long et al, 2015). Both types of neural networks belong to the class of convolutional neuronal networks (CNN), commonly used for image analysis (Greener et al, 2022). Convolutional layers do not receive their inputs from the whole previous layer but only from a certain area (called ‘filter’ or ‘kernel’) which increases their computational efficiency (Wang et al, 2019). Moreover, they assign to each pixel in the image a class, such as cellular nucleus or background. This type of analysis is called semantic segmentation. The performance of deep learning based semantic segmentation depends strongly on the neural network architecture, training parameters and in particular the quality of the training dataset. By specific training and further optimizing these two network-types for distinct tasks, we generated six distinct algorithms, as specified in **Table1 and 2**.

For cell counting and immunofluorescence signal quantification (DLAP-3 to 6), the large pre-trained neural network DeepLab3 (Chen et al, 2018) was further trained and optimized post training for the respective specific tasks. DeepLab3 is one of the best performing image segmentation algorithms on the Pascal Visual Object Classes (VOC) dataset (Everingham et al, 2006). The Pascal VOC is a standardized image dataset for building and evaluating algorithms, often used as a benchmark for segmentation quality. In general, the key innovation of DeepLab algorithms is the use of so called dilated or atrous convolutions. This type of convolutions has a dilated filter size, increasing the area covered by each filter, and the context that can be incorporated. Therefore, they have a higher resolution and can integrate information over larger areas, while keeping the number of parameters of the convolutions the same (Chen et al, 2017). Specifically, for our algorithms 3 to 6, we used xception65 as the CNN backend, provided by Google [https://github.com/tensorflow/models/tree/master/research/deeplab] (Chollet, 2017), which was pre-trained on the Pascal VOC 2012 segmentation dataset (Everingham et al, 2015).

For analysis of RNAscope data, we developed a more simple, better-suited FCN (DLAP-1 and 2), as the trained respective algorithms based on DeepLab3 were not optimal for reliable detection of RNAscope-derived dot-numbers (**Table S1**). The most important modification of the FCN architecture is that it uses only convolutions, rather than including fully connected linear layers (Long et al, 2015). Our FCN has a smaller receptive field and conserves image resolution at each convolutional layer. Replacing the linear layers with fixed dimensions by convolutions allows the network to accept arbitrarily large images for processing and making predictions (called interference), and to process them very efficiently - limited mainly by the graphics processing unit (GPU) memory size. The FCN architecture, illustrated in **Figure 2**, is our own development, using the Tensorflow library [https://www.tensorflow.org/]. Our FCN network uses five convolutional layers, connected by the Rectified Linear Units (ReLU) activation function. This function only transfers the direct output of the previous layer to the next layer, if it is positive, while it returns zero for negative outputs (Nair & Hinton, 2010). The softmax activation function normalises the network output to the probability distribution. Our network does not include any downsampling/downsizing to reduce image resolution through dropout or strided (kernel step size > 1) convolutions. Therefore, the output image has the same resolution as the input image, and fine structures are not smoothed out. We used another approach of neural network architectures, the so-called 1×1 convolutions, to reduce the number of feature maps and keep the parameter number tractable (Lin et al, 2013). Such a convolution is used between feature map 2 and feature map 3 to reduce the number of kernels by a factor of 4, thus reducing the parameter size of the following convolution by a factor of 16. To conserve image dimension, we extend the outer frame at the boundary of each image, which increases the space for the filter/kernel to scan the image, a process also called mirror padding. This architecture provided a good compromise between simplicity (and thus computational speed) and segmentation quality.

#### CNN training parameters for six individual tasks

Training dataset need to have sufficient quality and size, depending on data variability (Moen et al, 2019; von Chamier et al, 2019). However, we observed that using too large training datasets lead to overfitting, which resulted in decreased generalization. We ensured the quality (e.g. resolution, frame size) of our training datasets, and performed a quality evaluation during training (see below), in addition to data-validation by comparison with manual analyses. Images showing different neuronal shapes/forms in different brightness intensities were used for training to ensure the identification and recognition over a vast array of brightness and cellular structure.

As our FCN is not based on a pre-trained neural network (no “transfer learning”), it was trained by us from scratch, by only using our own training data. This is possible because of its smaller size compared to the Deeplab neural network, which also means that less training data are required. The FCN-based algorithms 1 and 2 analyse image patches of size 63 x 63 pixels. To train the network, we randomly cropped the images from our manually labelled training dataset (ground truth data) into such patches. Training was performed using the stochastic first-order gradient-based Adam optimizer with default settings (learning rate α=0.001, estimates decay rates β_1_=0.9, β_2_=0.999, to prevent division by zero ε=10^−8^) (Kingma & Ba, 2014). For DeepLab3 based algorithms DLAP-3 to 6, we adjusted the neural network weights by further training with our own datasets using the training script provided by Google Research (Chollet, 2017). This was performed on randomly sampled patches of size 512^2^ pixels much larger than the 63^2^ pixels we use in the FCN algorithm, as Deeplab3 is designed to detect large, complex objects within an image.

For both network-types, the training time on a modern GPU was several hours for each of the datasets. The number of images and neurons each that were used to train each of the individual algorithms is given in **Table 2**. We used 80% of manually labelled images for the training dataset, and the remaining 20% of the images for the “test data” set. Train-test-split on the level of images was chosen to avoid data leakage. For this performance-evaluation of the individual algorithms, the “test data” was analysed and the results were compared to the ground truth provided by the labels. For all algorithms, we assessed typical quality measures (**Table 1**, **S1**): The pixel error rate, i.e. the global percentage of pixels that are wrongly classified. This global error rate was calculated by defining for each segmentation class the rates of true positives (TP), true negatives (TN), false positives (FP) and false negatives (FN), each given in % of the total pixels of the whole training dataset. The specificity [defined as TN/(TN+FP)] and the sensitivity [defined as TP/(TP+FN)], for each segmentation class, were also calculated, as these values are more suitable measures in cases of large class imbalance.

We trained each dataset until the training error/loss function (difference between output and ground truth) converged to a minimum. We also use “Early Stopping” in the training process to avoid overtraining (see above). We verified that we did not overtrain, by tracking the pixel error in the “test data”. If overtraining is happening, only the training error is still getting lower, while the test error increases. Training was stopped as soon as the pixel error was getting worse not better. Performances of all algorithms were evaluated on the respective “test data” (compare **Table 2**).

#### Post training image analysis optimization

After CNN training was completed, the quality of the CNN outputs for each task (summarized in **Table 1**) was further improved by, implementing additional classic image analysis procedures (a) to (e), specified below, into the individual CNN procedures. If an analysis step was further increasing the correlation between algorithm- and manually-obtained results for a distinct task, it was implemented into the respective algorithm. The specific steps (a-e) that were implemented post-training into each of the final algorithms are summarized in **Table 2**.

a. *Watershed algorithm.* This algorithm requires a set of markers, which are associated with the centers of respective target structures (Kornilov & Safonov, 2018). These markers are obtained by calculating the Euclidean distance transform of the binary mask of target structures cells (detected by the neural network), and then finding peaks in the distance image. Starting from these markers, the watershed algorithm “floods” the image until all pixels are assigned with a watershed region. We used this algorithm in both FCN based algorithms, for a better separation of mRNA molecule derived dots. The FCN output tend to connect single fluorescent dots of individual mRNA molecules, which biases their number count. We found that by applying a classical watershed algorithm, we can separate these points and get a number count much closer to the truth (manual count). For the DeepLab3 based algorithms 4 and 5, watershed algorithm were used to improve segmentation of neighboring nuclei/cells. This was not necessary for DLAP-3.
b. *Morphological operations*. A series of morphological operations of type dilation and erosion (Fisher, 1996) were applied to algorithms 1 and 2. These operations add to or remove pixels from the object boundaries. This improves separation of individual neurons and smoothes out uneven boundaries provided by the CNN. This approach was used in both FCN-based algorithms for a better separation of individual neurons.
c. *Binary fill-holes-function.* This function was applied to the DeepLab3-based CNN outputs for DLAP-4 to 6, were pixel-wise semantic segmentation resulted in holes within the detected compartments, as such holes are of no biological meaning. Therefore, all holes in the segmentation masks of each region were filled using the “*binary_fill_holes*” function (González-Hernández et al, 2010) of the SciPy library http://scipy.github.io/devdocs/reference/generated/scipy.ndimage.binary_fill_holes.html.
d. *Remove target gene colour channel procedure.* This procedure was applied to the DeepLab3 based algorithms 5 and 6 to facilitate identification of the target areas within the ROI (i.e neuronal cell bodies or neuronal compartments), in which the relative target-gene derived fluorescence-signal was quantified. This procedure is removing the target gene channel for the algorithm-based detection of the target areas, to ensure its recognition, independently of the target gene fluorescence-signal. For excluding the target-gene channel, the respective colour layer of the image was set to zero before passing the image through the CNN (both for training and analysis), and it was added back, for the target-signal quantification step of the algorithms.
e. *RGB intensity threshold*. Such threshold was included to the DeepLab3 based algorithms 5 and 6, to optimize background intensity quantification for calculation of relative target-gene signal intensities in identified target areas (i.e. neuronal cell bodies or neuronal compartments) to correct for non-uniform intensity conditions in different images/experiments. It was not always straight-forward to determine the individual background-intensities, because some images contained large regions with no detected target areas, but also with no apparent blank background. We included RGB intensity thresholds (for widefield and for confocal images) of pixel values ranging from 10-40 for each channel, depending on its raw intensities, to exclude pixels higher than this threshold from determining background signal intensities.

### Mice

Juvenile (∼PN13, Figure 5 data), and adult (∼PN90) male mice were bred at Ulm University at a 12-hour light/12-hour dark cycle and were fed ad libitum. Kv4.3 KO mice were obtained from Jean-Marc Goaillard (Haddjeri-Hopkins et al, 2021). All animal procedures were approved by the German Regierungspräsidium Tübingen (Ref: 35/9185.81-3; TV-No. 1043, Reg. Nr. o.147) and carried out in accordance with the approved guidelines.

*Drug and saline treated animals* (Figure 4 data), were the same, cohort as already published (Cav2.3 +/+ mice from Benkert et al (2019)). Drug treated mice were treated in vivo with MPTP/probenecid to introduce SN DA neuron degeneration. Briefly, MPTP hydrochloride was injected 10 times (every 3.5 days for 5 weeks) subcutaneously at a concentration of 20 mg/kg saline (sigma) together with probenecid intraperitoneally at a concentration of 250 mg/kg in 1x PBS (Thermo Fisher). Control mice were injected with saline and probenecid. For all details, including, TH-DAB/hematoxillin staining, stereology, and Aforia-based automated analysis, (see Benkert et al (2019)).

### Human brain samples

Human midbrain tissue including the substantia nigra (SN) was collected and obtained from the German Brain Bank (www.brain-net.net), Grant-No. GA76 and GA82), as native cryo-preserved tissue blocks (-80 °C/dry ice). Analysis of the human material was approved by the ethic commission of the German Brain Bank as well as of Ulm University (277/07-UBB/se). Informed consent from all participants was given to the German Brain Bank. Detailed information on human midbrain samples, including sex, is summarized in **Table S5**. We have no information if/how the information regarding sex and gender was obtained. Human tissue was processed, stained, and the RNA integrity was assessed via determining the RNA integrity number (RIN), using the Agilent 2100 Bioanalyzer as described (Duda et al, 2018).

### RNAscope® *in situ* hybridization, image acquisition, and data analysis

*RNAscope experiments* were performed essentially as described (Benkert et al, 2019; Matsuda et al, 2009). *The* RNAscope® technology (Advanced Cell Diagnostics, ACD) was according to the provided protocol for fresh frozen tissue sections [https://acdbio.com]. Details of the used RNAscope probes are given in **Table S2**. All used chemicals were RNAse free grade. 12 µm coronal cryosections of mouse or human midbrains were prepared, using a Leica CM3050S cryotome as previously described (Gründemann et al. 2008; Duda et al. 2018), mounted on SuperFrost® Plus glass slides (VWR), and allowed to dry in a drying chamber containing silica gel (Merck) at -20°C for one hour. After fixation with 4% PFA (Thermo Fisher Scientific) in 1x PBS (pH 7.4) for 15 min at 4°C and dehydration via an increasing ethanol series (50%, 75%, 100%, 100%, Sigma), sections were permeabilized for 30 minutes by digestion with protease IV (ACD) at room temperature (RT). Following digestion, respective RNAscope probes (TH probes and target gene probes were processed in parallel) were hybridized for 2 h at 40°C in a HybEZ II hybridization oven (ACD). Signal was amplified using the RNAscope Fluorescent Multiplex Detection Kit (ACD) containing four amplification probes (AMP1-4). In between each amplification step (incubation at 40 °C for 30 min with AMP1/3) or 15 min with AMP2/4), sections were washed twice for 2 min with wash buffer (ACD). After RNAscope hybridization, sections were counterstained with 4′,6-diamidino-2-phenylindole (DAPI) ready-to-use solution (ACD, included in Kit) for 30s at RT. Sides were coverslipped with HardSet mounting medium (VectaShield), and allowed to dry overnight at 4 °C. RNAscope probes for TH were labelled with AlexaFluor488, and target gene probes with Atto550.

#### Image acquisition

Fluorescent images containing the ROI (i.e. the SN), of murine and human midbrain sections were acquired at 63x magnification using a Leica DM6 B epifluorescence microscope. All images were acquired as z-stacks, covering the full depth of cells, and reduced to maximum intensity Z-projections, utilizing Fiji [http://imagej.net/Fiji]. TH-derived RNAscope fluorescence signals were visualized at 480/40 nm excitation, 527/30 nm emission and 505 nm dichroic mirror, target gene-derived fluorescence signals were visualized at 546/10 nm excitation, 585/40 nm emission and 560 nm dichroic mirror, DAPI-signals were visualized at 350/50 nm excitation, 460/50 nm emission and 400 nm dichroic mirror. Files were aquired using the LASX software (Leica), and stored as PNG files.

#### Image analysis

For manual analysis, the images were further processed using the Fiji software [http://imagej.net/Fiji]. First, cell bodies of individual TH- and DAPI-positive neurons were encircled, using the “Polygon selection” tool to define the area for RNAscope signal quantification. In the next step, a classifier was trained on images only showing target gene signal, by manually labelling and annotating background and dot regions (around 30 labels for each class) in the “Trainable Weka Segmentation”. This classifier was then used to classify all target gene images into the two groups “dots” and “background”. Afterwards, the threshold for particle recognition was set on classified images, and dots were counted for each cell-specific ROI using the “analyze particle” function. Target probe hybridization results in a small fluorescent dot for each mRNA molecule, allowing absolute quantification of mRNA molecules (via dot counting) independent from fluorescent signal intensity.

For automated quantification of RNAscope signals, custom-designed FCN based algorithms DLAP-1 (for mouse-brain sections) and 2 (for human brain sections) and the Wolution-platform were used [https://console.wolution.ai/] (Wolution GmbH & Co. KG, Planegg, Germany). Processed images were uploaded on the Wolution platform, and the algorithms automatically marked cell bodies of TH-positive cells and quantified the number (and the dot area) of target gene derived fluorescence dots. After algorithm processing, correct identification of cells was controlled by hand and resulted in inclusion of ∼50 % correctly identified cells for analysis. If a quantification of all TH-positive neurons is required, neuronal nuclei must be counted, before excluding neurons with not optimally separated cell bodies for subsequent RNAscope based mRNA quantification in the target cells.

### Statistical modelling for brightfield light adjustment

To asses and correct for a possible influence of neuromelanin (NM) human-derived RNAscope data for, a linear mixed effect model was applied, similar as we had previously described (Duda et al, 2018; Schlaudraff et al, 2014). Briefly, we applied a bayesian probabilistic model approach conducted in RStan (Stan Development, 2018) with the R package version 2.21.7, https://mc-stan.org/) using R (version 4.2) under RStudio (version 2022.07). We applied normal distributions for priors of all parameters. To achieve a stable non-degenerate solution priors needed to be properly informed by restricting the effective parameter space. The chosen model, detailed below, is well-informed by the data on all its parameters and shows an acceptable fit on the age level (see posterior-predictive plots in **Figure S2**).

The number of detected dots/mRNA-molecules for each component/gene (defined as *Y_G_*) is assumed to follow a binomial distribution in which the trial size is derived from cell area s:

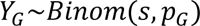

The probability pG of detecting a target molecule is expressed according to a logit-link function (linear combination) as:

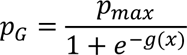

where *g*(*x*) is an equation, indexed by the mixture-component and a binary discretization of a brain donor (adult vs. aged) that considers a cell’s NM concentration (as given by brightfield light transmission, *T*) according to the Lambert-Beer law which states that concentration is proportional to the negative logarithm of light transmission

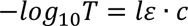

Light transmission *T* is the fraction of transmitted to incident light intensity

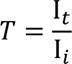

Since I*_t_* < I*_i_* and light intensities are strictly positive, it follows that 0 < *T* < 1. For the probabilistic model a distribution for T has to be defined. A continuous distribution bounded on [0, 1] is the Beta-distribution.

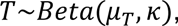

The Beta distribution is parameterized by its mean *μ_T_* and the precision *κ*.

For *μ_T_* we apply a logit-link function

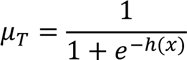

*h*(*x*) is a linear equation, indexed by the mixture component and binarized age. In addition, the intercept considers a random effect on individual brains. The mixture proportion is modeled to depend on binarized age and to consider a random effect on the brain level, given in Wilcoxon notation (Wilkinson & Rogers, 1973)

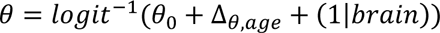

To adjust experimental results for a mutual NM influence, data were fitted and then reproduced from the model, with each cell’s *T* set to a constant value according to the mean of all brains’ *μ_T_*. The influence of BF light transmission (associated to NM content by –log *T*) on gene expression / dot detection probability can be considered significant with 95%-credible intervals from -5.1 to -1.5 or from -12.2 to -7.7 for low and high expressing cells, respectively (corresponding intervals from the prior distribution vary from -7.8 to +8.0).

### Immunohistochemistry, image acquisition, and data analysis

*DAB Immunohistochemistry (IHC)* experimental procedures were performed essentially as previously described (Benkert et al, 2019). Briefly, mice were transcardially perfused with 1x PBS-heparin (37°C) for 2 minutes followed by ice cold 4% PFA for 4 min with a flow rate of 6ml/min and post-fixed (overnight at 4°C) in 4% PFA (Thermo Fisher Scientific) in 1x PBS (pH 7.4). Brains were stored in 0.05% NaN_3_ (Sigma) in PBS at 4°C until vibratome (VT 1000S, Leica) cutting (30 μm coronal midbrain sections). All washing and incubation steps during the staining procedure were performed while shaking (300 rpm, microplate shaker, VWR). Free-floating sections were washed (three times in 1xPBS for 10 min) and blocked for 2 hours with 10% normal goat serum (NGS, Vector Laboratories), 0.2% BSA (Carl Roth) and 0.5% Triton X-100 (Merck) in 1x PBS to prevent non-specific antibody binding. After further washing (one time in 1xPBS for 10 min), sections were incubated with rabbit anti-TH primary antibody (1:5000, Merck) in 1% NGS, 0.2% BSA and 0.5% Triton X-100 (in 1x PBS) overnight at room temperature (RT). Sections were then washed three times in 0.2% Triton X-100 in 1xPBS for 10 min and incubated with biotinylated goat anti-rabbit (1:1000, Vector Laboratories) for 2 hours at room temperature. Immunostaining was visualized via VECTASTAIN® ABC system based on Horseradish peroxidase (HRP) detection (Vector Laboratories) using 3,3’-Diaminobenzidine (DAB, Vector Laboratories) as substrate. The slices were mounted on SuperFrost® Plus glass slides (VWR), dehydrated in ascending ethanol series (50%, 70%, 90%, 100%, 100%, Sigma) for 10 min each, and cleared with xylene (Sigma) two times for 10 min each. Slides were mounted with VectaMount Permanent Mounting Medium (Vector Laboratories).

*Hematoxylin-counterstaining:* For counterstaining of already DAB-stained and permanently mounted adult mouse brain sections, slides were incubated in xylene (two times for 5 min) to remove coverslips and rehydrated by an ethanol series (100%, 100%, 90%, 70%, 50%) and H_2_O (5 min each). After drying for 5 min, slides were incubated with Vector hematoxylin QS (2 min) to counterstain nuclei (Vector Hematoxylin QS Kit, Vector Laboratories). After dehydration in ethanol series and clearing in xylene they were again mounted using VectaMount (all steps as described for DAB-staining).

*Stereology:* Stereological estimates of TH-positive neuron numbers were determined using a Leica CTR5500 microscope and the unbiased optical fractionator method (StereoInvestigator software; MBF Bioscience), similar as previously described (Benkert et al, 2019; Liss et al, 2005). The SN region throughout the whole caudo-rostral SN axis (Bregma -3.8 to -2.7, according to (Paxinos & Keith B. J. Franklin, 2007) was identified, using well established landmarks (Fu et al, 2012)), and was marked as ROI for TH-positive neuron count. The ROI was marked ab analysed unilaterally on each of the serial DAB-stained sections (37 for juvenile, 40 for adult mice). Sampling grid dimensions were 75 x 75 μm (x,y-axes), counting frame size were 50 x 50 µm (x,y-axes), and counting frame height was 9 or 11 µm for juvenile and adult mice, respectively. Estimated total number of TH-positive neurons (N) was calculated for each animal according to equation (1):

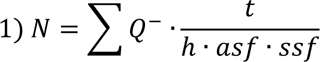

with ΣQ^-^ = number of counted neurons, t = mean mounted section thickness (i.e. ∼10-11 µm), h = counting frame height (i.e. 80 % of the mounted section thickness), asf = area sampling fraction (i.e. 0.44), and ssf = section sampling fraction (i.e. 1 for SN).

Reliability of the estimation was evaluated by the Gundersen coefficient (CE, m = 1) according to equation (2). CE values were all ≤0.05 for all analysed animals.

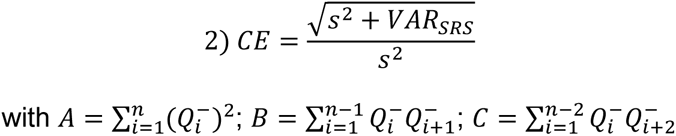

s^2^ = variance due to noise, and VAR_SRS_ = variance due to systematic random sampling, according to equation (3) for m = 1.

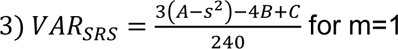

#### Automated neuron counting

Image acquisition: Digital images of DAB-stained sections were acquired using a whole slide scanner (3D-Histech Pannoramic 250 Flash III, Sysmex Deutschland GmbH, Norderstedt, Germany or Aperio Versa 8, Leica Biosystems Nussloch GmbH, Nußloch, Germany) and processed using the Fiji software [http://imagej.net/Fiji] were used to combine these layers.

Image analysis: The digital slides were processed using QuPath [https://qupath.github.io/] to label and cut out the sections of interest (37/40 consecutive SN sections covering the entire caudo-rostral axis). As for stereology, the SN ROI was identified according to the typical landmarks (Fu et al, 2012; Paxinos & Keith B. J. Franklin, 2007) and analysed unilaterally. Processed sections (showing the ROIs) were afterwards uploaded on the Aiforia® Cloud platform [https://www.aiforia.com/] (Aiforia Technologies Oy, Helsinki, Finland) or the the Wolution platform [https://console.wolution.ai/] (Wolution GmbH & Co. KG, Planegg, Germany). For both automated platforms, in case of flawed sections that could not be automatically analysed, the counts from the section before and after were averaged to prevent methodological bias (for stereology, these sections were omitted, and this information was taken account for the calculation of the stereological estimates).

The Aiforia-platform was only used for analysis of Hematoxylin-counterstained DAB-stained brain sections. It uses a supervised training to establish a non-disclosed deep CNN algorithm that recognized TH-positive neurons, based on nuclear/cell morphology and TH signal. The Aiforia algorithm was trained with 4064 TH-positive neurons (Benkert et al, 2019) and comprised two layers. The individual TH-positive cells were segmented in the first one and counted in the second one (see Penttinen et al (2018)). For generating the ground truth data used for algorithm training, one circle is placed on top of the nucleus within each TH-positive neuron (instead of marking the whole nucleus), similar as for our respective Wolution-based algorithms DLAP-3 and 4. Only TH-positive cells with a clearly visible/focused nucleus were considered for neuron counting, to avoid counting the same cell on more than one section. Processed images were uploaded on the Wolution platform. For counting of only DAB-stained TH-positive neurons (juvenile mice), DLAP-3 was used, for counting of hematoxylin counterstained TH-positive neurons (adult mice), DLAP-4 was utilized.

### Immunofluorescence, image acquisition, and analysis

*Immunofluorescence (IF)* experimental procedures were performed essentially as previously described (Benkert et al, 2019). Mouse brains were perfused, stored and cut as already described for IHC. All washing and incubation steps were performed while shaking (300 rpm, microplate shaker, VWR).

For automated counting and relative signal intensity quantification analysis with the DLAP-5 algorithm, immunostaining was performed in five separate cohorts of mice, using sequential or simultaneous antibody incubation. 30 μm coronal midbrain free-floating sections were washed (three times in 1xPBS for 10 min). An additional antigen retrieval step was applied for D2- and Aldh1A1-antibodies (100°C for 10 min in pH6 (D2) or 80°C for 30 minutes in pH 9 (Aldh1A1). Sections were blocked for 2 hours with 10% normal goat serum (NGS, Vector Laboratories) and 0.5% Triton X-100 in 1x PBS. After blocking, sections were incubated with rabbit or chicken anti-TH primary antibody (1:1000, Merck), and/or mouse anti-DAT (1:500, Thermo Fisher Scientific), chicken anti-calbindin d28K (1:1000, Novus bio), rabbit anti-DRD2 (1:200, Proteintech), rabbit anti-Aldh1a1 (1:200, Abcam) primary antibody in carrier solution (1% NGS, 0.5% Triton X-100 in 1x PBS) overnight at 4°C. For subsequent primary antibody incubations, sections were washed (three times in 1x PBS for 10 min at RT) before the next overnight incubation at 4°C. Sections were washed (three times in 1x PBS for 10 min at RT) and incubated with Alexa Fluor 488 goat anti-mouse secondary antibody (1:1000, Thermo Fisher Scientific), Alexa Fluor 647 goat anti-rabbit (1:500, Thermo Fisher Scientific), and Alexa Fluor 647 goat anti-chicken (1:1000, Thermo Fisher Scientific), in carrier solution for 3 hours at RT in darkness. After washing (three times in 1x PBS for 10 min at RT), the sections were mounted on SuperFrost® Plus glass slides (VWR) with VectaMount Permanent Mounting Medium with DAPI (Vector Laboratories).

For relative quantification of fluorescence-signal intensities in cellular compartments (plasma-membrane, cytoplasm, nucleus) with DLAP-6, two similar Kv4.3 staining protocols were utilized, adapted from (Haddjeri-Hopkins et al, 2021), leading to very similar results. More precisely, one WT and one Kv4.3 KO brain (dataset 1) was processed, and images were acquired in the Goillard lab essentially as described (Haddjeri-Hopkins et al, 2021). The other brains (dataset 2) were perfused in the Goillard lab, and further processed in the Liss lab, according to the following protocol. 50 μm free-floating coronal midbrain sections were blocked for 1 hour 30 minutes with 5% NGS and 0.3% Triton X-100 in 1x PBS, followed by incubation with a rabbit anti-Kv4.3 primary antibody (1:10000, Alomone Labs) together with chicken (G)/mouse (L) anti-TH primary antibody (1:1000, Abcam / Merck) in a carrier solution (1% NGS, 0.3% Triton X-100 in 1x PBS) overnight at 4°C. Sections were then washed (three times in 0.3% Triton X-100 in 1x PBS for 15 min) and incubated with Alexa Fluor 488 goat anti-rabbit secondary antibody (1:1000, Thermo Fisher Scientific) and Alexa Fluor 546 goat anti-mouse secondary antibody (1:1000, Thermo Fisher Scientific) in carrier solution for 2 hours at room temperature. After washing (three times in 0.3% Triton X-100 in 1x PBS for 15 min), sections were incubated with DAPI (1.5 μg/mL; Sigma-Aldrich) for 10 min, washed (two times in 1x PBS for 10 min), and mounted on glass slides using Vectashield mounting medium.

#### Image acquisition

For automated counting of TH-IF positive neurons und and relative signal intensity quantification (DAT-IF), fluorescent images of all sections were acquired with Leica DM6 B epifluorescence microscope (Leica Microsystems) using a 63X oil objective. All image parameters were set using the LAS X software (Leica microsystems) to avoid saturated pixels, and identical acquisition settings were maintained (a prerequisite for relative signal intensity quantification). TH-IF was visualized at 546 nm (exposure = 200-500 ms, gain = 1-2), DAT-IF at 488 nm (exposure = 400-600 ms, gain = 1-2), CB-IF, D2-IF and Aldh1A1-IF at 660 nm (exposure = 300-500 ms, gain = 2) and DAPI at 405nm (exposure = 60-300 ms, gain = 1). Fluorescence lamp illumination was kept at 30%. Images were acquired as tile scans from a single plane of focus. Merged images were exported as Leica Image File (LIF).

For relative quantification of fluorescence-signal intensities in cellular compartments, 63X confocal quality images were acquired using a Zeiss LSM780 with the settings defined in (Haddjeri-Hopkins et al, 2021) for dataset 1. In dataset 2, confocal images using 100X/1.4 NA oil objective were acquired with STEDYCON (Abberior Instruments) mounted on an Olympus BX53 upright microscope (Olympus LS), and image parameters were adjusted using the STEDYCON web-based software (Abberior Instruments). To avoid saturated pixels (a prerequisite for relative signal intensity quantification), photon counts from all channels were monitored in the photon count range indicator using the “fire blue” colour-map. This map indicates saturated image parts in blue over a yellow-red gradient for each individual channel. Fluorochromes were excited using an Argon laser, and the following wavelengths were used for excitation: TH-IF at 546 nm, Kv4.3-IF at 488 nm, and DAPI at 405 nm. The following acquisition parameters were adjusted at the stedycon user interface: Pixel size 78 nm; pinhole diameter 0.71 AU; one-directional line accumulation acquisition; laser power 10% for TH and Kv4.3, 12% for DAPI. Images were acquired from a single plane of focus and were exported as OsmAnd Binary Maps (OBF) files.

#### Image analysis

For automated counting of TH-IF positive neurons und and relative signal intensity quantification, images were converted to 8 bit RGB colour TIFF images. The Fiji software was used to label and cut out the sections of interest (∼28 consecutive SN sections within the caudo-rostral axis, bilaterally), and the ROI was identified and strictly marked according to typical anatomical landmarks and the cartesian coordinates according to the mouse brain atlas for each section, similar as described for IHC/DAB-stained neuron counting (Fu et al, 2012; Paxinos & Keith B. J. Franklin, 2007). Exemplary ROI marking is depicted in **Figure 7a** and **Figure S5a**. Processed images were uploaded on the Wolution platform and DLAP-5 (Wolution GmBH & Co. HG) was used for analysis. The algorithm automatically identified cell bodies and nucleus of both TH-(red channel) and DAPI-positive (blue channel) cells within the ROI and directly quantified the mean IF signal intensity per area for TH, DAPI and the target genes (green channel, DAT; magenta channel, CB, D2, and Aldh1A1) in the cell body and nucleus. Background (BG) regions (for normalization of signal intensities) were taken from regions outside the ROI by application of individual IF intensity thresholds for each RGB channel (as described in post-training procedures). All identified SN cells were manually checked for the correct and clear identification of full cell bodies and nuclei. If cell bodies were not completely separated from each other, or if they were cropped, fragmented, very small, or did not show a full nucleus, these cells were excluded from further semi-quantitative analysis (∼30% of identified TH-positive SN neurons, **Figure S5a**, **Table S12**). This strategy was used, as our aim was analysis of TH and DAT co-expression and signal-quantification. For absolute quantification of all TH-positive (or DAT-positive) neurons, a count of neuronal nuclei is required (and possible), as described for DLAP-3/4.

Raw signal intensities of each cell were normalised to the determined BG signals from the respective section. The BG signals determined from each analysed section were then normalised to the mean BG-value for each animal, and outliers were removed. The threshold for defining DAT-negative SN neurons was manually determined, and set as the mean normalised BG value for each mouse, plus 1.5 to 2.5 times SD of the DAT relative fluorescence (RF) signal (depending on the different signal intensities of the different mouse cohorts; **Figure S5c**). Similarly, a threshold of mean normalised BG value for each mouse plus 1 time SD for D2, plus 4 times SD for CB, and plus 5 times SD for Aldh1A1 was used (**Figure S8, Table S15**).

For the generation of anatomical 2D/3D maps, the R-computational environment (Team, 2014) version 4.2.1 with the Tidyverse package (Wickham et al, 2019), Plotly [https://plot.ly/] and RMarkdown (Holmes et al, 2021) was used, and coordinates as well as DAT and TH signal intensities were systematically adjusted in order to plot the different cohorts and animals into one graph. 2D-Maps were generated by using and adjusting the x and y coordinates from each analysed TH-positive neuron, automatically provided by the DLAP-5 algorithm. The respective z-coordinates were derived according to the thickness of the individual consecutive back-to-back brain sections (30 µm, bregma: caudal to rostral; -3.9 to -2.7). Alternatively, as DLAP processes single images within a part of a z-stack of images and processes them slice by slice, the output-files of the algorithms are easily amended, so the user can reassemble the z-stack of segmentations afterwards, to analyze data in three-dimensions. Training a 3-dimensional CNN is also possible with the DLAP platform, but was not necessary here. For normalization of anatomical coordinates of individual animals, we applied animal and SN hemisphere dependent z-score normalization of x and y coordinates using the “scale ()” function in R. Thereby, the center of each SN hemisphere (defined as the midpoint of the coordinate scale for all identified SN DA neurons on this hemisphere) was set as 0,0-value for both axes, a commonly used strategy (Curtis et al, 2016). We also utilized these adjusted x and y coordinates for defining the lateral SN region as starting at >1.5 scaled x-axis units lateral (i.e. >377.8 µm) from each SN hemisphere-center (0,0; **Figure S6, Table S17**). For generation of 2D and 3D gradient plots, we used the “scale ()” function to generate scaled DAT and TH RF values: similar as for the anatomical coordinates, the mean of the DAT and the TH RF intensity of all anaysed TH-positive neurons was set as 0-value and the deviation from the mean was plotted for each individual neuron, using a colour gradient. For better gradient-visualization, we excluded the highest values from the gradient-plots of (0.3% of all neurons for DAT, 0.005% for TH).

For relative quantification of fluorescence-signal intensities in cellular compartments with DLAP-6 algorithm, OBF images were converted to 8 bit RGB colour TIFF images using the Fiji software. For manual analysis, images were further processed using Fiji. First, the “polygon selection” tool was used to manually delineate a continuous membrane ROI around the cell based on the Kv4.3 IF signal (green channel). The cytoplasm was marked according to the immunofluorescence signal for the cytoplasmic TH (red channel), and nucleus was marked according to the DAPI signal (blue channel). In the next step, the “measure” function (under the “analyze” toolbar) was used to calculate the mean, RF signal intensity for each compartment, already normalised to the respective area. For automated analysis, the same high resolution RGB colour TIFF images were uploaded on the Wolution platform and analysed with DLAP-6. Each identified TH-positive neuron was segmented into “cytoplasm”, “membrane” and “nucleus”, based on the cytoplasmic TH and nuclear DAPI fluorescence signal (after training, the plasma-membrane was reliably identified without a third membrane-marker). BG signal intensities were determined from images acquired outside the region of analysis, using individual RGB channel thresholds, and were used to normalise signal intensities for both, manual and automated analyses. All images were manually checked for the correct identification of membrane, cytoplasm and nuclear compartments similar as performed for analysis with DLAP-5 and ∼90 % of cells were used for further quantitative analysis. In case of non-continuous membrane compartment identification, the weighted mean (area) of RF signal intensity in all identified membrane ROIs from each cell was used for analysis.

### Statistics and software

Data analysis and graphical illustrations were performed using GraphPad Prism 9 (GraphPad Software, Inc.), Adobe Illustrator CC2015.3 (Adobe Systems Software), Wolution (https://wolution.com/), and Fiji (https://imagej.net/Fiji) software. Statistical tests were performed with GraphPad Prism 9. Anatomical maps were plotted using R computational environment (Team, 2014) version 4.2.1 with Tidyverse package (Wickham et al, 2019), Plotly [https://plot.ly/], and RMarkdown (Holmes et al, 2021).

Normal distribution was tested with D’Agostino-Pearson omnibus normality test. Correlations were performed using Pearson correlation test, proportionality constant was calculated from the ratio between two directly corresponding values. In graphs, data are given as boxplots showing median and whiskers representing 10^th^ -90^th^ percentile. For analyzing one parameter non-parametric Mann-Whitney U test and Kruskal-Wallis test, with post hoc Dunn’s multiple comparison was used. For comparison between categorical %, Fisher’s exact tests and Chi-squared tests were used, as indicated. Testing for two independent parameters was performed via two-way ANOVA with Tukey’s multiple comparison test. ROUT outlier test (Q=1) was used to remove outliers. Statistical significances are indicated as ns > 0.05, *p* < 0.05 (*), *p* < 0.01 (**), *p* < 0.001 (***) and *p* < 0.0001 (****).

## Supplementary Figures

**Figure S1:**
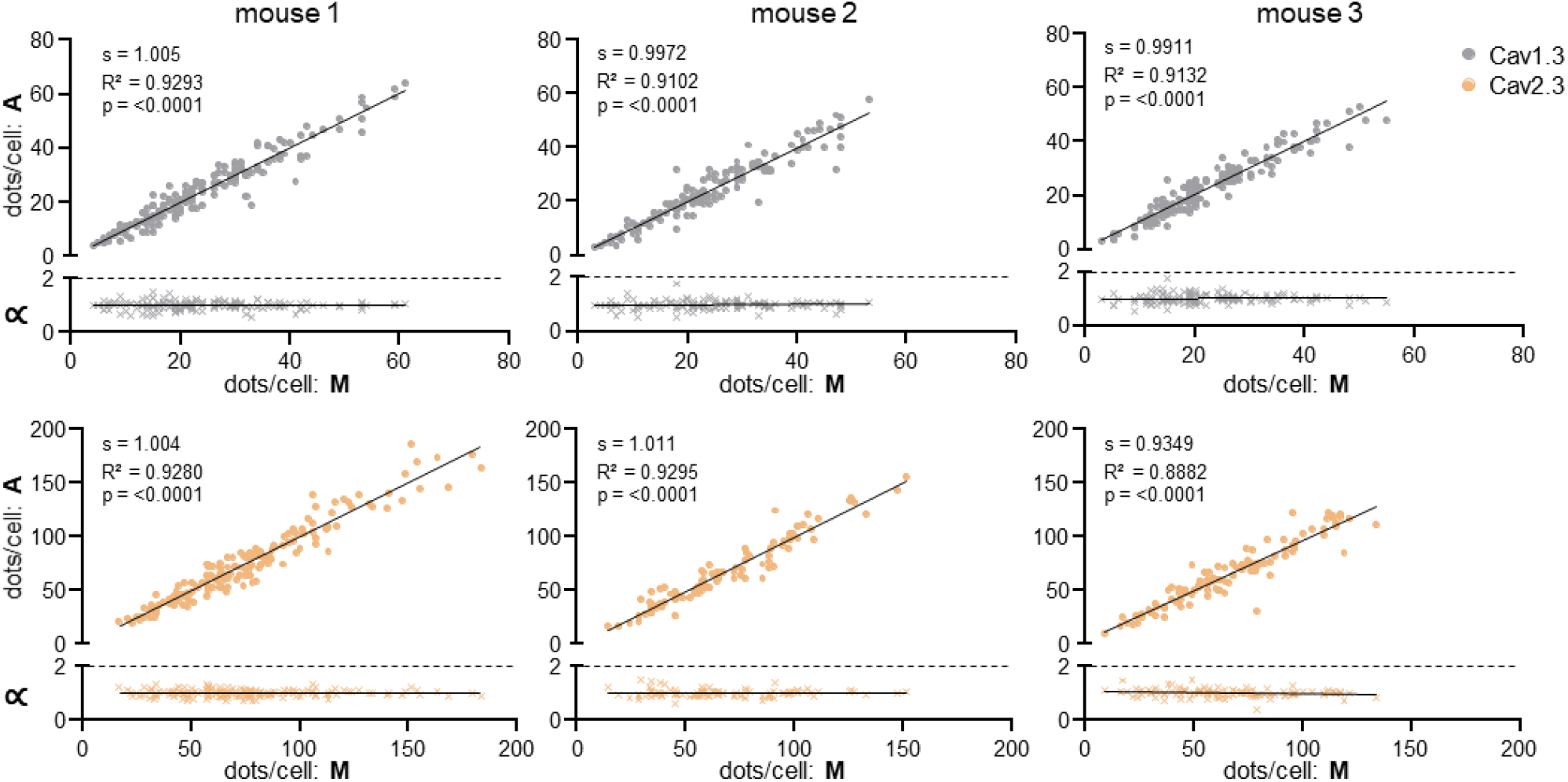
Comparison between manual and automated (DLAP-1) RNAscope-derived image analysis of mouse midbrain sections. Upper: single neuron correlations of Cav1.3 (grey) and Cav2.3 (light brown) mRNA-derived dot counts/TH-positive SN neuron between manual (M) and automated (A) analysis according to Pearson correlation test separately for each analysed mouse. Lower: corresponding proportionality constants ∝, calculated from the manual and automated dot count ratios.

**Figure S2:**
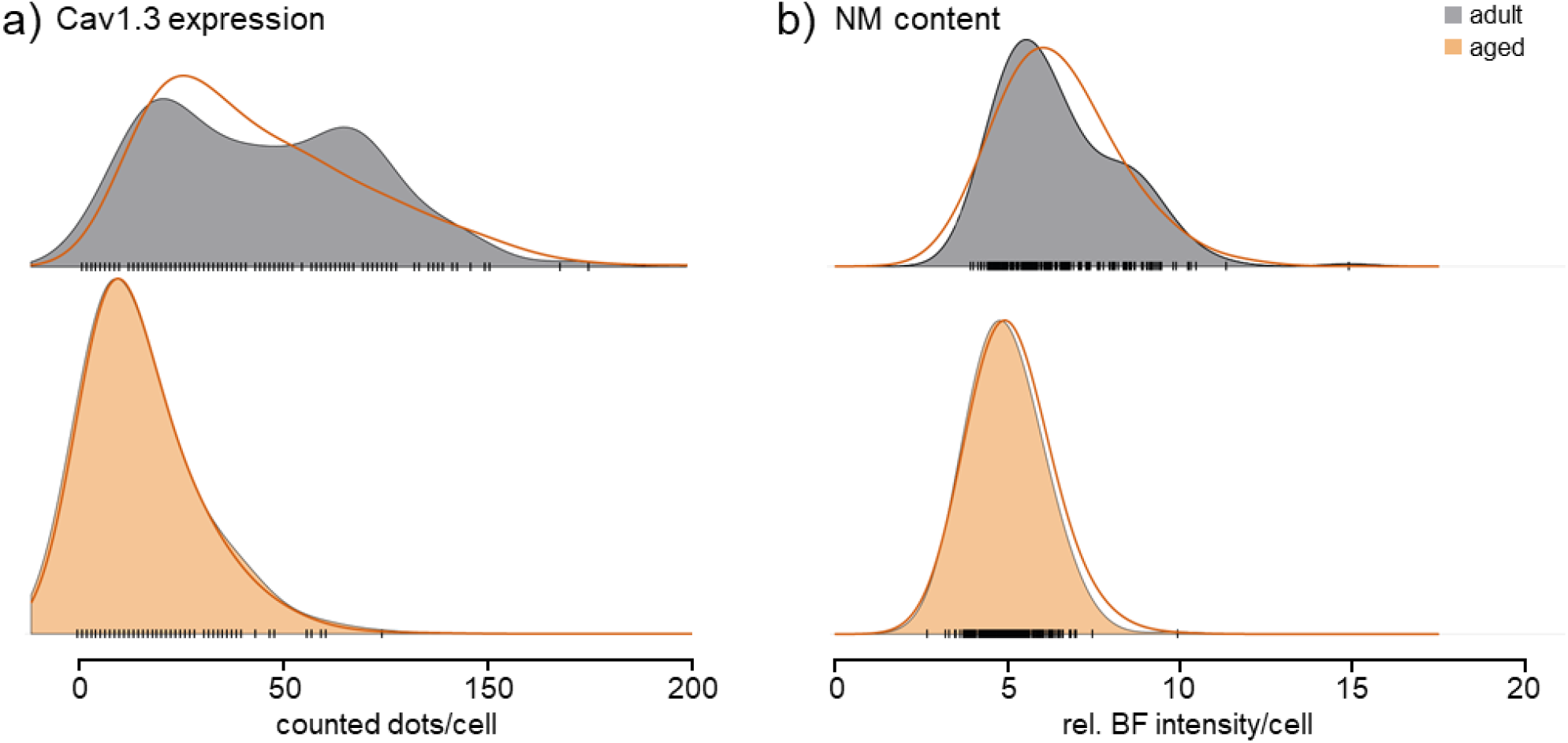
Posterior-predictive control of human RNAscope data model-fits. Cav1.3 expression (a) and NM content (BF light transmission, b) data is represented by vertical black lines and the colour-filled kernel densities for adult and aged brains, as indicated. Red continuous lines display the kernel distribution of data reproduced from the fitted model. Note the bimodal distribution in the group of adult brains in Cav1.3 expression and BF light transmission. The model does attribute for the wider base of the distribution of the number of detected Cav1.3 mRNA molecules (two components mixture of binomials) and BF light transmission (two components of beta), but does not represent the mixture proportion.

**Figure S3:**
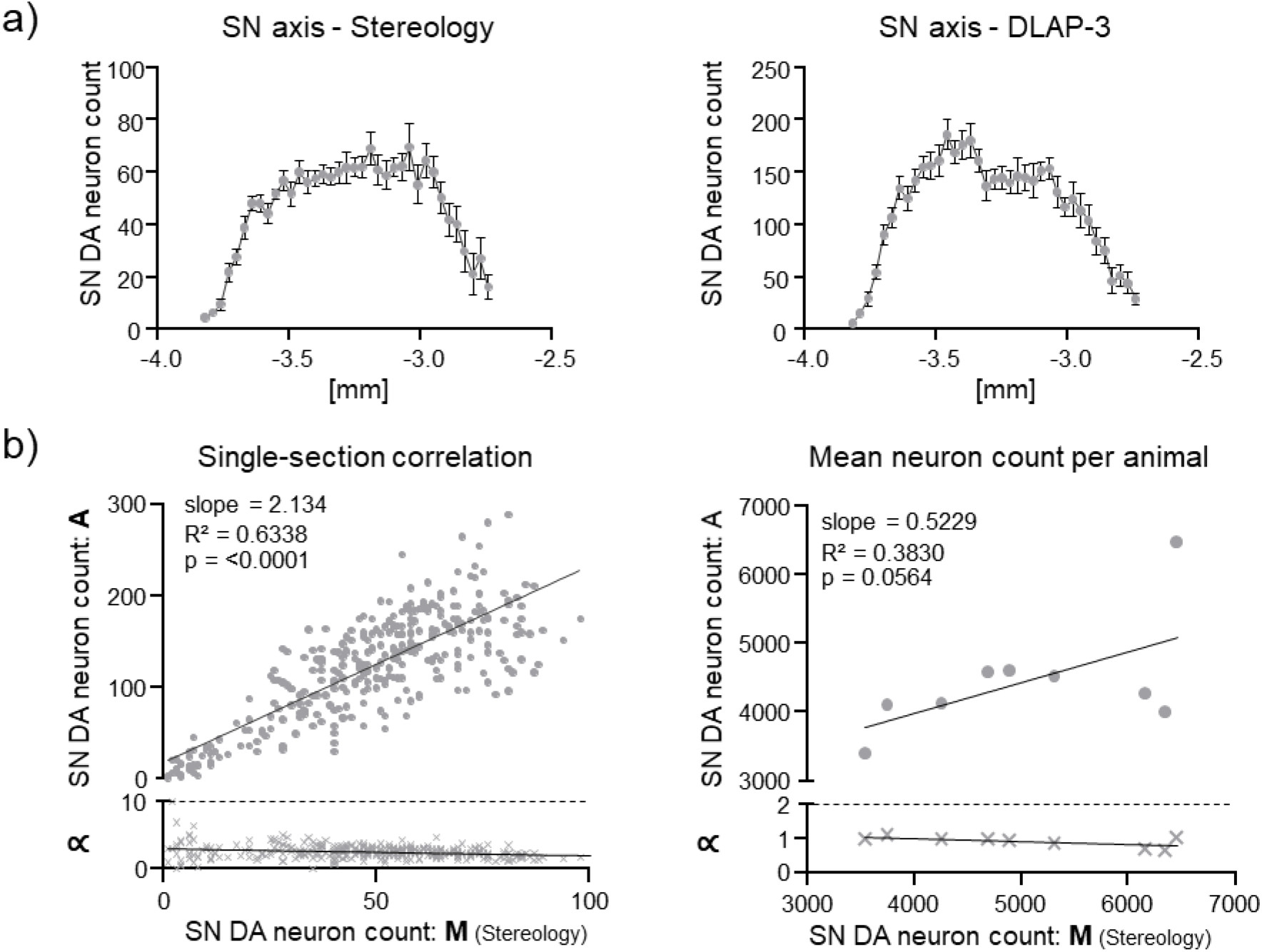
Comparison between unbiased stereology and automated (DLAP-3) IHC-derived image analysis of juvenile mouse midbrain sections. a) Mean SN DA neuron counts for all animals (N = 10) in all analysed sections along the caudal-rostral axis quantified via unbiased stereology (extrapolated via the optical fractionator method, M) or automatically (A) via DLAP-3. Data are given as mean ± SEM. b) Upper: correlations of individual SN DA neuron counts/section and animal (left) and mean SN DA neuron counts/animal (right) between unbiased stereology and automated analysis according to Pearson correlation test. Lower: corresponding proportionality constants ∝, calculated from the manual and automated cell countratios.

**Figure S4:**
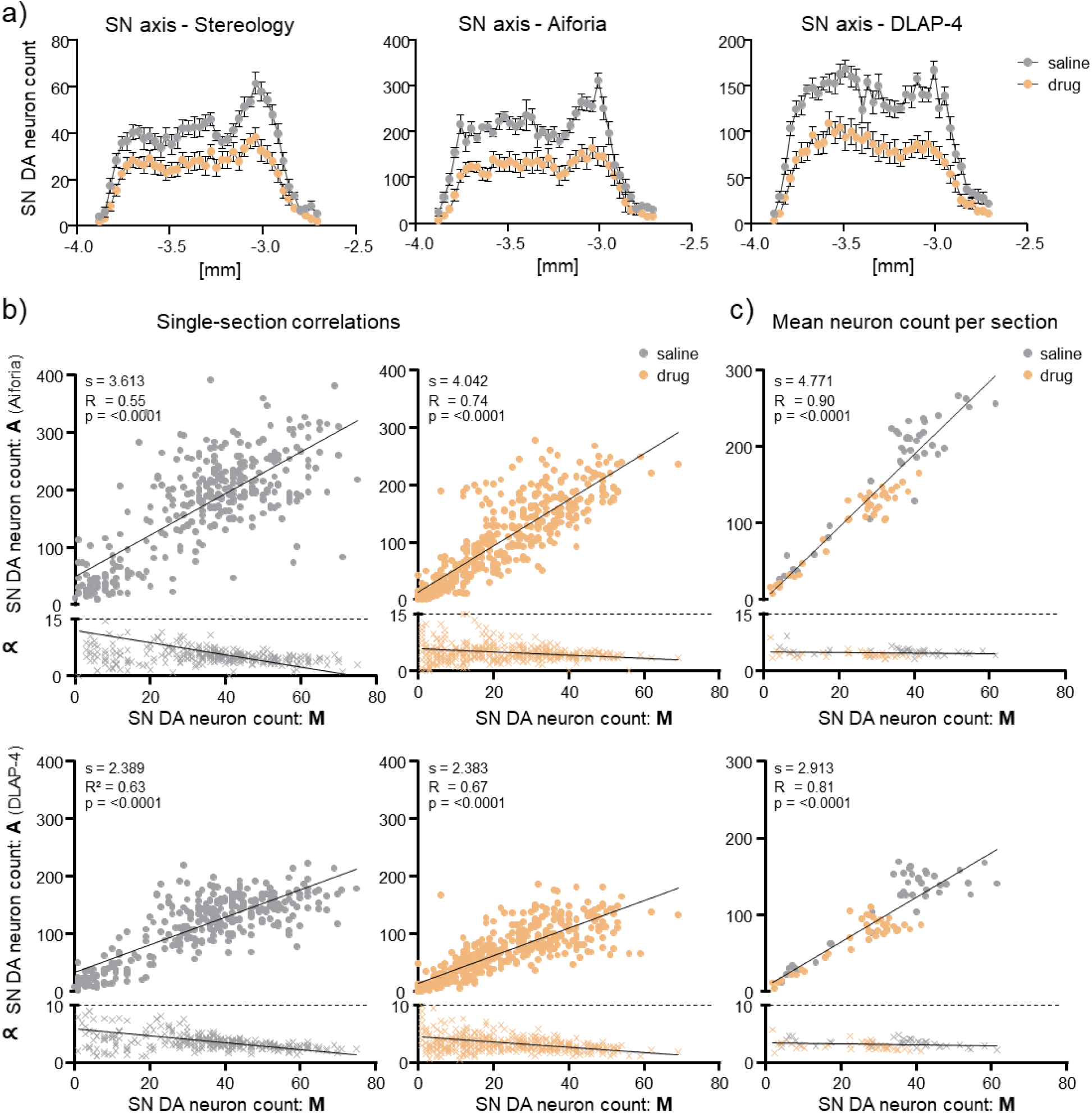
Comparison between unbiased stereology and automated (DLAP-4) IHC-derived image analysis of adult mouse midbrain sections. a) Mean SN DA neurons for all animals (saline: N = 9; neurodegenerative drug: N = 12) in all analysed sections along the caudal-rostral axis, quantified via unbiased stereology (extrapolated via the optical fractionator method, M) or automatically (A) via an algorithm provided by Aiforia, or via DLAP-4, as indicated. Data are given as mean ± SEM. b/c) Upper: correlations of individual SN DA neuron counts/section and animal (b) and mean animal counts/section along caudo-rostral axis (calculated from the individual values for each animal, c) between unbiased stereology and automated analysis (Aiforia, or DLAP-4, as indicated) according to Pearson correlation test. Lower: corresponding proportionality constants ∝, calculated from the manual and automated cell count ratios. Stereology data and Aiforia data modified from *Benkert et al., Nat. Commun. 2019*.

**Figure S5:**
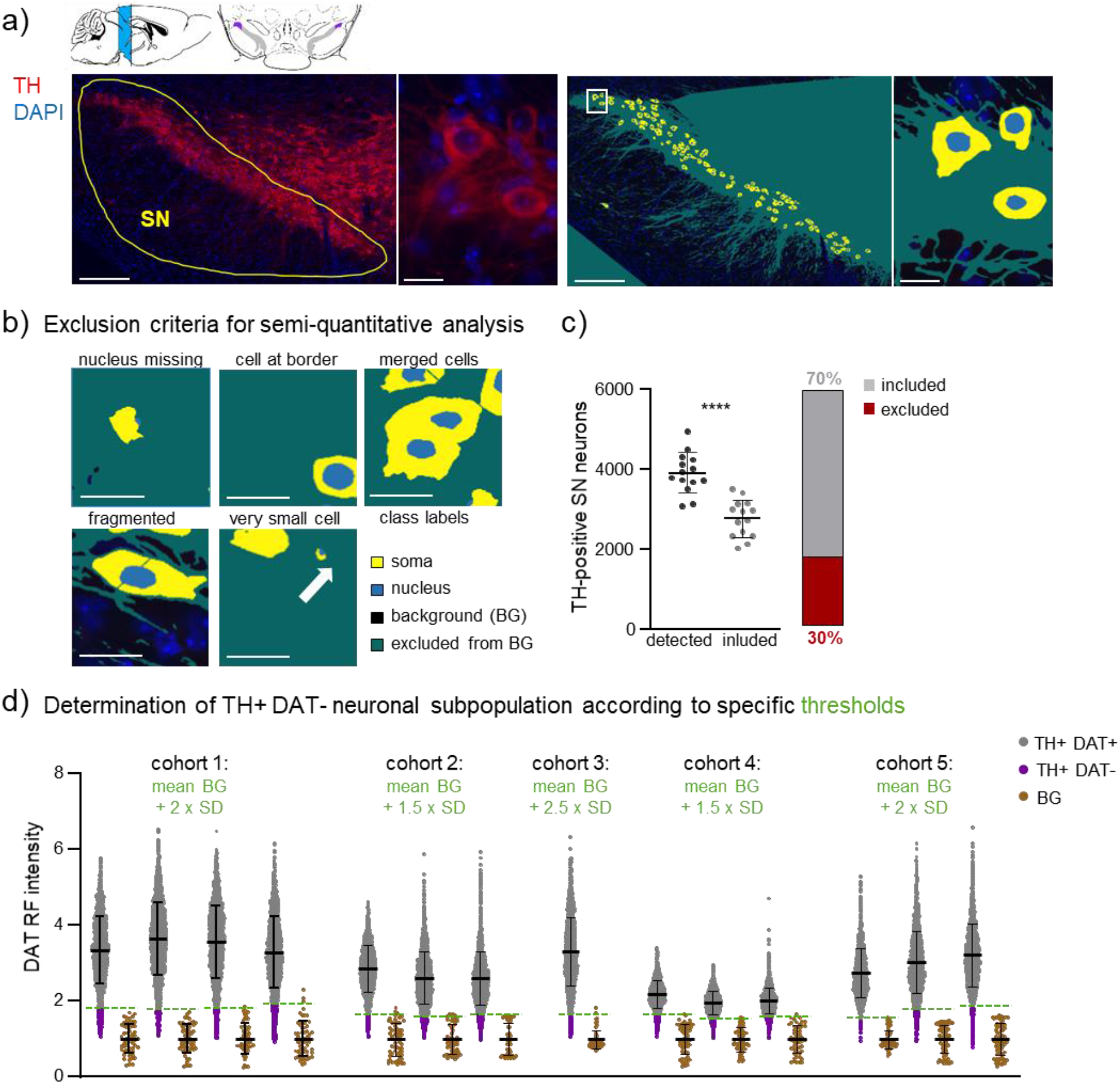
Relative DAT-signal quantification in TH-positive SN neurons from mouse midbrain sections with DLAP-5. a) Upper: exemplary sagittal and coronal mouse brain sections, modified from (Paxinos & Keith B. J. Franklin, 2007), illustrating the analysed caudo-rostral extent of the SN (bregma: -3.9 to -2.7, sagittal, blue), and its lateral parts in coronal sections (lateral SN: violet, non-lateral SN grey, compare Figure 7, S5). SN neurons after IF for TH (red). Nuclei are marked by DAPI (blue), the SN is delineated in yellow, the white box in the lateral SN indicates the position of DAT-negative neurons. Right: enlarged image of the white-boxed area from the left, output image after automated recognition of TH labelled cell bodies (yellow area) and corresponding nuclei (blue area) with DLAP-5. Scale bars: 200 μm (left) and 20 µm (right). b) Examples of detected TH-positive neurons, where cell bodies were cropped (at the border), merged/not clearly separated from each other, fragmented, very small, or did not show a full nucleus. Such cells were excluded from relative quantification analysis. Scale bars: 20 µm. c) Left: numbers of TH-positive SN neurons/mouse, determined via DLAP-5. Shown are the numbers of all detected TH-positive SN neurons without any exclusion, according to the criteria in b), as well as the neuron numbers after exclusion-analysis, only displaying those TH-positive neurons were relative DAT-intensities were determined. Significant difference according to Mann-Whitney test (****: p < 0.0001). Right: percentage of excluded neurons. d) Plotted are the individual relative DAT fluorescence intensities (RF) for all included TH-positive SN neurons and for all backgrounds (BG, derived separately for each image, normalised to the mean BG in each animal), for each individual mouse, quantified via DLAP-5. The DAT-intensity thresholds for positive cells were determined individually for each mouse, according to the mean and the SD of the BG-signals, indicated as dashed green lines. Data are given as scatterplots and mean ± SD for all analysed mice (N = 14). All data are detailed in Table S13 and S14.

**Figure S6:**
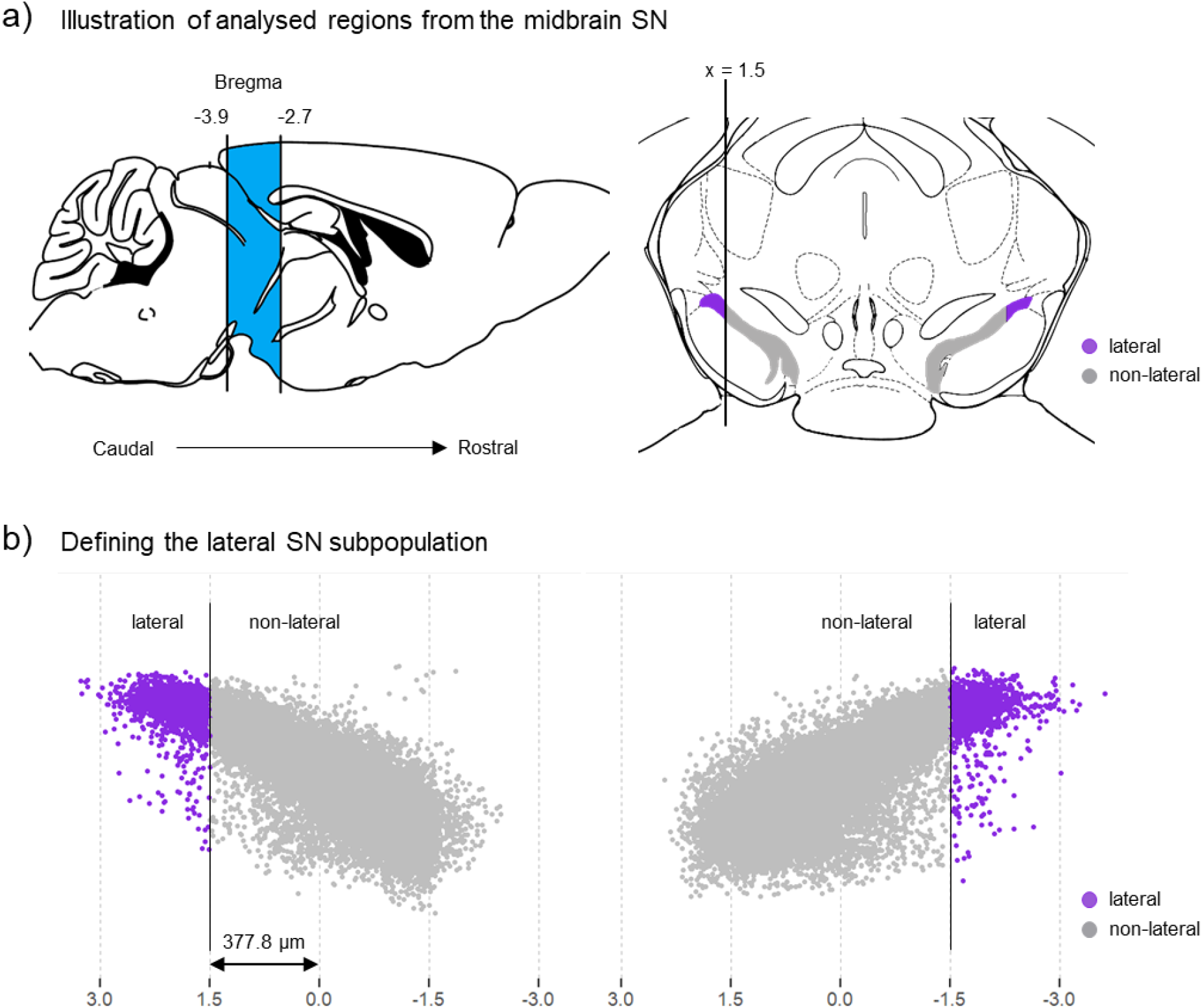
Definition of lateral and non-lateral SN-parts in coronal mouse brain sections, according to scaled x,y-coordinates. a) Sagittal (left) and coronal (right) mouse brain sections, modified from (Paxinos & Keith B. J. Franklin, 2007), illustrating the analysed caudo-rostral extent of the SN (bregma: -3.9 to -2.7, sagittal, blue), and its lateral parts in coronal sections (defined as >1.5 scaled x-units (>377.8 µm) lateral from each SN hemisphere-center; lateral SN: violet, non-lateral SN grey). Compare Figures 7-9. b) Plotted (here and in Figures 7-9) are the scaled x,y-coordinates for all TH-positive SN neurons that were further analysed for relative DAT signal intensities, from all analysed animals (N = 14). The lateral (violet) and the non-lateral (grey) SN are defined according to their scaled x,y-coordinates (determined via DLAP-5) and their distance (>1.5 x-units lateral) from the respective SN hemisphere-center (defined as 0.0 coordinates), visualized here by normalise and the vertical lines at 1.5/-1.5 (lateral SN: left, n = 1749; right, n = 1547 TH-positive neurons; non-lateral: left, n = 17693; right, n = 17515).

**Figure S7:**
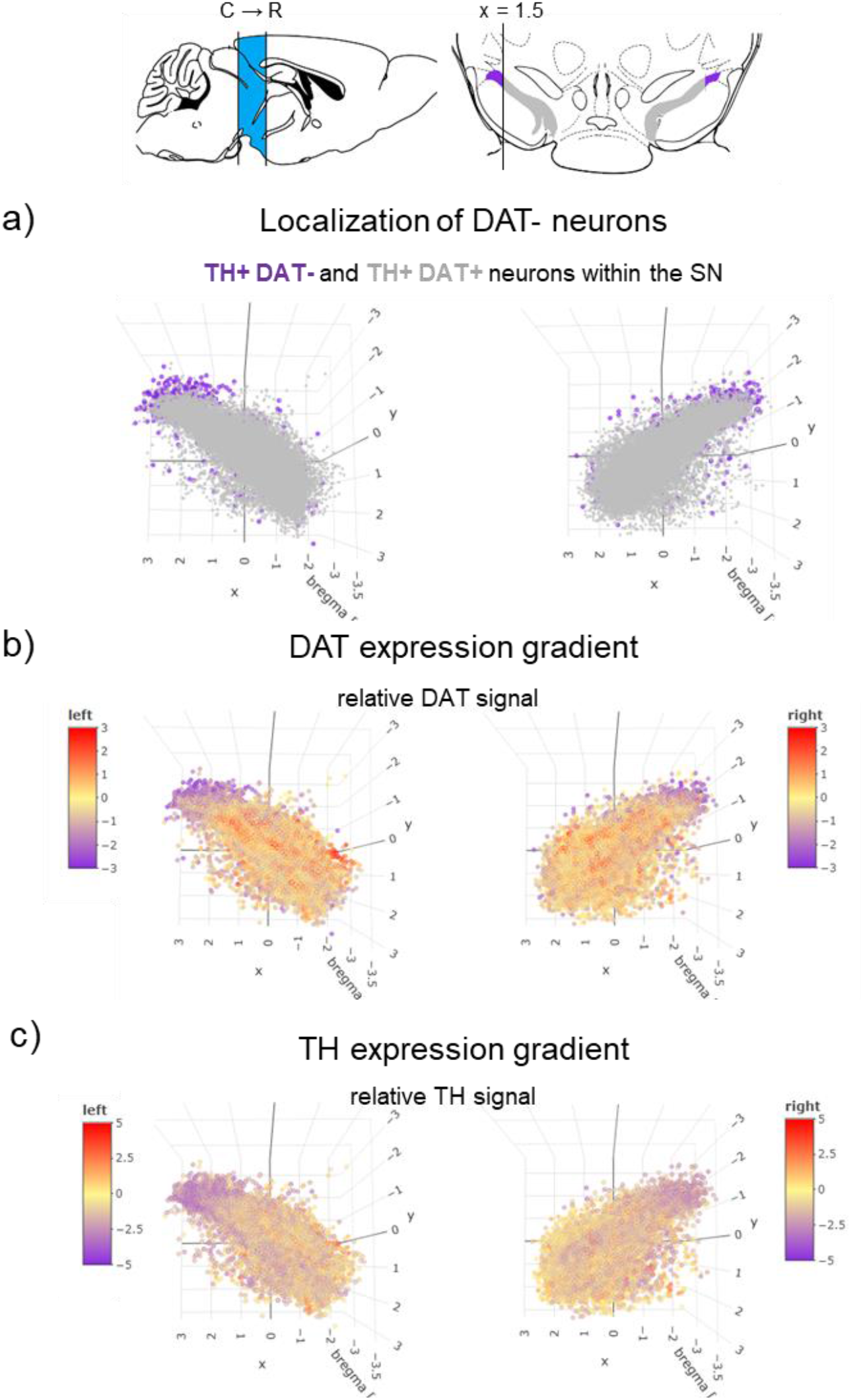
Thumbnail of the respective online interactive 3D-figure (HTML-file), plotting the anatomical location of all further analysed TH-positive SN neurons. a) Upper: sagittal (left) and coronal (right) mouse brain sections, modified from (Paxinos & Keith B. J. Franklin, 2007), illustrating the analysed caudo-rostral extent of the SN (bregma: -3.9 to -2.7, sagittal, blue), and its lateral parts in coronal sections (defined as >1.5 scaled x-units (>377.8 µm) lateral from each SN hemisphere-center (0,0); lateral SN: violet, non-lateral SN: grey, as in Figures 7-9, S6). Lower: plotted are the individual TH-positive SN neurons for all analysed animals (n = 38504, N = 14, as in Figure 7-9, S5), according to their scaled x,y,z-coordinates. The resulting anatomical 3D-representations display the medio-lateral distribution of the TH-positive DAT-negative SN neurons (violet) over the full rostro-caudal extent for all analysed mice. b/c) colour coded are the scaled relative DAT (b) and TH (c) signal intensities, of the individual TH-positive neurons from a), plotted according to their scaled x,y,z-coordinates. Colour coding for each neuron according to its individual deviation from the scaled mean signal-intensity (0.0) for each animal. The corresponding 2D maps are given in Figures 7-9. To access the interactive HTML-file, download it at. https://cloud.physio.uni-ulm.de/s/jo28kjk2edjTf8r

**Figure S8:**
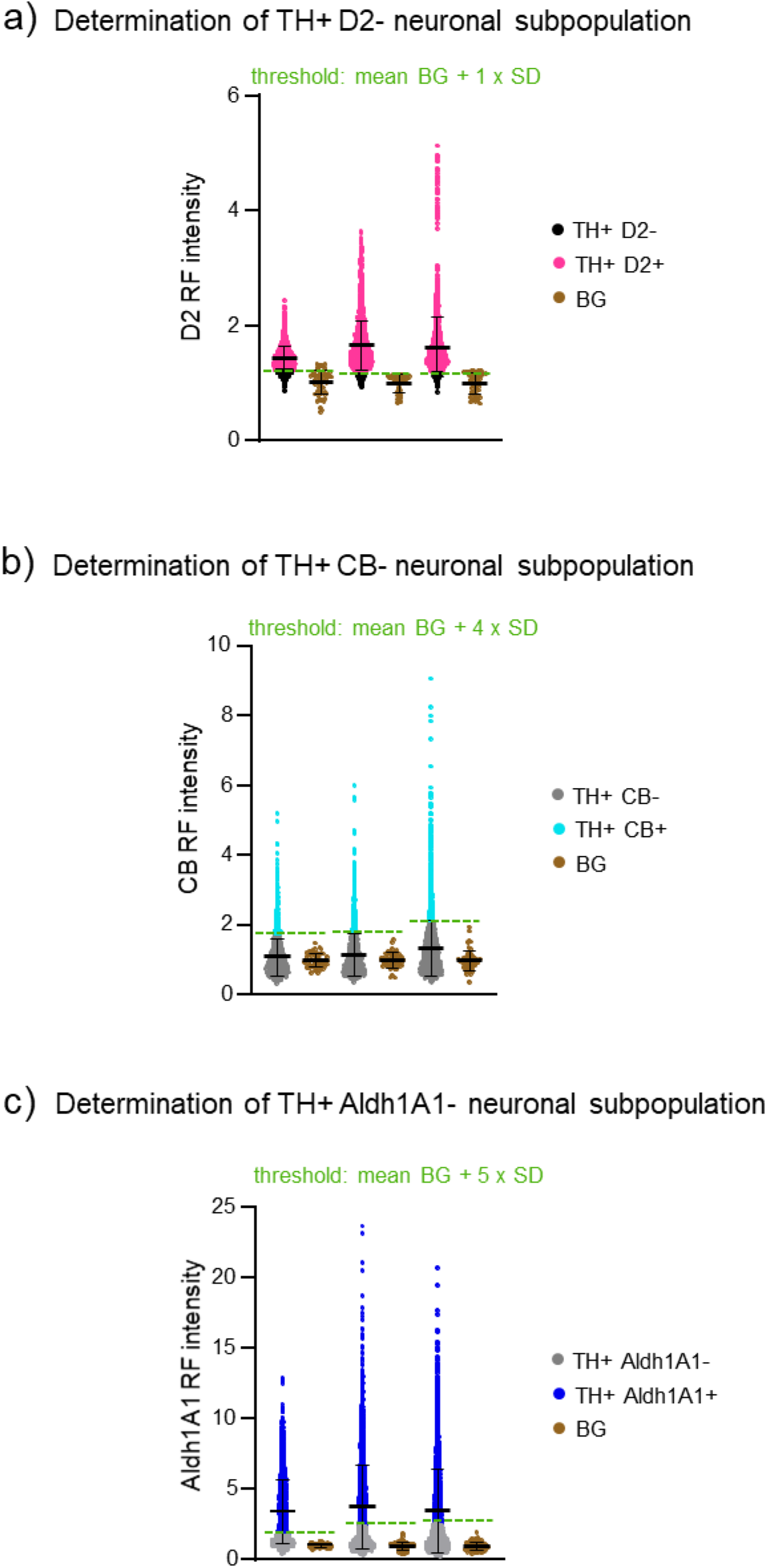
Relative D2, CB, and Aldh1A1-signal quantification in TH-positive SN neurons from mouse midbrain sections with DLAP-5. Plotted are the individual relative fluorescence intensities (RF) for a) D2, b) CB, c) Aldh1A1, for all included TH-positive SN neurons and for all backgrounds (BG, derived separately for each image, normalised to the mean BG in each animal), for each individual mouse, quantified via DLAP-5. The respective-intensity thresholds for D2-/CB-/Aldh1A1-positive cells were determined individually for each mouse, according to the mean and the SD of the BG-signals, and indicated as dashed green line. Data are given as scatterplots and mean ± SD for all analysed mice (N = 3; TH+ n = 2023-3417, BG n = 51-72) and detailed in Table S14.

**Figure S9:**
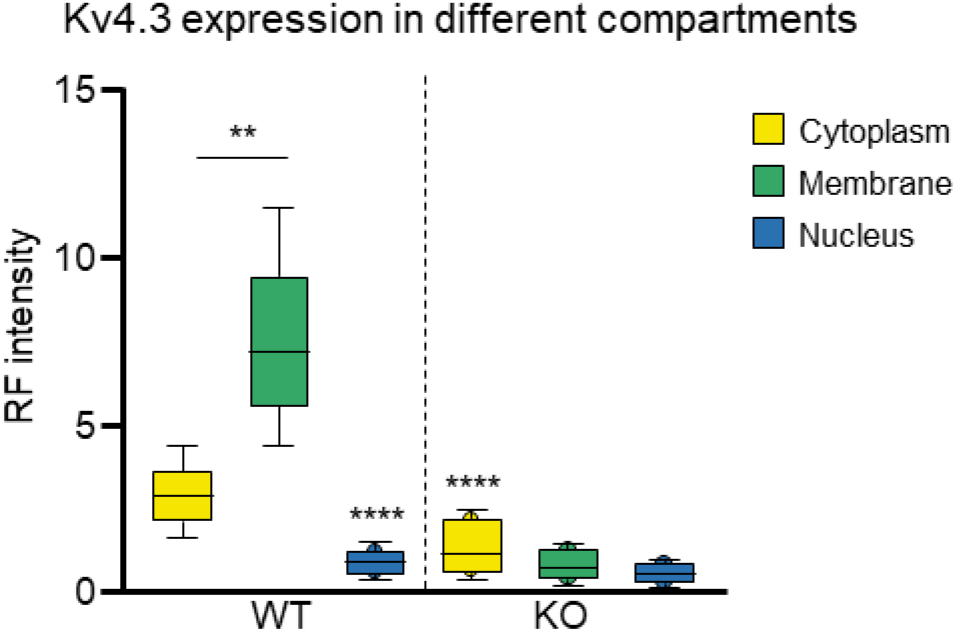
Automated (DLAP-6) IF-derived image analysis for cellular compartments of TH-positive SN neurons from WT and Kv4.3 KO mice. Kv4.3 relative fluorescence (RF) intensity/neuron in different sub-cellular compartments, quantified in WT and Kv4.3 KO mice, as indicated. Data are given as boxplots (median, 10-90 percentile) for all analysed neurons (WT: n = 179, Kv4.3 KO: n = 200). Significant differences according to Kruskal-Wallis with Dunn’s multiple comparison tests (**: p<0.01, ****: p < 0.0001).

## Supplementary tables

**Table S1:** Comparison of FCN and Deeplab3 network performances in distinct tasks. Performances were determined by comparing analysis results the test data to the ground truth data (provided by manual labeling). The following quality measures were assessed separately for each class: pixel error rate, rates for true positive (TP), true negative (TN), false positive (FP), false negative (FN) detection as well as specificity (spec.) [defined as TN/(TN+FP)] and sensitivity (sens.) [defined as TP/(TP+FN)].

| D-LAP algorithm |  | Error [%] | class name | TP [%] | TN [%] | FP [%] | FN [%] | Spec [%] | Sens [%] |
| --- | --- | --- | --- | --- | --- | --- | --- | --- | --- |
| 1 | FCN | 5.08 | mRNA | 0.3 | 98.9 | 0.7 | 0.0 | 99.3 | 92.5 |
|  |  |  | neuron | 10.3 | 84.7 | 4.1 | 0.8 | 95.3 | 92.7 |
|  | DeepLab3 | 4.39 | mRNA | 0.0 | 99.7 | 0.0 | 0.3 | 100.0 | 0.0 |
|  |  |  | neuron | 11.0 | 84.6 | 4.2 | 0.2 | 95.2 | 98.3 |
| 2 (human) | FCN | 4.13 | mRNA | 0.2 | 98.9 | 0.9 | 0.0 | 99.1 | 94.7 |
|  |  |  | neuron | 5.2 | 91.0 | 2.5 | 1.4 | 97.3 | 80.8 |
|  | DeepLab3 | 2.49 | mRNA | 0.0 | 99.8 | 0.0 | 0.2 | 100.0 | 0.0 |
|  |  |  | neuron | 6.1 | 91.4 | 2.1 | 0.4 | 97.8 | 95.6 |
| 3 | FCN | 0.64 | nucleus | 0.0 | 99.4 | 0.0 | 0.6 | 100.0 | 1.7 |
|  | DeepLab3 | 0.61 | nucleus | 0.5 | 98.9 | 0.5 | 0.1 | 99.5 | 85.5 |
| 4 | FCN | 0.31 | nucleus | 0.2 | 99.5 | 0.3 | 0.0 | 99.7 | 46.7 |
|  | DeepLab3 | 0.26 | nucleus | 0.1 | 99.7 | 0.2 | 0.0 | 99.8 | 59.8 |
| 5 | FCN | 7.08 | nucleus | 1.0 | 97.3 | 1.5 | 0.2 | 98.4 | 80.9 |
|  |  |  | cytoplasm | 3.7 | 90.8 | 4.9 | 0.7 | 94.8 | 83.8 |
|  | DeepLab3 | 3.3 | nucleus | 0.9 | 98.5 | 0.3 | 0.3 | 99.7 | 74.5 |
|  |  |  | cytoplasm | 4.1 | 92.9 | 2.2 | 0.8 | 97.7 | 83.7 |
| 6 | FCN | 8.34 | nucleus | 9.4 | 87.9 | 1.2 | 1.5 | 98.7 | 87.2 |
|  |  |  | cytoplasm | 12.4 | 82.8 | 4.1 | 0.8 | 95.3 | 93.4 |
|  |  |  | membrane | 2.3 | 94.2 | 1.7 | 1.7 | 98.2 | 58.9 |
|  | DeepLab3 | 5.49 | nucleus | 3.0 | 93.5 | 2.5 | 1.0 | 97.4 | 75.6 |
|  |  |  | cytoplasm | 9.7 | 89.0 | 0.8 | 0.5 | 99.1 | 95.0 |
|  |  |  | membrane | 11.7 | 85.2 | 2.2 | 0.9 | 97.5 | 92.4 |

**Table S2:** Details of RNAscope probes. RNAscope target probes were obtained from the Advanced Cell Diagnostics (ACD) library. All probes detected but did not discriminate between known splice variants of the respective target genes. Abbreviations: NCBI: National Center for Biotechnology Information.

| Species | Gene | ACD Cat No. | Assay target region according to NCBI Genbank gene accession no. (NM) |
| --- | --- | --- | --- |
| mouse | Tyrosine hydroxylase (*Th*) | 317621 | 483 – 1603 of NM_009377.1 (exon 3-13) |
|  | Cav1.3 (*Cacna1d*) | 502591 | 522 – 1994 of NM_001302637.1 (exon 1-10) |
|  | Cav2.3 (*Cacna1e*) | 449211 | 2319 – 3317 of NM_009782.3 (exon 20-22) |
| human | Tyrosine hydroxylase (*TH*) | 441651 | 596 – 1681 of NM_199292.2 (exon 4-14) |
|  | Cav1.3 (*CACNA1D*) | 502591 | 522 – 1994 of NM_001302637.1 (exon 1-10) |

**Table S3:**
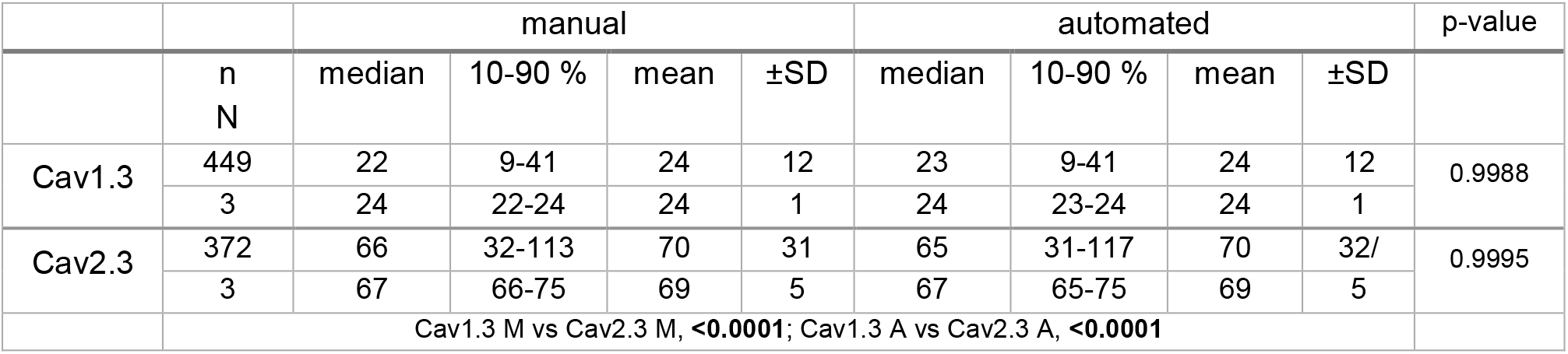
Mean mRNA molecule numbers in individual mouse SN DA neurons, determined manually and via algorithm DLAP-1. Data and statistics for graphs in Fig. 3b. Number of total analysed neurons is given by n, N represents the number of analysed mice. P-values according to Two-way ANOVA followed by Tukey’s multiple comparison (significant values in bold).

**Table S4:** Single cell correlation analyses between manually and DLAP algorithm 1 determined results. Data and statistics for graphs in Fig. 3c and S1a, according to Pearson correlation (slope of the linear regression with its own 95% confidence interval).

|  | Cav1.3 | Cav2.3 |
| --- | --- | --- |
| all mice (N = 3) |  |  |
| Pearson r (95% confidence interval) | 0.959 (0.950-0.966) | 0.959 (0.950-0.967) |
| R squared | 0.919 | 0.920 |
| p-value | <0.0001 | <0.0001 |
| No. of XY pairs | 444 | 372 |
| Slope (95% confidence interval) | 0.998 (0.971-1.026) | 0.989 (0.960-1.020) |
| Proportionality constant $\alpha$ | 1.01 ± 0.16 | 1.00 ± 0.14 |
| mouse 1 |  |  |
| Pearson r (95% confidence interval) | 0.964 (0.952-0.973) | 0.963 (0.951-0.973) |
| R squared | 0.929 | 0.928 |
| p-value | <0.0001 | <0.0001 |
| No. of XY pairs | 168 | 167 |
| Slope (95% confidence interval) | 1.005 (0.962-1.047) | 1.004 (0.961-1.047) |
| Proportionality constant $\alpha$ | 1.01 ± 0.15 | 1.00 ± 0.13 |
| mouse 2 |  |  |
| Pearson r (95% confidence interval) | 0.954 (0.936-0.967) | 0.964 (0.947-0.976) |
| R squared | 0.910 | 0.930 |
| p-value | <0.0001 | <0.0001 |
| No. of XY pairs | 131 | 96 |
| Slope (95% confidence interval) | 0.997 (0.943-1.052) | 1.011 (0.954-1.068) |
| Proportionality constant $\alpha$ | 1.00 ± 0.17 | 1.00 ± 0.14 |
| mouse 3 |  |  |
| Pearson r (95% confidence interval) | 0.956 (0.939-0.968) | 0.943 (0.917-0.960) |
| R squared | 0.913 | 0.888 |
| p-value | <0.0001 | <0.0001 |
| No. of XY pairs | 145 | 109 |
| Slope (95% confidence interval) | 0.991 (0.941-1.042) | 0.935 (0.871-0.998) |
| Proportionality constant $\alpha$ | 1.02 ± 0.16 | 1.01 ± 0.15 |

**Table S5:** Details of human *post mortem* midbrain samples. Samples were provided by the German BrainBank. The mean age of the aged cohort was significantly higher compared to that of the adult cohort (** = 0.0012; t-test), the mean RIN was significantly lower (*0.038; t-test), PMI was not significantly different. Abbreviations: AD: Alzheimer’s disease; CERAD: Consortium to establish a registry for Alzheimer’s disease; f: female; LB: Lewy Body; m: male; NFT: neurofibrillary tangles; PD: Parkinson’s disease; PMI: *post mortem* interval, RIN: RNA integrity number.

| Cohort | Age, sex | PMI [h] | RIN | Braak AD/NFT, PD | CERA-stage | LB-pathology |
| --- | --- | --- | --- | --- | --- | --- |
| adult | 46, f | 27 | 8.5 | - , 0 | 0 | no |
|  | 34, m | 10 | 8.3 | - , 0 | 0 | no |
|  | 46, m | 19 | 8.4 | - , 0 | 0 | no |
| *mean±SD* | 42 ± 7** | 19 ± 9 | 8.4 ± 0.1* |  |  |  |
| aged | 76, m | 24 | 6.9 | I, - | A | no |
|  | 76, f | 11 | 6.4 | 0, - | 0 | no |
|  | 81, f | 18 | 4.0 | II, - | 0 | no |
| *mean±SD* | 78 ± 3 | 18 ± 7 | 5.8 ± 1.6 |  |  |  |

**Table S6:**
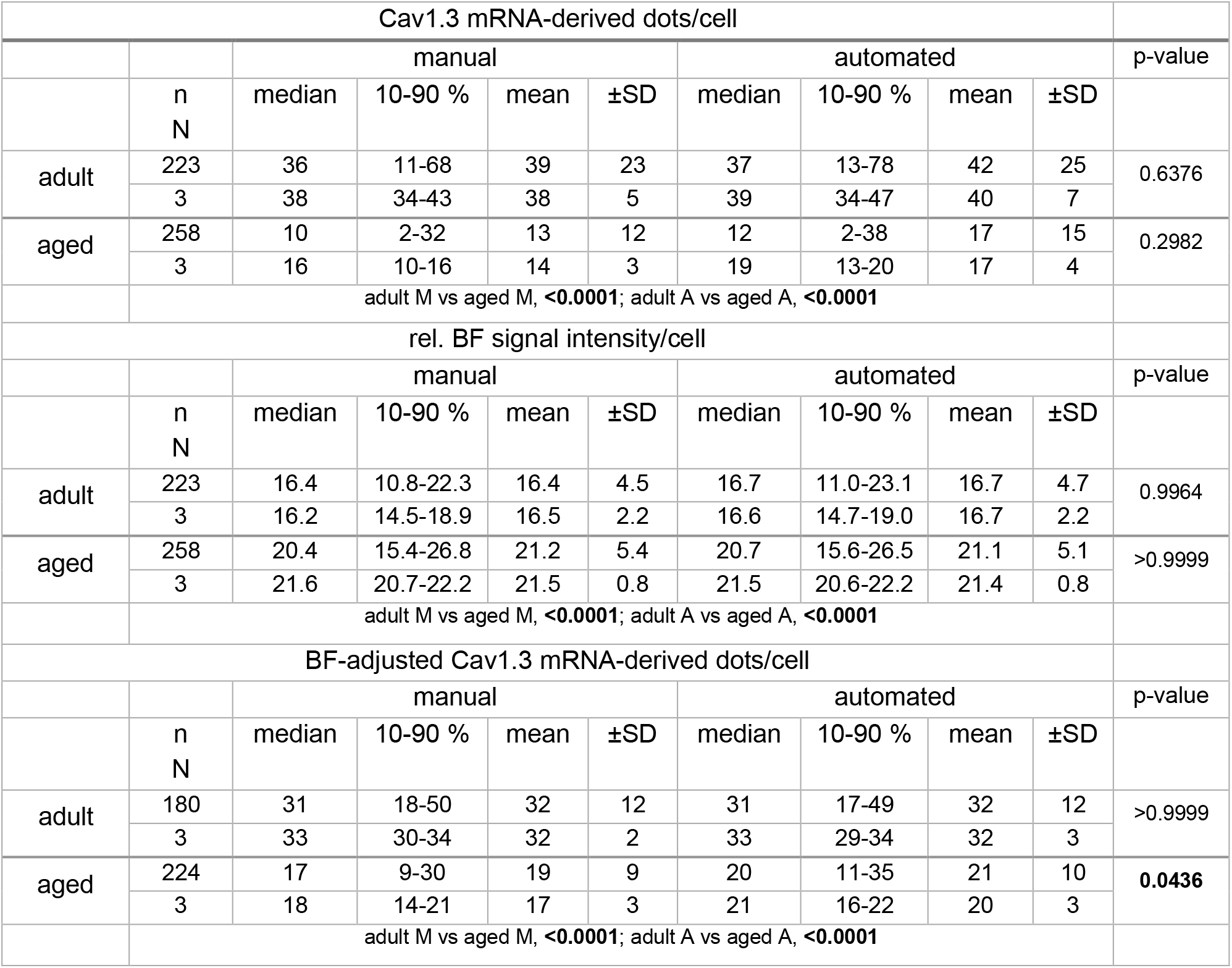
mRNA molecule numbers per human SN DA neuron, determined manually and via DLAP-2. Data and statistics for graphs in Fig. 4b and c. Number of total analysed neurons is given by n, N represents the number of analysed brain samples. P-values according to Two-way ANOVA followed by Tukey’s multiple comparison (significant values in bold).

**Table S7:** Single cell correlation analyses between manually and DLAP-2 determined results. Data and statistics for graphs in Fig. 4d and S1b and c, according to Pearson correlation (slope of the linear regression with its own 95% confidence interval).

|  | Cav1.3 mRNA<br>dot counts / cell | rel. BF intensity /<br>cell | Cav1.3 molecules,<br>NM/BF-adjusted<br>according to model |
| --- | --- | --- | --- |
| all human samples (N = 6) |  |  |  |
| Pearson r (95% confidence interval) | 0.964 (0.957-0.970) | 0.992 (0.990-0.993) | 0.897 (0.877-0.915) |
| R squared | 0.928 | 0.983 | 0.805 |
| p-value | <0.0001 | <0.0001 | <0.0001 |
| No. of XY pairs | 481 | 481 | 404 |
| Slope (95% confidence interval) | 1.034 (1.008-1.060) | 0.968 (0.957-0.979) | 0.867 (0.827-0.910) |
| Proportionality constant $\alpha$ | 1.28 ± 1.18 | 1.01 ± 0.03 | 1.10 ± 0.25 |
| adult samples (N = 3) |  |  |  |
| Pearson r (95% confidence interval) | 0.975 (0.967-0.980) | 0.996 (0.995-0.997) | 0.876 (0.837-0.906) |
| R squared | 0.950 | 0.992 | 0.767 |
| p-value | <0.0001 | <0.0001 | <0.0001 |
| No. of XY pairs | 223 | 223 | 180 |
| Slope (95% confidence interval) | 1.044 (1.012-1.076) | 1.033 (1.021-1.046) | 0.851 (0.781-0.920) |
| Proportionality constant $\alpha$ | 1.23 ± 1.51 | 1.01 ± 0.02 | 1.01 ± 0.18 |
| aged samples (N = 3) |  |  |  |
| Pearson r (95% confidence interval) | 0.890 (0.861-0.913) | 0.988 (0.984-0.990) | 0.872 (0.836-0.900) |
| R squared | 0.792 | 0.976 | 0.872 |
| p-value | <0.0001 | <0.0001 | <0.0001 |
| No. of XY pairs | 258 | 258 | 224 |
| Slope (95% confidence interval) | 1.130 (1.059-1.201) | 0.940 (0.9221-0.958) | 0.977 (0.905-1.050) |
| Proportionality constant $\alpha$ | 1.33 ± 0.78 | 1.00 ± 0.03 | 1.17 ± 0.27 |

**Table S8:** SN DA neuron numbers determined via unbiased stereology and via DLAP-3 on DAB-stained sections. Data and statistics for graphs in Fig. 5b. Given are the numbers of all TH-positive SN neurons, unilaterally determined for each mouse by stereology or algorithm. N represents the number of analysed mice. P-values according to Mann-Whitney test (significant values in bold).

|  |  | manual (Stereology) |  |  |  | automated |  |  |  |
| --- | --- | --- | --- | --- | --- | --- | --- | --- | --- |
| | N | median | 10-90 % | mean | ±SD | median | 10-90 % | mean | ±SD |
|  | 10 | 4789 | 3561-6444 | 4951 | 1079 | 4199 | 3100-6297 | 4317 | 911.8 |
| p-value | Mann-Whitney test: 0.2176 |  |  |  |  |  |  |  |  |

**Table S9:**
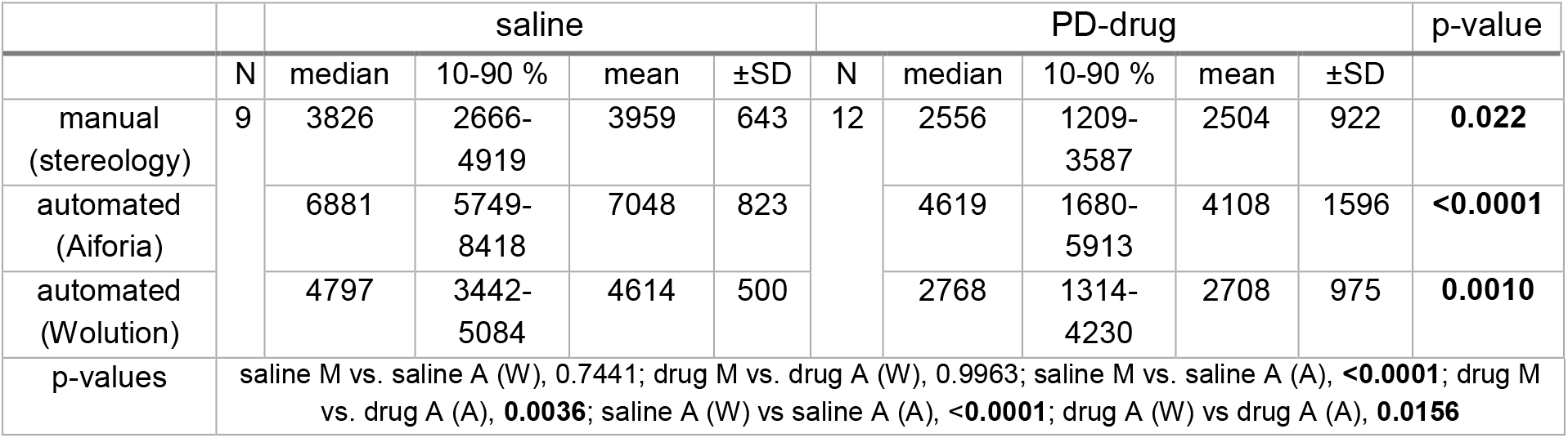
SN DA neuron numbers determined via unbiased stereology and via DLAP-4 on DAB- and hematoxylin-stained sections. Data and statistics for graphs in Fig. 6b. Given are the numbers of all TH-positive SN neurons/mouse, unilaterally determined by stereology or algorithm. N represents the number of analysed mice. P-values according to Two-way ANOVA followed by Tukey’s multiple comparison (significant values in bold).

**Table S10:** Remaining SN DA neurons after neurodegenerative drug (MPTP) treatment, determined via unbiased stereology and via DLAP-4. Data and statistics for graphs in Fig. 6b. Given are the percentages of remaining TH-positive SN neurons/mouse after drug treatment relative to saline, unilaterally determined by stereology or algorithm. N represents the number of analysed mice. P-values according to Kruskal-Wallis with Dunn’s multiple comparison (no significant differences).

| group | N | median | 10-90 % | mean | ±SD |
| --- | --- | --- | --- | --- | --- |
| manual (Stereology) | 12 | 64.6 | 30.5-90.6 | 63.3 | 23.3 |
| automated (Aiforia) |  | 65.5 | 23.8-83.9 | 58.3 | 22.7 |
| automated (Wolution) |  | 60.0 | 28.5-91.7 | 58.7 | 21.1 |
| p-values | M vs A (W), >0.9999; M vs A (A), >0.9999, A (W) vs A (A), >0.9999 |  |  |  |  |

**Table S11:**
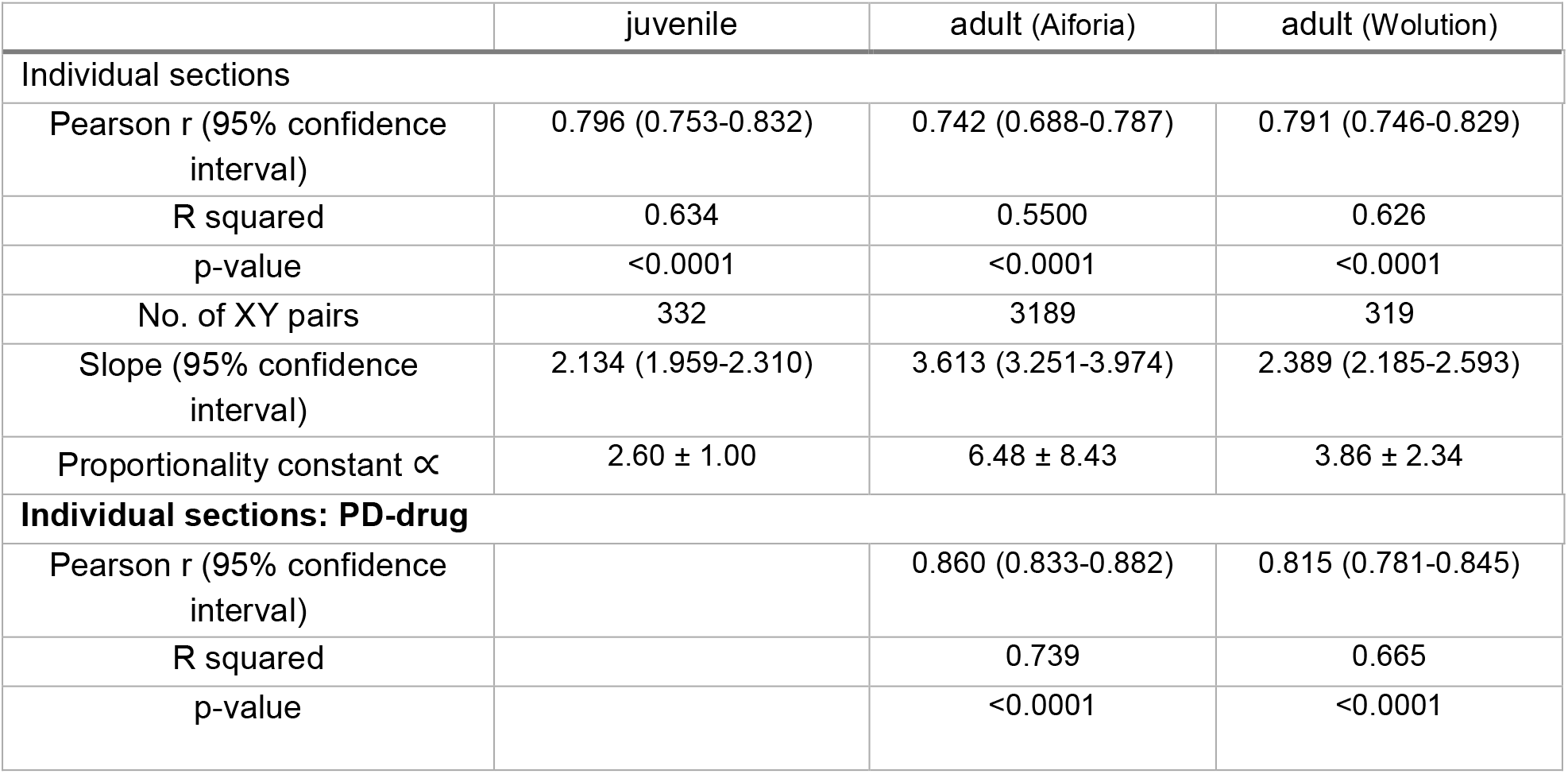
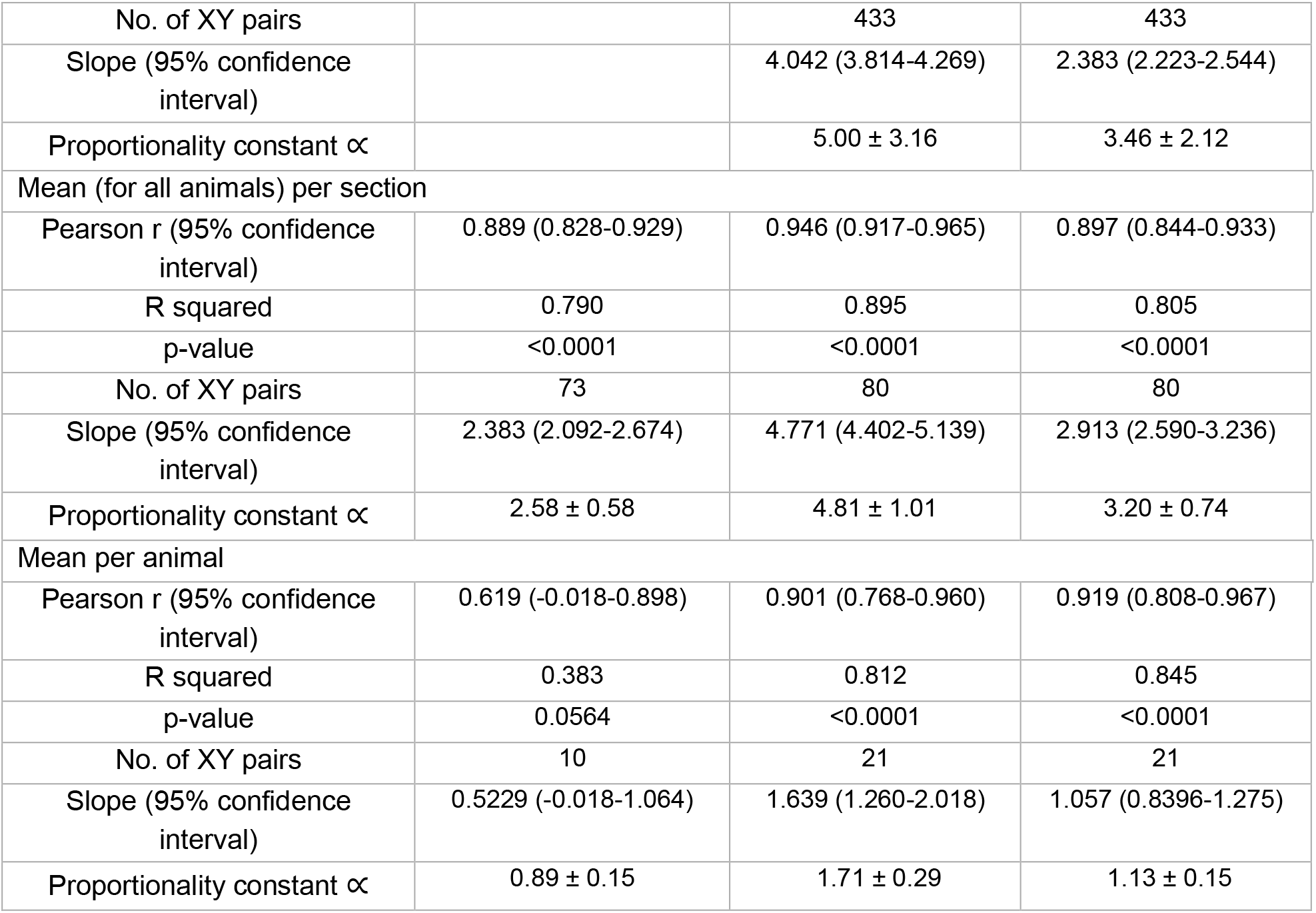
Single cell correlation analyses between manually and DLAP-3 and DLAP-4 determined results. Data and statistics for graphs in Fig. 5c/6c, according to Pearson correlation (slope of the linear regression with 95% confidence interval).

**Table S12:** Number of TH-positive SN neurons analysed via DLAP-5. Data and statistics for graphs in Fig. 7b, 8b and S5c. Given are the numbers of all TH-positive SN neurons/mouse, bilaterally determined by the algorithm, the number of neurons excluded/included neurons for further DAT analysis (criteria specified in Figure S5b), and the determined number of TH positive DAT positive/negative SN neurons. Rostral, medial and caudal locations were determined according to the respective stereotaxic coordinates (bregma; rostral: -2.7 to -3.1 mm; medial: -3.1 to -3.5; caudal: -3.5 to -3.9). Lateral and non-lateral SN-areas were defined as detailed in Figure S6. N represents the number of analysed mice. P-values according to Wilcoxon or Mann-Whitney tests (significant values in bold).

| | N | mean | ±SD | % | mean | ±SD | % | mean | ±SD | % |
| --- | --- | --- | --- | --- | --- | --- | --- | --- | --- | --- |
|  |  | all detected TH+ SN neurons |  |  | excluded TH+ neurons from further DAT analysis |  |  | included TH+ neurons for further DAT analysis |  |  |
| whole SN | 14 | 3902 | 507 | 100 | 1151 | 220 | 30 ± 6 | 2750 | 462 | 70 ± 6 |
| p-value |  | MWU: all vs analysed TH+ neurons **<0.0001** |  |  |  |  |  |  |  |  |
|  |  | identified TH+ DAT+ |  |  | identified TH+ DAT- |  |  | all included TH+ neurons |  |  |
| whole SN | 14 | 2633 | 442 | 96 ± 2 | 118 | 50 | 4 ± 2 | 2750 | 462 | 100 |
| rostral | | 876 | 176 | 99 ± 1 | 10 | 8 | 1 ± 1 | 886 | 178 | 100 |
| medial | | 987 | 248 | 95 ± 2 | 53 | 24 | 5 ± 2 | 1040 | 260 | 100 |
| caudal | | 767 | 126 | 93 ± 3 | 57 | 25 | 7 ± 3 | 824 | 132 | 100 |
| p-value |  | Wilcoxon: all analysed TH+ vs analysed TH+ DAT+ **0.0001**<br>Chi-square: rostral vs medial **<0.0001**, rostral vs caudal **<0.0001**, caudal vs medial **<0.0001** |  |  |  |  |  |  |  |  |
| **lateral SN** |  | identified TH+ DAT+ |  |  | identified TH+ DAT- |  |  | all included TH+ neurons |  |  |
| whole axis | 14 | 194 | 27 | 83 ± 6 | 41 | 19 | 17 ± 6 | 235 | 41 | 100 |
| rostral | | 56 | 19 | 95 ± 5 | 4 | 5 | 5 ± 5 | 60 | 22 | 100 |
| medial | | 104 | 26 | 85 ± 5 | 19 | 9 | 15 ± 5 | 123 | 32 | 100 |
| caudal | | 34 | 17 | 63 ± 15 | 19 | 9 | 37 ± 15 | 53 | 20 | 100 |
| p-value |  | Wilcoxon: all analysed TH+ vs analysed TH+ DAT+ **0.0001**<br>Chi-square: rostral vs medial 0.1026, rostral vs caudal **0.0001**, caudal vs medial **0.0025** |  |  |  |  |  |  |  |  |
| **non-lateral SN** |  | identified TH+ DAT+ |  |  | identified TH+ DAT- |  |  | all included TH+ neurons |  |  |
| whole axis | 14 | 2436 | 405 | 97 ± 2 | 79 | 37 | 3 ± 2 | 2515 | 410 | 100 |
| rostral |  | 820 | 170 | 99 ± 1 | 7 | 5 | 1 ± 1 | 827 | 170 | 100 |
| medial |  | 883 | 219 | 96 ± 2 | 34 | 16 | 4 ± 2 | 917 | 223 | 100 |
| caudal |  | 733 | 118 | 95 ± 3 | 38 | 22 | 5 ± 3 | 771 | 122 | 100 |
| p-value |  | Wilcoxon: all analysed TH+ vs analysed TH+ DAT+ **0.0001**<br>Chi-square: rostral vs medial **<0.0001**, rostral vs caudal **<0.0001**, caudal vs medial 0.2163 |  |  |  |  |  |  |  |  |

**Table S13:**
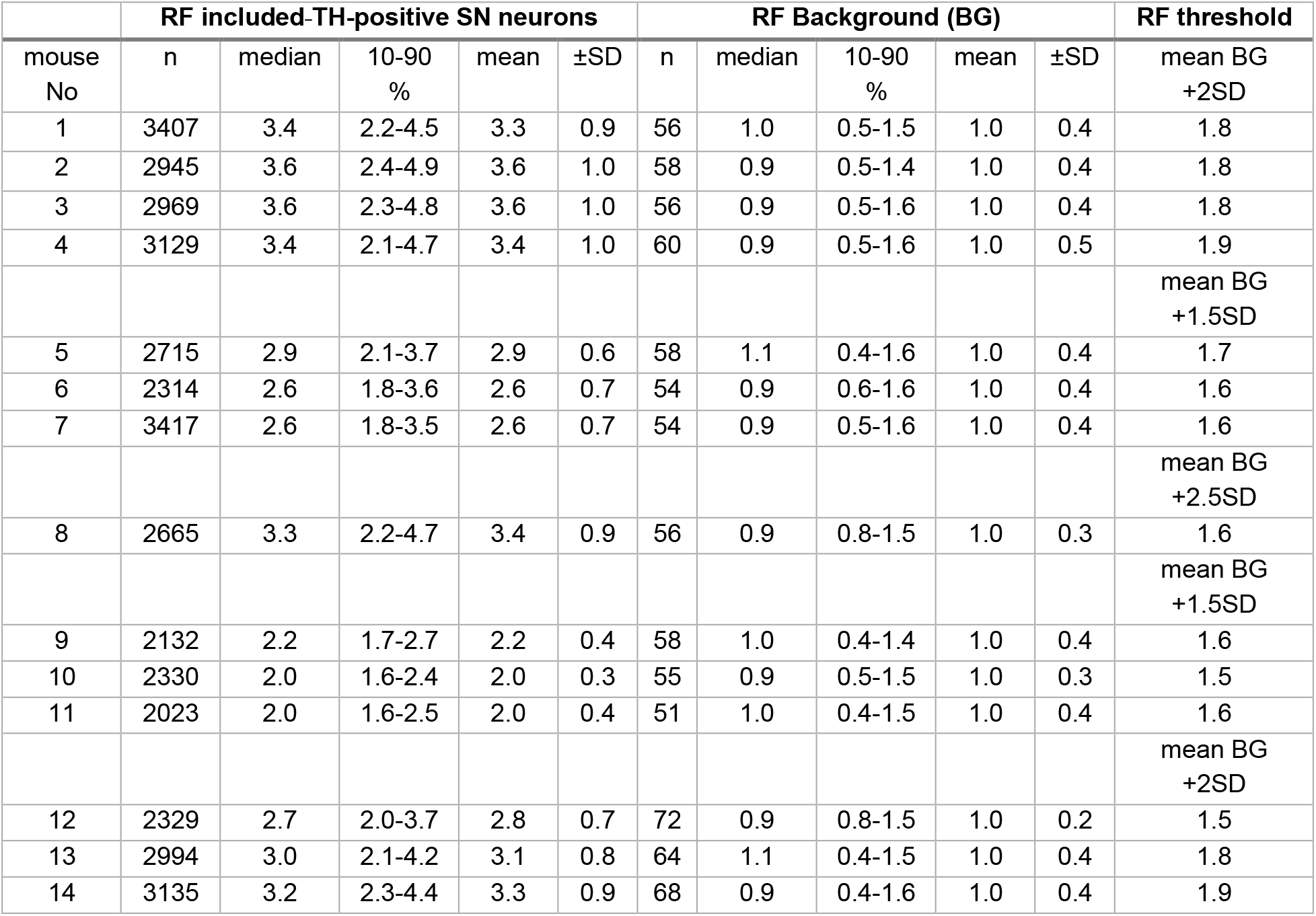
Relative DAT immunofluorescence signal intensities (RF) for included TH-positive SN neurons, for each mouse, determined via DLAP-5. Data and statistics for graphs in Fig. S5d, and S7 (background normalised). n represents the number of total analysed SN DA neurons or the number background (BG) images. TH-positive neurons with DAT-signals over the respective “threshold”-values were defined as DAT-positive.

**Table S14:**
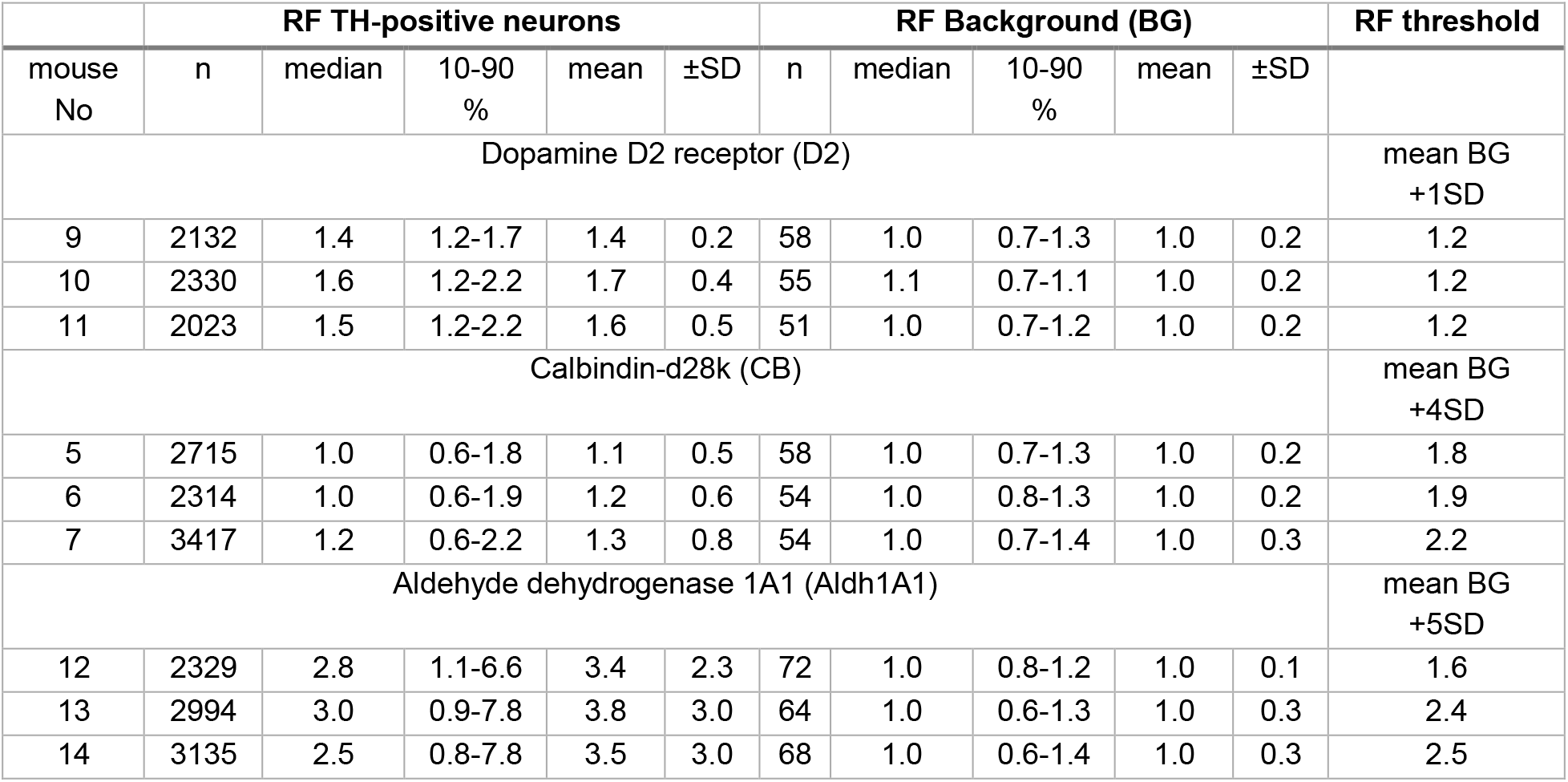
Relative D2, CB and Aldh1A1 immunofluorescence signal intensities (RF) in TH-positive SN neurons for each mouse, determined via DLAP-5. Data and statistics for graphs in Fig. 11 and S8 (background normalised). n represents the number of total analysed TH-positive neurons or the number background (BG) images, as indicated. TH-positive neurons with D2/CB/Adh1A1-signals over the respective “threshold”-values were define as immono-positive for the respective protein.

**Table S15:** Relative D2, CB and Aldh1A1 immunofluorescence signal intensities (RF) in DAT-positive and DAT-negative TH-positive SN neurons, determined via DLAP-5. Data and statistics for graphs in Fig. 10 (background normalised). Further processed data from table S14. n represents the number of total analysed TH-positive neurons. P-values according to Mann-Whitney tests (significant values in bold).

| marker gene | TH-positive SN neuron subtype | DAT-positive SN neuron subtype |  |  | DAT-negative SN neuron subtype |  |  | p-value |
| --- | --- | --- | --- | --- | --- | --- | --- | --- |
|  |  | n | Mean RF | ±SD | n | Mean RF | ±SD |  |
| **Dopamine D2-autoreceptor (D2)** | all TH-positives | 6108 | 1.58 | 0.41 | 377 | 1.41 | 0.49 | **<0.0001** |
|  | D2-positive | 5663 | 1.62 | 0.41 | 255 | 1.56 | 0.53 | **0.0035** |
|  | D2-negative | 445 | 1.13 | 0.05 | 122 | 1.09 | 0.07 | **<0.0001** |
| **Calbindin-d28k (CB)** | all TH-positives | 8063 | 1.18 | 0.64 | 383 | 1.91 | 1.02 | **<0.0001** |
|  | CB-positive | 722 | 2.63 | 0.89 | 143 | 2.96 | 0.83 | **<0.0001** |
|  | CB-negative | 7341 | 1.03 | 0.39 | 240 | 1.28 | 0.42 | **<0.0001** |
| **Aldehyde dehydrogenase (Aldh1A1)** | all TH-positive | 8206 | 3.65 | 2.80 | 252 | 1.02 | 0.74 | **<0.0001** |
|  | Aldh1A1-positive | 4810 | 5.31 | 2.57 | 14 | 3.55 | 0.80 | **<0.0001** |
|  | Aldh1A1-negative | 3396 | 1.30 | 0.47 | 238 | 0.87 | 0.37 | **<0.0001** |

**Table S16:**
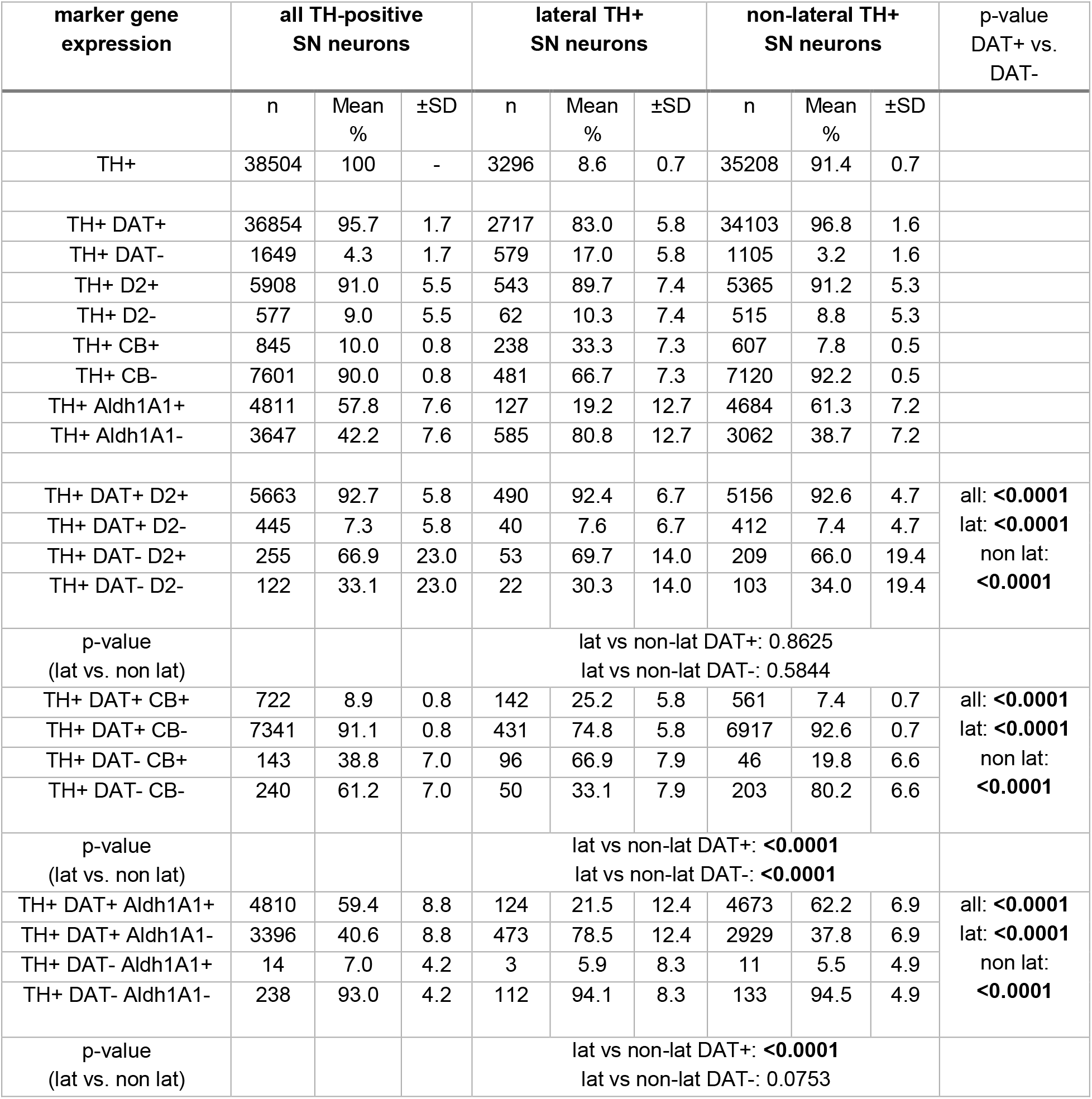
TH-positive SN neuron marker gene co-expression analysis. Data and statistics for graphs in Fig. 7b/d and 11. The total numbers of marker gene-positive and -negative neurons is represented by n. The relative % of marker gene and TH co-expression are given for all analysed TH-positive (TH+) SN neurons as well as separately for lateral (lat) and non-lateral (non-lat) TH+ neurons. P-values according to Chi-square test (significant values in bold).

**Table S17:** Soma size of all analysed TH-positive SN neurons, determined via DLAP-5. Data for graphs in Fig. 7e. Cell body sizes [µm^2^] were estimated according to the TH-positive area that was marked by DLAP-5, for all analysed TH-positive neurons, and are given for different SN-neuron subtypes, defined according to their co-expression profile.

| marker gene<br>co-expression type | Number of<br>analysed<br>mice | all TH+ SN neurons<br>cell body area [$\mu\text{m}^2$] | | lateral TH+<br>SN neurons<br>cell body area [$\mu\text{m}^2$] | | non-lateral TH+<br>SN neurons<br>cell body area [$\mu\text{m}^2$] | |
| --- | --- | --- | --- | --- | --- | --- | --- |
| | N | Mean | ±SD | Mean | ±SD | mean | ±SD |
| all TH+ DAT+ | 14 | 195.5 | 14.4 | 186.4 | 16.6 | 196.2 | 14.8 |
| all TH+ DAT- | 14 | 131.0 | 16.5 | 117.9 | 19.1 | 138.2 | 16.6 |
| caudal TH+ DAT+ | 14 | 194.5 | 16.2 | 185.1 | 23.0 | 195.5 | 16.3 |
| caudal TH+ DAT- | 14 | 127.7 | 22.2 | 115.5 | 19.8 | 142.5 | 24.2 |
| medial TH+ DAT+ | 14 | 189.5 | 15.3 | 197.0 | 18.8 | 187.4 | 16.0 |
| medial TH+ DAT- | 14 | 131.7 | 15.9 | 120.9 | 17.5 | 140.0 | 16.3 |
| rostral TH+ DAT+ | 14 | 202.7 | 14.9 | 210.1 | 14.3 | 200.9 | 15.7 |
| rostral TH+ DAT- | 14 | 150.2 | 36.0 | 124.2 | 33.3 | 167.9 | 39.8 |
| TH+ D2+ | 3 | 183.9 | 4.5 | 192.4 | 4.6 | 182.1 | 5.2 |
| TH+ D2- | 3 | 167.7 | 7.8 | 161.5 | 2.7 | 168.7 | 9.0 |
| TH+ CB+ | 3 | 161.2 | 9.7 | 133.2 | 1.8 | 174.5 | 9.8 |
| TH+ CB- | 3 | 193.4 | 8.1 | 199.6 | 5.7 | 192.1 | 8.6 |
| TH+ Aldh1A1+ | 3 | 176.0 | 5.6 | 189.6 | 3.0 | 174.8 | 6.3 |
| TH+ Aldh1A1- | 3 | 183.1 | 3.5 | 163.4 | 3.3 | 193.1 | 5.4 |
| TH+ DAT+ D2+ | 3 | 185.9 | 3.8 | 198.7 | 6.2 | 183.3 | 4.9 |
| TH+ DAT+ D2- | 3 | 174.6 | 5.5 | 187.0 | 11.7 | 173.2 | 8.0 |
| TH+ DAT- D2+ | 3 | 140.8 | 9.0 | 122.1 | 7.9 | 149.5 | 8.3 |
| TH+ DAT- D2- | 3 | 145.5 | 8.7 | 133.8 | 13.5 | 148.6 | 11.2 |
| TH+ DAT+ CB+ | 3 | 172.7 | 6.6 | 158.2 | 2.6 | 176.8 | 7.3 |
| TH+ DAT+ CB- | 3 | 195.0 | 7.9 | 203.5 | 5.3 | 193.2 | 8.4 |
| TH+ DAT- CB+ | 3 | 104.2 | 7.6 | 96.0 | 4.6 | 141.6 | 28.1 |
| TH+ DAT- CB- | 3 | 145.3 | 10.6 | 132.0 | 8.8 | 151.3 | 13.8 |
| TH+ DAT+<br>Aldh1A1+ | 3 | 176.1 | 5.5 | 189.3 | 3.1 | 175.0 | 6.2 |
| TH+ DAT+ Aldh1A1- | 3 | 187.6 | 5.0 | 171.8 | 5.2 | 194.8 | 5.9 |
| TH+ DAT- Aldh1A1+ | 3 | 132.6 | 27.3 | 263.9 | 0.0 | 126.3 | 28.5 |
| TH+ DAT- Aldh1A1- | 3 | 112.9 | 11.7 | 101.9 | 11.3 | 135.0 | 12.9 |

**Table S18:**
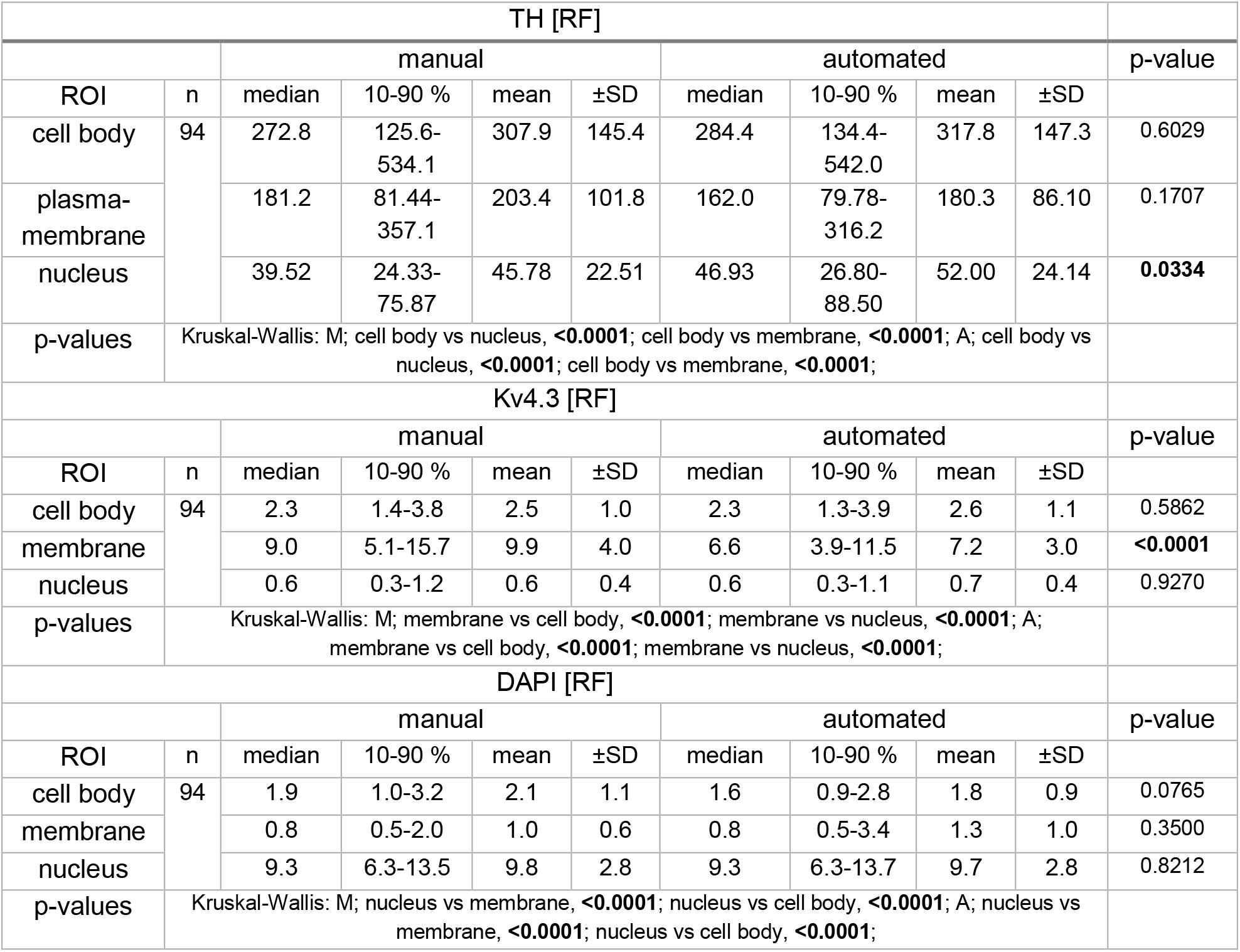
Relative immunofluorescence signal intensities (RF) in SN DA neuron cellular compartments, determined manually and via DLAP-6. Data and statistics for graphs in Fig. 12b and S9 (background normalised). Number of total analysed neurons is given by n, N represents the number of analysed mice. P-values according to Kruskal-Wallis with Dunn’s multiple comparison and Mann-Whitney test (significant values in bold).

**Table S19:** Single cell correlation analyses between manually and DLAP-6 determined results. Data and statistics for graphs in Fig. 12c., according to Pearson correlation (slope of the linear regression with 95% confidence interval).

|  | Cytoplasm:<br>TH | Plasma-membrane:<br>Kv4.3 | Nucleus:<br>DAPI |
| --- | --- | --- | --- |
| Pearson r (95% confidence interval) | 0.995 (0.992-0.997) | 0.966 (0.949-0.977) | 0.996 (0.994-0.008) |
| R<sup>2</sup> | 0.990 | 0.933 | 0.993 |
| p-value | <0.0001 | <0.0001 | <0.0001 |
| No. of XY pairs | 94 | 94 | 94 |
| Slope (95% confidence interval) | 1.01 (0.989-1.029) | 0.737 (0.696-0.777) | 0.986 (0.969-1.00) |
| Proportionality constant $\alpha$ | 0.96 ± 0.05 | 1.38 ± 0.15 | 1.01 ± 0.03 |

**Table S20:**
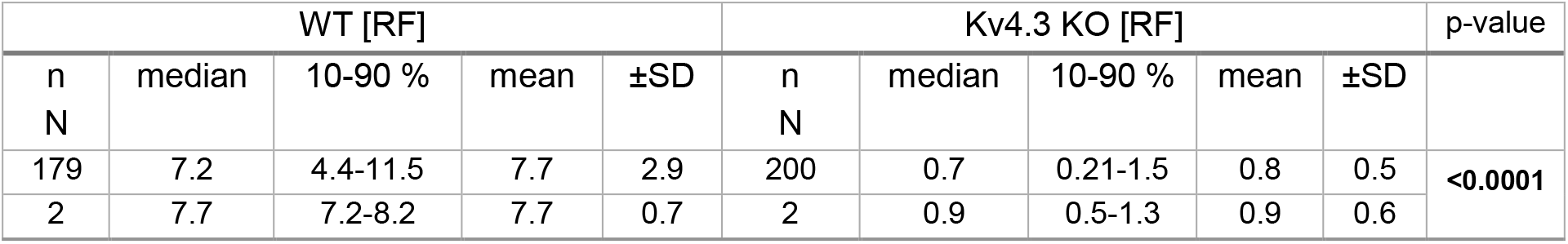
Relative immunofluorescence-signal quantification for Kv4.3 in SN DA neuron membranes from WT and Kv4.3 KO mice, determined via DLAP-6. Data and statistics for graphs in Fig. 12e and S9 (background normalised). Number of analysed neurons is given by n, N represents the number of analysed mice. P-values according Mann-Whitney test (significant values in bold).

